# Human-specific features and developmental dynamics of the brain N-glycome

**DOI:** 10.1101/2023.01.11.523525

**Authors:** Thomas S. Klarić, Ivan Gudelj, Gabriel Santpere, André M. M. Sousa, Mislav Novokmet, Frano Vučković, Shaojie Ma, Ivona Bečeheli, Chet C. Sherwood, John J. Ely, Patrick R. Hof, Djuro Josić, Gordan Lauc, Nenad Sestan

## Abstract

Comparative “omics” studies have revealed unique aspects of human neurobiology, yet an evolutionary perspective of the brain N-glycome is lacking. Here, we performed multi-regional characterization of rat, macaque, chimpanzee, and human brain N-glycomes using chromatography and mass spectrometry, then integrated these data with complementary glycotranscriptomic data. We found that in primates the brain N-glycome has evolved more rapidly than the underlying transcriptomic framework, providing a mechanism for generating additional diversity. We show that brain N-glycome evolution in hominids has been characterized by an increase in complexity and α(2-6)-linked N-acetylneuraminic acid along with human-specific cell-type expression of key glycogenes. Finally, by comparing the prenatal and adult human brain N-glycome, we identify region-specific neurodevelopmental pathways that lead to distinct spatial N-glycosylation profiles in the mature brain.

**One-Sentence Summary:** Evolution of the human brain N-glycome has been marked by an increase in complexity and a shift in sialic acid linkage.

## Main Text

The human brain is an immensely complex organ and accordingly must undergo a protracted period of development before reaching maturity (*1*). There is an increasing desire to uncover the cellular and molecular evolutionary specializations that make the human brain, and its development, unique. The majority of our knowledge regarding such human-specific features has been derived from comparative “omics” studies involving humans and non-human primates at the genome, transcriptome, proteome and metabolome level (*2*–*8*). However, the contribution of post-translational sugar modifications in generating inter-species molecular diversity has not been explored in the context of primate brain evolution.

Glycans mediate many important biological phenomena that occur at the cell surface, including functions critical to the host organism and interactions with pathogens. Consequently, the evolution of the glycome is uniquely driven by both intrinsic and extrinsic selection pressures (*9*). Asparagine-linked glycosylation (N-glycosylation) is critically involved in many aspects of central nervous system (CNS) biology including membrane excitability, synaptic transmission, neurite outgrowth, and neuronal plasticity (*10*, *11*). Recent insights have revealed that the brain N-glycome is dynamic and that its composition changes under various physiological and pathological conditions (*12*), yet data regarding its evolutionary trajectory are scarce. Only a single comparative N-glycomics study has investigated inter-species differences in N-glycosylation at the whole N-glycome level though it was limited to two species, mice and humans, and a single brain region (*13*). Moreover, no data exist regarding the neuroglycome of non-human primates. Consequently, we lack a holistic view of the evolutionary trends that have shaped the neuroglycome in primates.

Here, we performed a multi-regional comparative N-glycomics study to define, and anatomically pinpoint, evolutionary specializations unique to primates with an emphasis on identifying human-specific features of brain N-glycosylation. Four species representing key phylogenetic groups within the Euarchontoglires branch of mammals were investigated: humans, chimpanzees, rhesus macaque monkeys, and rats. In each species, four functionally and cytoarchitecturally distinct brain regions (or their corresponding homologous equivalents) were investigated: the dorsolateral prefrontal cortex (dlPFC or DFC), hippocampus (HIP), striatum (STR), and cerebellar cortex (CBC). We performed detailed structural characterization of the brain N-glycomes using an ultra-performance liquid chromatography method based on hydrophilic interactions (HILIC-UPLC) coupled with matrix assisted laser desorption/ionization time of flight/time of flight tandem mass spectrometry (MALDI-TOF/TOF-MS/MS) (Fig S1-S2). In addition, we then integrated these N-glycomics data with complementary tissue and single-nucleus level RNA sequencing (RNA-seq) data to explore the relationship between glycogene expression and N-glycosylation in the context of primate brain evolution. Finally, by comprehensively characterizing the spatiotemporal dynamics of N-glycosylation during human brain development, we shed light on the neurodevelopmental trends that govern the maturation of the human brain N-glycome.

### Overview of the Euarchontoglires brain N-glycome

Total N-glycans were isolated from brain tissue (Fig S3, Table S1-S2), labelled with 2-aminobenzamide (2-AB), and separated chromatographically as previously described (*14*), yielding a complex N-glycoprofile (Fig 1A). MALDI-TOF/TOF-MS/MS analysis revealed 145 distinct monosaccharide compositions which account for a total of 287 distinct N-glycan structures across all adult brain regions and species (Table S3), including those with distinctive structural features typical of brain N-glycans (Supplementary text, Figs S4-S6). Strikingly, only 60 N-glycans (Table S4) were ubiquitously present and their combined abundance disproportionately accounts for approximately 75% of the entire brain N-glycome. This conserved ‘core network’ (Fig 1B) predominantly consists of oligomannose and neutral fucosylated hybrid/complex structures supporting the assertion that although there is a tremendous amount of structural diversity among brain N-glycans (*12*), a relatively small number of simple N-glycans comprise the bulk of the brain N-glycome (*15*).

**Fig. 1.**
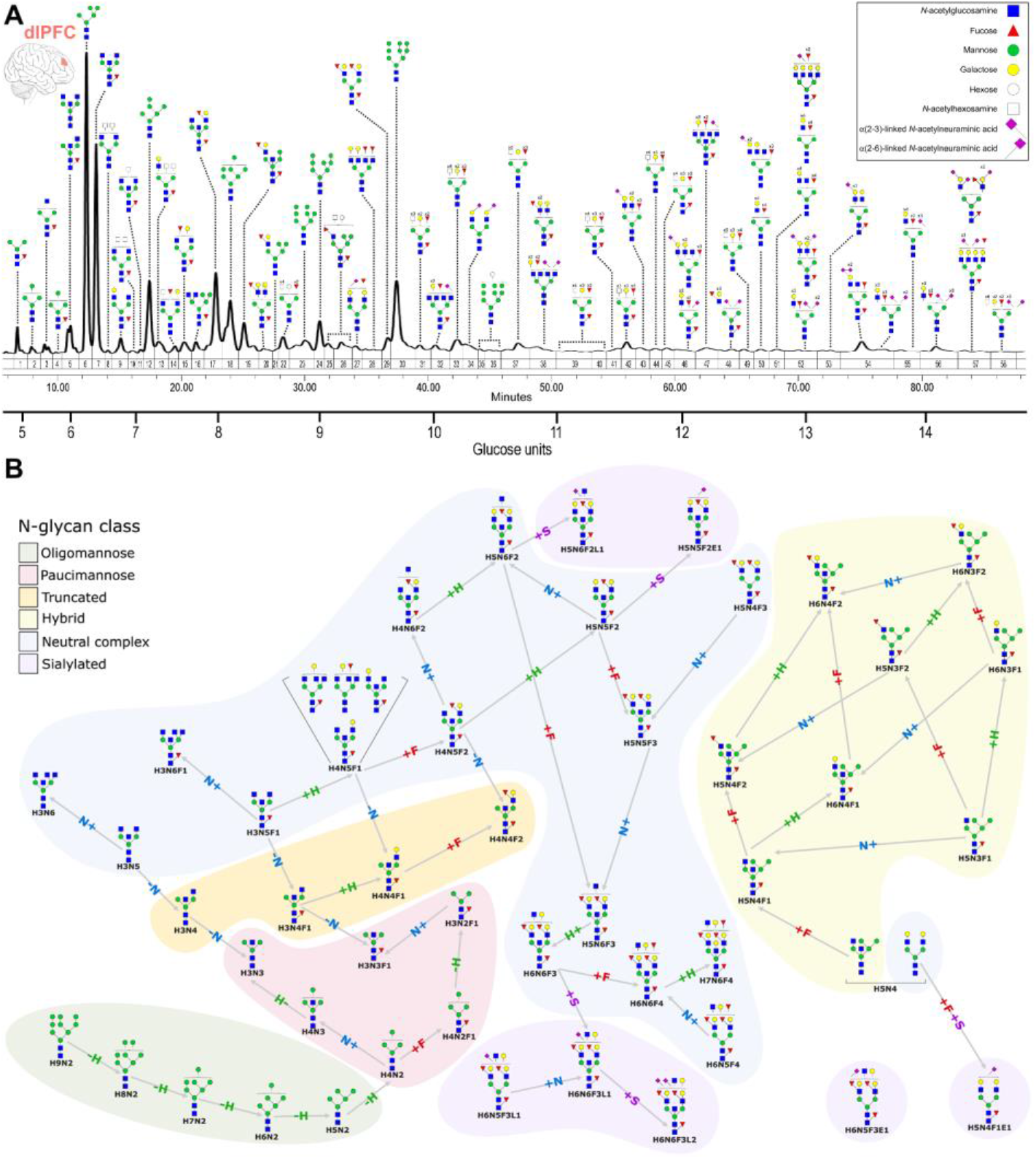
Organization of the Euarchontoglires brain N-glycome. (**A**) Representative chromatogram of N-glycans derived from the human dlPFC annotated with proposed structures. Only the major N-glycan species in each peak are shown (see Table S3 for complete list). Chromatographic peaks are labelled (1-58) and integration boundaries are shown as vertical lines. The x-axis displays analyte retention time, while the y-axis shows fluorescence at 428 nm. (**B**) Overview of the conserved brain N-glycome ‘core network’ based on the 60 ubiquitous N-glycans that were detected in all brain regions of all species. The network was constructed using with the assumption that the core N-glycan structures are biosynthetically related. H, hexose; N, N-acetylhexosamine; F, deoxyhexose; S, Neu5Ac; L, α(2-3)-linked Neu5Ac (lactonized); E, α(2-6)-linked Neu5Ac (esterified).

For each group, we quantified the abundance of individual N-glycan peaks (Figs S7-S8, Table S5-S6) and, in turn, the resulting derived N-glycan traits (Figs S9-S11, Table S7-8), which provided us with an overview of the composition of all the N-glycomes (Fig S12). It emerges that, quantitatively and qualitatively, the various brain N-glycoprofiles are generally similar and conform to the same overall ‘brain N-glycosylation template’ characterized by an abundance of oligomannose (33.7-44.0%) and hybrid structures (5.9-11.5%), with the remainder being complex N-glycans (41.0-53.7%) and a minor proportion of truncated (<6%) and paucimannose (<3%) N-glycans. Despite their overall similarity, careful examination of the differences between N-glycomes revealed a wealth of anatomical and phylogenetic variation in both the abundance and content of individual peaks (Supplementary text, Fig S13, Table S5-6), which is ultimately reflected in statistically significant differences in derived N-glycan traits (Table S8).

### The cerebellum has a distinctive and well conserved N-glycome

To explore the relationship between the various brain N-glycomes at a global level, we analyzed N-glycan peak abundance data using hierarchical clustering and t-distributed stochastic neighbor embedding (t-SNE). These analyses showed that the CBC exhibits the most distinctive N-glycome to the extent that it even overrides large phylogenetic distances (Fig 2A-B). This observation mirrors the results of similar analyses of the brain transcriptome and proteome (*16*, *17*) and suggests that the cerebellar N-glycome diverged from the remaining brain regions early and has remained well conserved throughout evolution. The defining feature of the cerebellar N-glycome is an increased abundance of the M6 N-glycan. (Fig 2C, S7, S11). Nonetheless, phylogenetic divergence within the CBC cluster was also apparent and can be attributed to lineage-specific alterations in various N-glycan traits (Fig S14). Subsequent clustering of brain N-glycomes was much more dependent on phylogeny than regional differences indicating that substantial species-specific differences in N-glycosylation are a feature of the remaining brain regions.

**Fig. 2.**
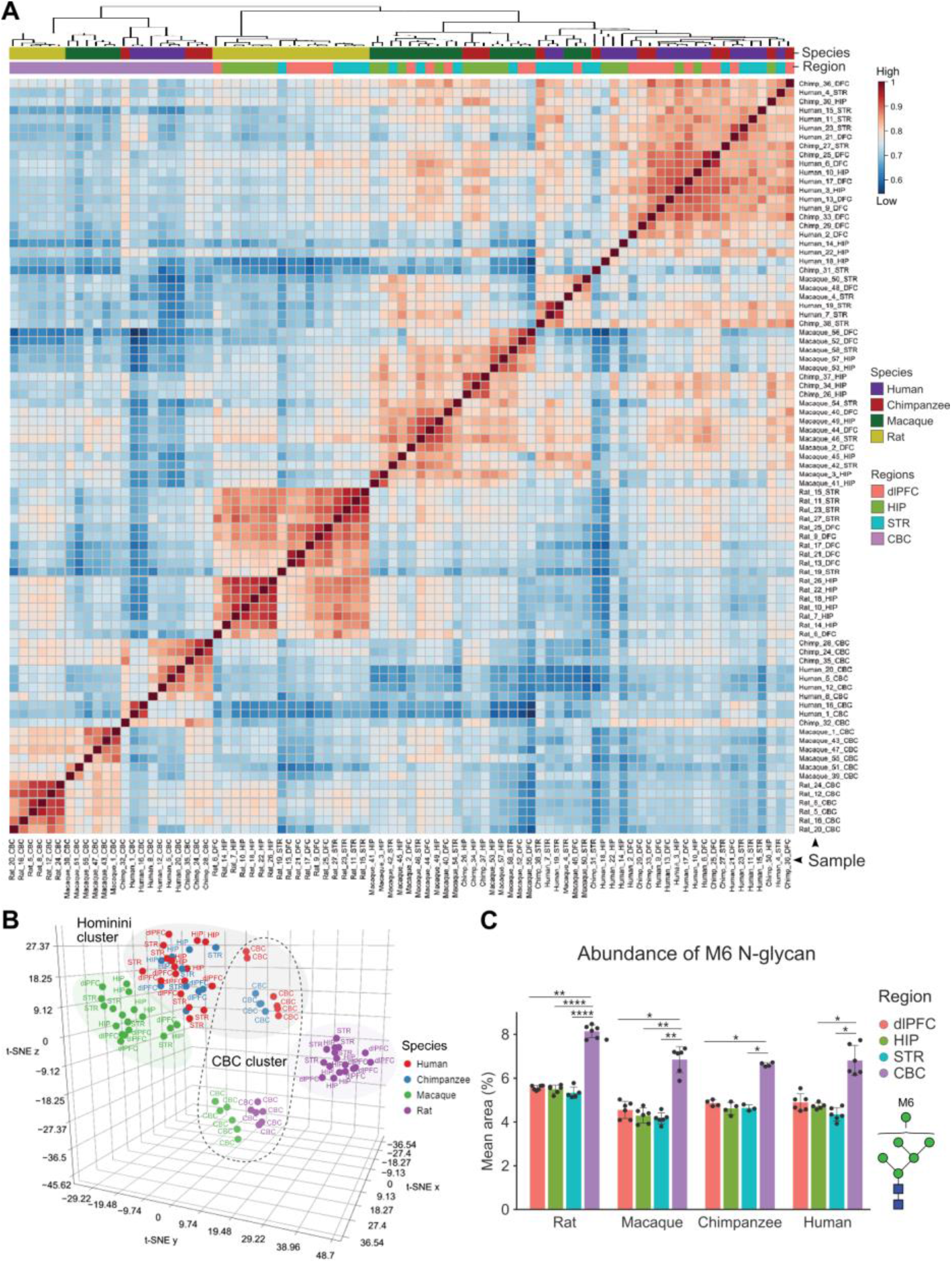
The brain N-glycome is characterized by both anatomical and phylogenetic variation. The correlations between the various mammalian brain N-glycomes were assessed using their UPLC-derived N-glycoprofiles (i.e. the relative abundance of corresponding N-glycan peaks). (**A**) Kendall rank correlation heatmap and dendrograms illustrating the degree of correlation among the different brain N-glycomes. The color key indicates the strength of the correlation. (**B**) Three-dimensional tSNE scatter plot illustrating the degree of relatedness among the different brain N-glycomes. Note that the CBC N-glycomes from all species form a distinct cluster separate from the other brain regions and that the hominini CBC N-glycomes are separated from those of macaques and rats. Axes represent the t-SNE x, y, and z dimensions. (**C**) In all species, the CBC N-glycome is characterized by an increased abundance of the M6 oligomannose N-glycan found in chromatographic peak 12. Statistical significance between groups was tested using ANCOVA and correction for multiple testing was performed using the Bonferroni procedure. Adjusted P-values: * 0.05>P>0.01, ** 0.01>P>0.001, *** 0.001>P>0.0001, **** P<0.0001.

### Intra-species anatomical variation in brain N-glycosylation

Although the general features of ‘brain-type’ N-glycosylation are conserved, the composition of the N-glycome varies across different brain regions (Fig S7, S9, S11-S13, Table S3). Generally, the same nominally significant anatomical trends were consistently observed in all species though they more frequently survived correction for multiple testing in rats, likely due to their lower inter-individual variability (Fig S3). The most striking trend was a gradient of increasing abundance of complex N-glycans from the CBC, through the STR, to the cortical structures (Supplementary text, Fig S15). This was accompanied by a complementary increase in features associated with complex N-glycans (e.g. galactosylation, fucosylation, and branching) which again is most evident in rats (Fig S9). While analysis of larger and more homogeneous primate cohorts is needed to confirm whether these neuroanatomical patterns of N-glycosylation are conserved, our data suggest that brain regions that have undergone significant transformation over the course of mammalian evolution have a greater proportion of complex N-glycans compared to more conserved brain regions.

### Phylogenetic trends and human-specific features of the brain N-glycome

Comparing the N-glycosylation profiles of equivalent homologous regions across species allowed us to gain insights into how the neuroglycome has changed over the course of Euarchontoglires evolution (Supplementary text, Fig S10-11B). Firstly, we observed a marked linear trend in the relative abundance of complex N-glycans which, in all brain regions, was lowest in rats, slightly higher in macaques, and greatest in the hominid species (Fig 3A-D, Fig S10). The rates of change were consistent with the evolutionary distance between species indicating a gradual and uniform change over time (Fig S16). Interestingly, this phylogenetic gradient in brain N-glycome complexity parallels the gradient observed in a comparative brain lipidome study (*18*). Secondly, we noted a striking phylogenetic trend towards increased usage of α(2-6)-linked Neu5Ac moving from rats, through macaques, to the hominids which manifested as a decrease in the proportion of sialylated complex N-glycans homogeneously adorned with α(2-3)-linked Neu5Ac coupled with a corresponding increase in those carrying exclusively α(2-6)-linked Neu5Ac or those carrying both α(2-3)- and α(2-6)-linked Neu5Ac. The same overall trend was observed both at the whole brain level using aggregate data (Fig 4E-G) and when individual brain regions were analyzed separately (Fig S17). Detailed analysis of both Neu5Ac density and linkage among sialylated complex N-glycans revealed that the most notable difference occurs in disialylated N-glycans homogeneously adorned with α(2-6)-linked Neu5Ac which are considerably more prevalent in human brains (Fig S18). It also uncovered a striking Neu5Ac usage pattern specific to primates (Supplementary text). Overall, our data indicate that the evolution of the human brain N-glycome was characterized by a brain-wide increase in N-glycome complexity coupled with a shift towards greater usage of α(2-6)-linked Neu5Ac.

**Fig. 3.**
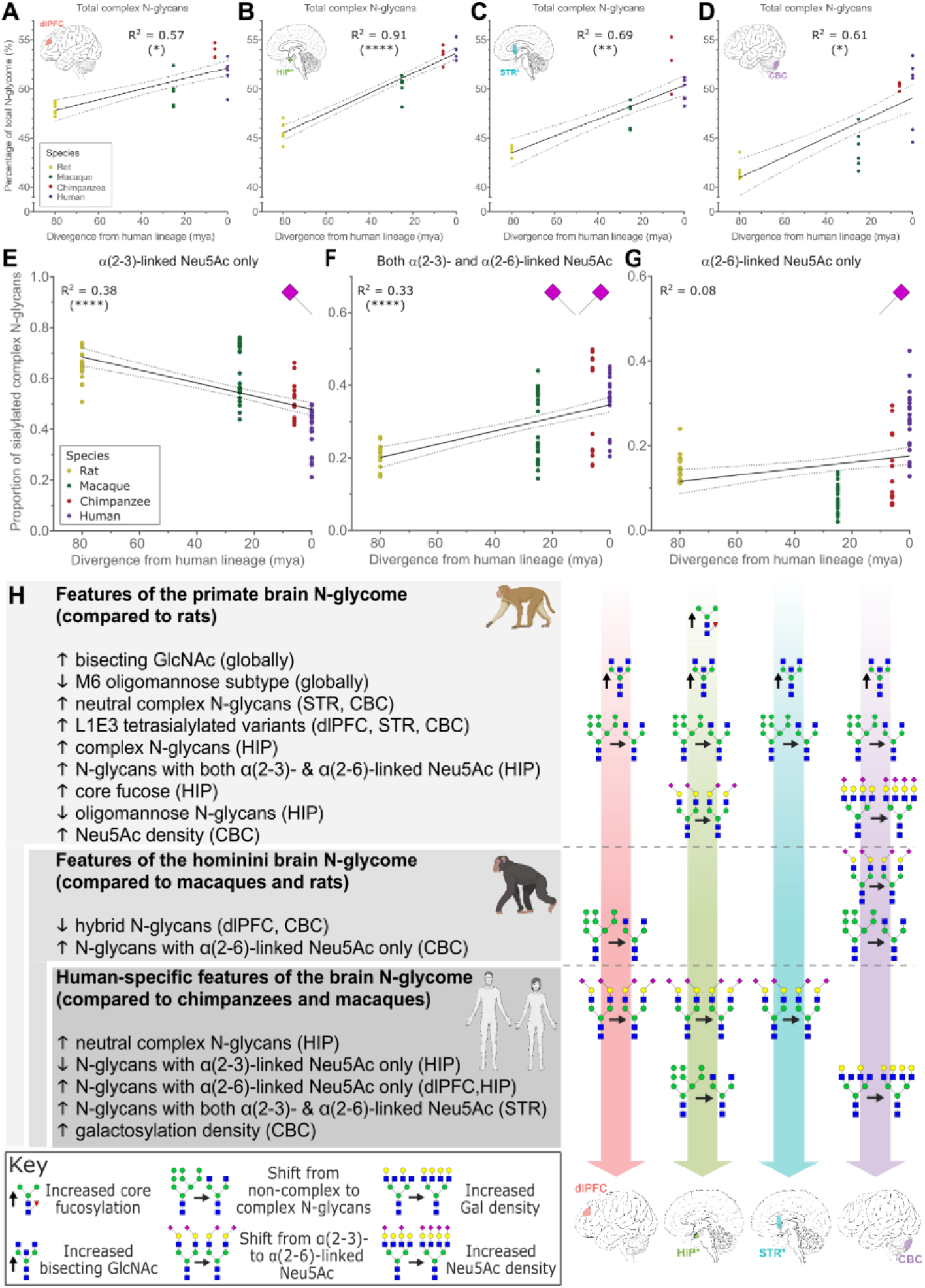
Phylogenetic trends and taxon-specific evolutionary specializations in Euarchontoglires brain N-glycomes. Phylogenetic gradient of complex N-glycan abundance in the (**A**) dlPFC, (**B**) HIP, (**C**) STR, and (**D**) CBC. Data were analyzed using linear regression (90% confidence intervals and R^2^ values are shown). Asterisks in parentheses indicate lines whose slopes were significantly non-zero after correction for multiple testing using the Bonferroni procedure. Adjusted P-values: (*) 0.05>P≥0.01, (**) 0.01>P≥0.001, (****) P<0.0001. mya - millions of years ago. (**E-G**) Phylogenetic trends in Neu5Ac linkage distribution. The plots show the proportion of sialylated complex N-glycans carrying only α(2-3)-linked Neu5Ac (E), both α(2-3)- and α(2-6)-linked Neu5Ac (F), or only α(2-6)-linked Neu5Ac (G). Aggregate (i.e. whole brain) data are plotted and were analyzed as in (A-D). See Fig S17 for individual brain region data. (**H**) Summary of statistically significant taxon-specific differences in N-glycosylation for each brain region (see Supplementary text and Table S8 for details). For clarity, traits that result in an increase in the proportion of complex N-glycans have been symbolically grouped together as have those that result in an increase in the proportion of α(2-6)-linked Neu5Ac. *Denotes that regions are internal and not visible from the view shown.

**Fig. 4.**
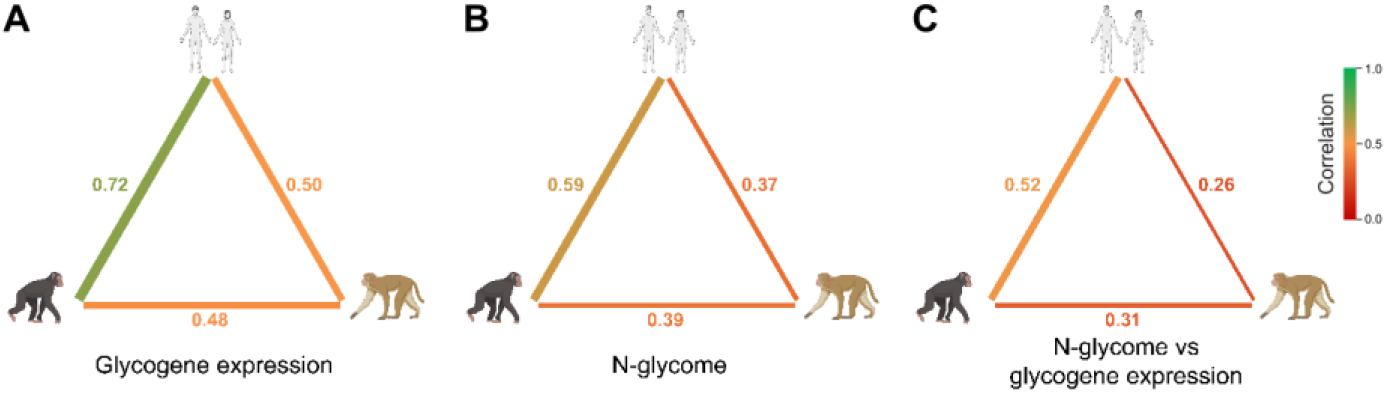
The N-glycomes of primate brains are less conserved than the expression of the underlying glycogenes. The correlation between each primate pair was determined with regard to (**A**) glycogene expression, (**B**) the N-glycome, and (**C**) the correlation between glycogene expression and the N-glycome phenotype. A Mantel test was used for (A) and (B), while correlation of Pearson coefficients was used for (C). The thickness and color of the lines connecting species indicate the strength of the correlation (thicker lines represent stronger correlations).

In addition to these broad evolutionary trends, we also observed clear taxon-specific differences in brain N-glycosylation (Supplementary text, Fig S10, S11B, summarized in Fig 3H), including several human-specific features of the brain N-glycome that were predominantly related to differences in Neu5Ac linkage (Supplementary text).

### Primate brain N-glycomes have diverged more rapidly than the underlying transcriptomic framework

We next investigated the relationship between glycogene expression and N-glycosylation in primate brains by integrating our N-glycomics data with complementary bulk RNA-Seq data (*17*). To narrow our focus to biologically relevant genes, we compiled a list of 47 glycogenes encoding enzymes that are predominantly involved in N-glycan synthesis (Table S9). Clustering the primate glycogene expression profiles according to overall similarity, we observed that, as was the case with the N-glycome, the CBC had the most distinct expression profile which overrides species differences (Fig S19). However, unlike the N-glycome, subsequent clustering was much more dependent on brain region than species indicating that inter-species differences are less significant when it comes to glycogene expression.

We further explored the extent to which glycogene expression in the brain is conserved among primates using Mantel correlograms to assess the inter-species correlation between each primate pair for the entire set of 47 glycogenes (Fig 4A). This resulted in strong correlations indicating that glycogene expression in primate brains is very highly conserved compared to non-glycogenes (Fig S20). Repeating the procedure for the primate brain N-glycome profiles resulted in weaker correlations (Fig 4B) indicating that N-glycosylation of brain proteins is generally less conserved than the expression of the associated glycosylation enzymes. Next, Pearson correlation coefficients were calculated to determine the linear correlation between glycogene expression and the N-glycome in each primate species. Inter-species correlation of these coefficients produced weaker correlations still (Fig 4C) indicating poor conservation of the relationship between these two traits across evolution and supporting previous observations that N-glycosylation status cannot always be predicted solely by the transcriptome (*19*, *20*). Lastly, we demonstrate that although the correlation between glycogene expression and N-glycosylation phenotype in primate brains is generally weak, it’s nevertheless possible to uncover putative associations between N-glycan structures and the expression of specific glycogenes using a bioinformatics prioritization strategy (Supplementary text, Fig S21). Overall, our observations of evolutionary divergence at the N-glycome level are consistent with examples of inter-species glycan diversity at the level of orthologous glycoproteins (*21*) and support the notion that divergent post-translational modification of conserved proteins via N-glycosylation is a means of generating additional diversity among genetically similar organisms.

### Regional and cell-type profiling of glycogene expression in primate brains

We compared glycogene expression across human brain regions and observed that the majority of glycogenes (29/47) could be classified into one of a handful of recurring expression profiles exhibiting clear regional specificity (Fig 4A). Of these, almost half showed either enriched or depleted expression specifically in the CBC. We validated the expression of a selection of glycogenes using digital droplet PCR (ddPCR) (Fig S22). In keeping with the clustering data, the spatial expression patterns of glycogenes are mostly conserved among primates (Fig S23), though there were some notable exceptions. We identified four glycogenes exhibiting a human-specific spatial expression profile in the brain; *B3GALT5*, *ST3GAL2*, *ST6GAL2*, and *ST8SIA6* (Fig S24). Intriguingly, all of these glycogenes were differentially expressed in the STR.

Next, we made use of primate dlPFC single-nucleus RNA-Seq (snRNA-Seq) expression data (*22*) to characterize glycogene expression across neural cell types. Clustering the glycogenes according to their expression profile revealed distinct heterogeneity in their cell-type distribution (Fig 5B, Fig S25). Generally, neurons have a broader palette of glycogene expression than non-neuronal cell types since expression of glycogenes from Clusters 3-5 is largely absent from the latter. Interestingly, there were few differences in glycogene expression between excitatory and inhibitory neurons. Although the overall clustering pattern was largely conserved in primates, some evolutionary divergence was evident (Fig 5B-C). In particular, our data show that the vast majority of differences in glycogene expression between macaques and hominids occurred in non-neuronal cell types in which glycogene expression has been extensively repressed in the hominid lineage. Interestingly, the expression of three sialyltransferases involved in the addition of α(2-6)-linked Neu5Ac to glycoconjugates (*ST6GAL1*, *ST6GALNAC3*, and *ST6GALNAC5*) was notably enriched in the human lineage across several cell types which may be one of the factors contributing to the increased abundance of α(2-6)-linked Neu5Ac in humans.

**Fig. 5.**
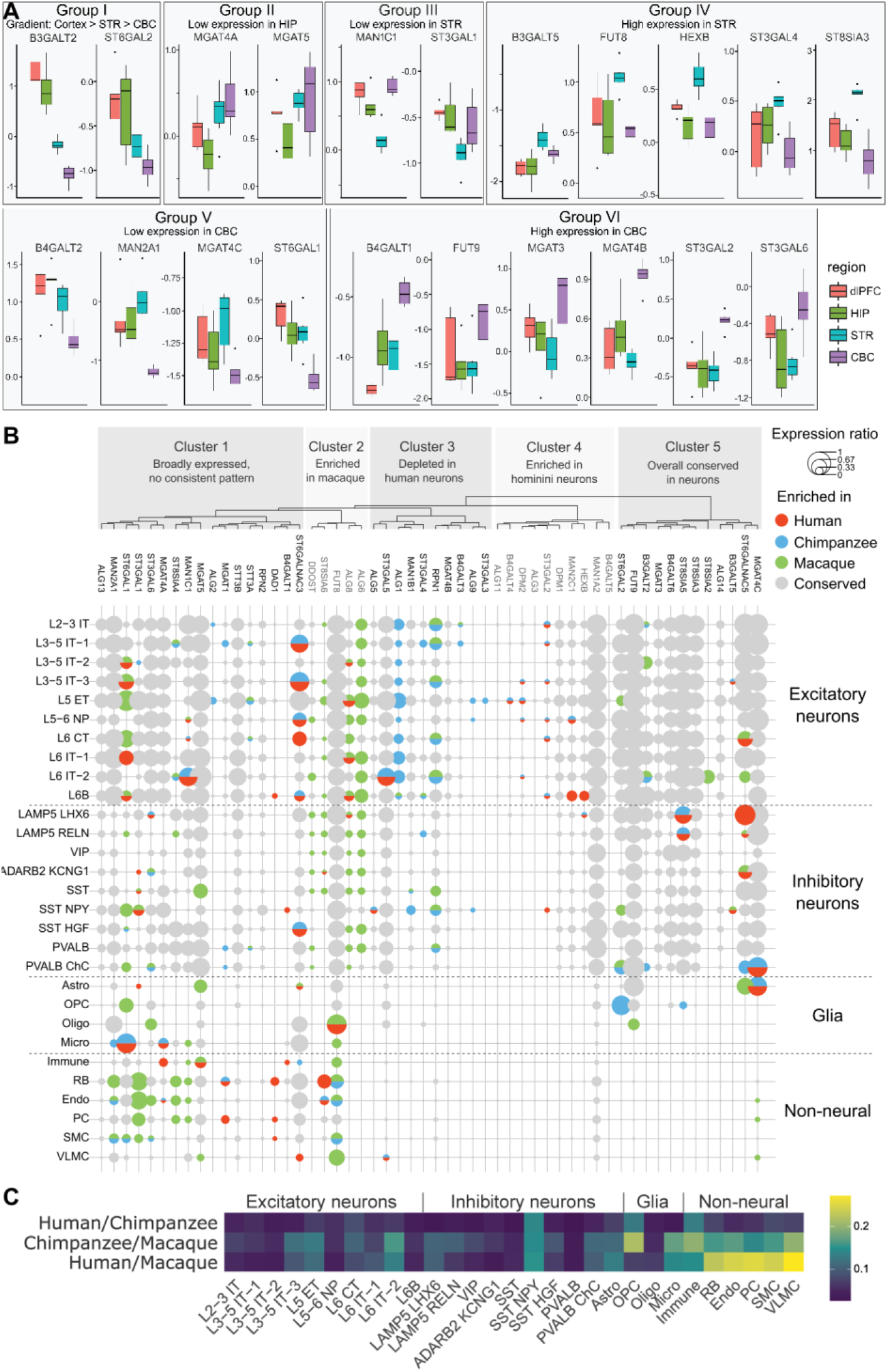
Human-specific and cell type-specific expression of certain glycogenes. (**A**) Categorization of glycogenes according to shared spatial expression profiles in human brains. The y-axes show scaled expression of logRPKM values within samples and among the 47 glycogenes. (**B**) Cell-type distribution of glycogene expression in the adult primate dlPFC. Dot color represents species expression enrichment or conservation (grey). Dot size indicates the average expression ratio in the enriched species (species-specific expression) or all the species (conserved expression). The dendrogram shows clustering of glycogenes according to their pattern of cell-type expression across species. (**C**) Phylogenetic divergence of glycogene expression in the primate dlPFC across various cell types. Here the dissimilarity in aggregate glycogene expression between pairs of primate species in each cell cluster was measured as the Pearson correlation coefficients of the average expression subtracted from 1 and is shown as a heatmap. Cell type abbreviations: L, layer; IT, intratelencephalic neurons; ET, extratelencephalic neurons; NP, near-projecting; CT, cortical-thalamic projection neurons; ChC, chandelier cells; Astro, astrocytes; OPC, oligodendrocyte precursor cells; Oligo, oligodendrocytes; Micro, microglia; Immune, immune cells; RB, red blood lineage cells; Endo, endothelial cells; SMC, smooth muscle cells; VLMC, vascular leptomeningeal cells.

### Human neurodevelopment is characterized by extensive changes in N-glycosylation

Next, we investigated the developmental pathways that shape the adult brain N-glycome by analyzing the corresponding brain regions in the prenatal human brain using a similar multi-omics approach. We show that the mid-fetal brain has already acquired the archetypal “brain-type” N-glycosylation profile by post-conception week (PCW) 21 (Fig S26). This early specification of the brain N-glycome has also been observed in rats and may be a conserved feature of mammalian neurodevelopment (*23*). Despite the overall similarity between the prenatal and adult brain N-glycomes (Fig S26-S30, Table S10), specific qualitative differences were observed at the level of individual chromatographic peaks that were either widespread (Fig S31) or region-specific (Fig S32-34) and ultimately reflected developmental differences in in the global composition of the N-glycome (Fig S35). We also identified numerous N-glycan peaks and structures that were enriched in either the fetal or the adult brain (Supplementary text, Fig S36-39). Hierarchical clustering and principal component analysis of the global N-glycome showed that N-glycosylation in the mid-fetal brain is regionally more homogeneous than in the adult brain and, moreover, that the CBC N-glycome has not yet become differentiated (Fig 6A-B). Interestingly, this observation contrasts with findings in which regional differences in gene expression were more pronounced during prenatal development than in adulthood (*24*). This may indicate that the more diverse N-glycome of the adult brain compensates for the restricted spatial diversity of the transcriptome and provides an additional layer of regional differentiation. When examining only sialylation features, we observed further regional differentiation in the adult brain with the HIP diverging from the other forebrain regions (Fig 6C). We identified four N-glycan clusters whose abundance was differentially regulated throughout neurodevelopment, particularly in the CBC (Fig 6A). Together, our data suggest that region-specific N-glycosylation programs lead to greater diversification of the human brain N-glycome throughout neurodevelopment, particularly in the CBC which acquires a distinctive N-glycome between the late mid-fetal stage and adulthood.

**Fig. 6.**
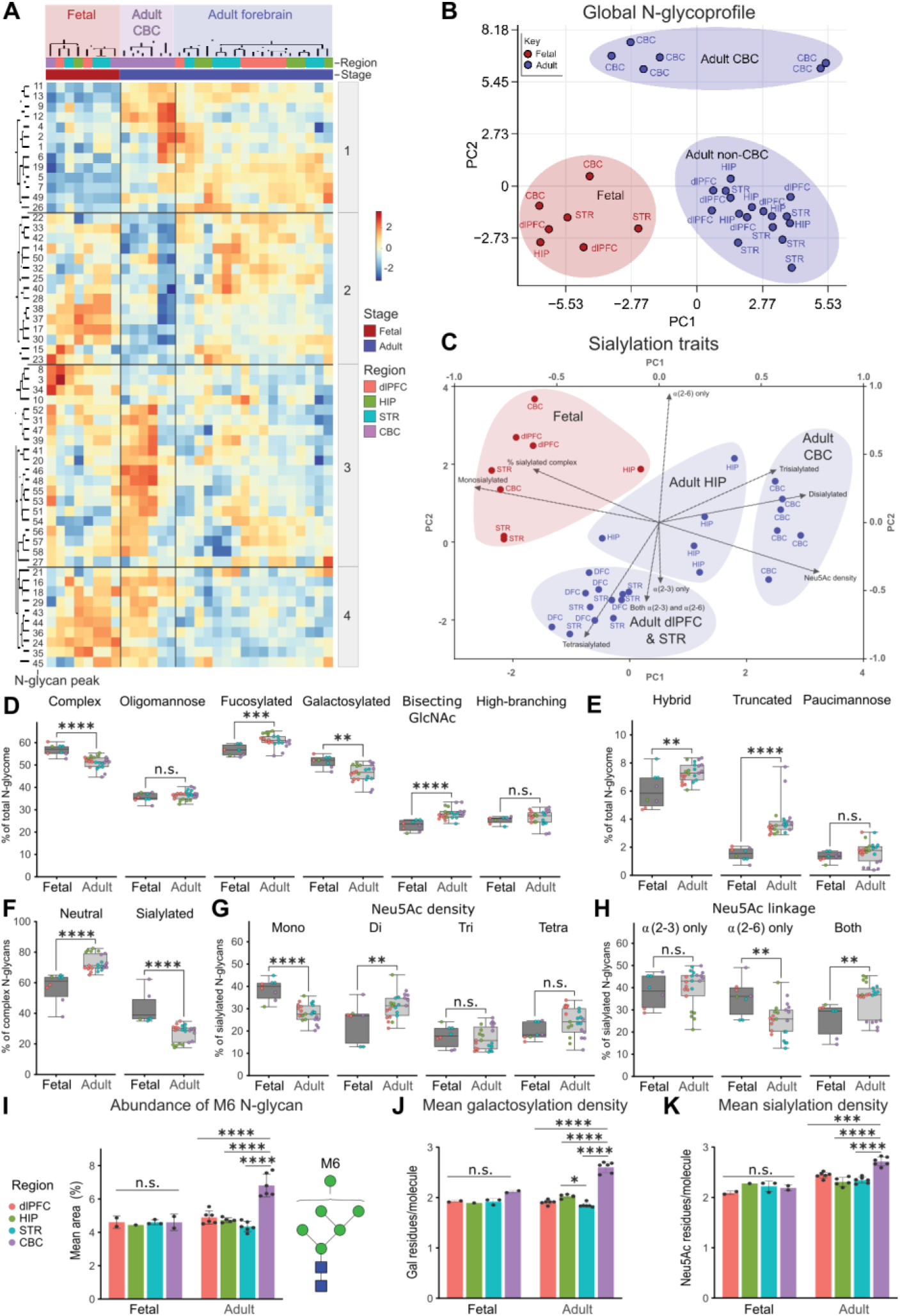
Global and region-specific maturation pathways lead to distinct spatial N-glycosylation profiles in the adult human brain. (**A**) Hierarchical clustering of quantitative UPLC N-glycan peak abundance. The heatmap shows the relative abundance of each chromatographic peak across samples. Developmentally-regulated N-glycan peak clusters are indicated (1-4). (**B-C**) Scatter plots illustrating the degree of relatedness among the human brain N-glycomes throughout ontogeny according to total the global N-glycome (B) or sialylation features only (C). The axes display the first (PC1) and second (PC2) principal components. In (C), the left and bottom axes refer to the loadings, while the top and right axes refer to the scatter plot. (**D-H**) Global developmental trends in brain N-glycosylation. Aggregate data for the indicated derived traits were analyzed using unpaired, two-tailed t-tests. Box and whisker plots show the data spread including the median value, interquartile range, maximum, and minimum. Data points are colored by brain region. (**I-K**) CBC-specific features of N-glycome maturation. Data for the indicated derived traits were analyzed using ANOVA with correction for multiple testing using the Bonferroni procedure. Adjusted P-values: **** 0.0001>P, *** 0.001>P>0.0001, ** 0.01>P>0.001, * 0.05>P>0.01, n.s. – not significant.

Global maturation of the brain N-glycome, which proceeds similarly in all regions (Fig S40), is characterized by an increase in the proportion of hybrid and truncated N-glycans at the expense of complex types, predominantly those that are sialylated (Fig 6D-F, Fig S41). The proportion of oligomannose N-glycans remained relatively stable across development throughout all brain regions (Fig 6D), though we observed a slight redistribution of the major oligomannose subtypes with the profile being skewed towards smaller subtypes in adults, mainly at the expense of the M9 form (Fig S42). These findings are consistent with results from rodent studies (*23*, *25*, *26*), though the shift in subtype usage seen in the human brain is not as marked. Notably, at 21 PCW the CBC does not yet exhibit elevated levels of the M6 subtype (Fig 6I). Regarding structural features, the proportion of N-glycans modified by either sialylation or galactosylation decreases as neurodevelopment proceeds, while the prevalence of fucosylated structures and those with bisecting GlcNAc increases (Fig 6D). We also observed changes in the density of such modifications (Supplementary text, Figs S43). CBC-specific maturation of the N-glycome was characterized by an increase in the abundance of the M6 structure along with increases in both galactosylation and sialylation density (Fig 6I-K). Finally, our data indicate that the human brain N-glycome undergoes extensive changes in the regulation of N-glycan sialylation during neurodevelopment (Fig 6G-H, Supplementary text, Fig S44-45). Briefly, we found that a smaller proportion of N-glycan structures are sialylated in the mature brain than the developing brain, though the ones that are sialylated tend to carry a greater number of Neu5Ac residues and, moreover, there is a trend towards increased usage of α(2-3)-linked Neu5Ac which is most prominent in the dlPFC and STR. Interestingly, similar developmental trends were observed in the mouse cerebral cortex (*27*) indicating that these may be conserved features of mammalian N-glycome maturation.

### Spatiotemporally-regulated glycogene expression in the human brain

Integration of our developmental N-glycomics data with complementary bulk RNA-Seq transcriptomic data enabled us to identify numerous developmentally-regulated glycogenes that showed either global or region-specific temporal dynamics (Supplementary text, Fig S46), a selection of which were validated using ddPCR (Fig S47). Generally, their expression profiles closely matched the developmental trajectory of orthologous genes in the mouse cerebral cortex (*26*) indicating that the developmental regulation of specific glycogenes in the brain is well conserved in Euarchontoglires. In accordance with the CBC’s distinctive N-glycome, the prevalence of region-specific developmental regulation of glycogene expression was highest in this brain region (Supplementary text).

Lastly, we searched for associations between developmental changes in glycogene expression and the abundance of specific N-glycan peaks in corresponding brain regions to uncover potential causal relationships between the two variables using a bioinformatics prioritization strategy (Supplementary text, Fig S48). Our analysis shows that, in certain instances, the differential expression of individual glycogenes can be linked to tangible and glycobiologically plausible changes in the N-glycome phenotype in a complex *in vivo* biological system, a finding that complements the work of others (*15*).

## Discussion

Our analysis revealed a wealth of anatomical, phylogenetic, and ontological variation in brain N-glycosylation patterns of Euarchontoglires mammals within the overall framework of a conserved template. Some of the most striking trends were related to changes in the relative abundance of complex versus non-complex N-glycans. Interestingly, evidence suggests that it’s the latter, and particularly the ancient oligomannose class of N-glycans (*28*), that are essential for normal neuronal functioning while complex N-glycans are dispensable (*29*). It’s unsurprising, therefore, that high oligomannose content is a conserved feature of the brain N-glycome – one that many CNS pathogens have evolved to exploit (*30*). In contrast, complex N-glycans in the CNS appear to be under fewer evolutionary constraints thus rendering them susceptible to progressive tinkering (e.g. switching Neu5Ac linkage) which can lead to innovation in particular phylogenetic lineages. As a result of overlapping anatomical and phylogenetic gradients in the abundance of complex N-glycans, peak N-glycome complexity was found in the hominid cortical regions, leading us to hypothesize that this increased diversity and complexity of sugar modifications found on neural N-glycoproteins contributed to the emergence of novel cognitive functions, including those unique to the human neocortex (*31*–*33*).

The other striking evolutionary trend concerned the shift towards increasing usage of α(2-6)-linked Neu5Ac in humans. A similar human-specific shift in Neu5Ac linkage has been reported in externally exposed tissues (i.e. airway epithelia and skin) and it’s thought that these changes were elicited by interactions with Neu5Ac-binding pathogens, such as the animal influenza virus which has a preference for α(2-3)-linked Neu5Ac (*34*). Our data provide the first evidence that α(2-6)-linked Neu5Ac is also enriched in the human brain, indicating that this shift may be more widespread than previously recognized. While the biological significance of this shift is still unclear, given that the influenza virus can invade the CNS (*35*), we speculate that brain cells may also have been under strong selective pressure to switch their Neu5Ac linkage.

Our data show that, in comparison to glycogene expression networks, the brain N-glycome evolves rapidly and, given our results, we propose that the evolution of the brain N-glycome has been shaped by two opposing forces. On the one hand, an increase in the proportion of complex N-glycans would increase diversity and hence provide more scope for the development of novel functions while at the same time enabling the host to evade pathogens that have evolved to exploit the brain’s high oligomannose content. However, reducing oligomannose content too much may compromise essential neurobiological processes which would negatively impact fitness. In such a trade-off, a delicate balance needs to be maintained, which may explain why the evolution of the brain N-glycome has been so incremental and, consequently, why the archetypal brain N-glycoprofile is so well conserved.

Regarding neurodevelopmental dynamics, we propose a model for the temporal transfiguration of the human brain N-glycome that incorporates elements of both global and region-specific maturation leading to greater spatial diversification over the course of brain development. We hypothesize that the changing neuroglycome reflects the evolving cellular landscape and functional requirements of a maturing organ (*23*). Some of these changes may arise due to changes in the cell composition of developing brain regions, such as the emergence of region-specific cell populations, and the resulting expression of cell-specific glycoproteins. For example, we hypothesize that the marked transformation of the cerebellar N-glycome is due to the emergence of granule cells, the majority of which appear postnatally (*36*).

Many of the developmental trends and developmentally-regulated N-glycans described here overlap with those seen in the rodent brain suggesting that these may be conserved, and thus important, features of brain N-glycome maturation in mammals (*23*, *26*, *37*, *38*). However, our work has also uncovered some novel developmentally-regulated N-glycans that have not previously been associated with brain development in rodents. This suggests that there are also species-specific differences in the regulation of individual N-glycan structures during neurodevelopment and further investigation of these carbohydrates, and their carrier glycoproteins, may provide insights into aspects of brain development that are unique to different species. Ultimately, although glycomics studies still pose a methodological and analytical challenge, our findings demonstrate that an understanding of brain function, evolution, and development, including human-specific specializations, is incomplete without knowledge of the sugar modifications that are found on neural glycoproteins.

## Funding

National Institutes of Health grant MH124619 (NS)

National Institutes of Health grant MH116488 (NS)

National Institutes of Health grant DA053628 (NS)

National Institutes of Health grant R01HG010898-01 (NS, GS)

Instituto de Salud Carlos III (Spain) grant MS20/00064 (GS)

Agencia Estatal de Investigación (Spain) grant

PID2019-104700GA-I00/AEI/10.13039/501100011033 (GS)

European Union Seventh Framework Programme (FP7 2007 2013) under grant agreement n° 291823 Marie Curie FP7-PEOPLE-2011-COFUND (The new International Fellowship Mobility Programme for Experienced Researchers in Croatia NEWFELPRO). This paper has been written as a part of the project “A spatio-temporal analysis of glycosylation in the human brain (Human Neuroglycome)” which has received funding through NEWFELPRO project under grant agreement n° 34 (TSK)

## Author contributions

Conceptualization: TSK, IG, GL, NS

Formal analysis: GS, FV, SM

Investigation: TSK, IG, MN, IB

Resources: CCS, JJE, PRH, DJ, NS

Writing - Original Draft: TSK, IG, NS

Writing - Review & Editing: TSK, IG, GS, AMMS, CCS, PRH, GL, NS

Visualization: TSK, IG, GS, FV, SM, NS

Supervision: GL, NS

Project administration: GL, NS

Funding acquisition: TSK, GS, GL, NS

## Data and materials availability

Raw MS and MS/MS data are available from the GlycoPOST repository under the following accession numbers: rat (GPST000311), macaque (GPST000309), chimpanzee (GPST000310), adult human (GPST000308), fetal human (GPST000312).

## Supplementary Materials

Materials and Methods

Supplementary Text

Figs. S1 to S48

Tables S1 to S10

References (39–*74*)

