## Supplementary materials for "Human-specific features and developmental dynamics of the brain N-glycome"

**This PDF file includes:**

Materials and Methods
Supplementary Text
Figs. S1 to S48
Captions for Tables S1 to S10

**Other Supplementary Material for this manuscript includes the following:**

Table S1 to S10 (Excel)

### Materials and Methods

Postmortem specimens

Information regarding the specimens used in this study is provided in Table S1. For this study, we used fresh postmortem brain tissue (rats) or high-quality, histopathologically-verified, neurotypical, fresh frozen brain tissue (primates). Tissue from all but one of the primate specimens has previously been used to generate other large-scale datasets and metadata regarding those specimens have been reported in earlier studies (*16*, *17*, *22*, *24*, *39–41*). Detailed descriptions of dissections and photo documentation are provided in these other studies. All experiments using non-human primates were carried out in accordance with a protocol approved by Yale University's Committee on Animal Research and NIH guidelines. The present study also complies with the definition of research exempt from the NIH Chimpanzee Research Use Panel (NOT-OD-14-024) regarding the use of chimpanzees or chimpanzee biomaterials. Following sample processing and data acquisition, samples not fulfilling quality control requirements were excluded. An overview of samples passing quality control is provided in Table S2.

Human (*Homo sapiens*): De-identified, clinically and neuropathologically unremarkable postmortem human brain tissue was procured from a bank of brain specimens; details regarding ethical approval and the collection process are reported in previous studies (*17, 24*). Adult tissue was sourced from six donors (male, n=3; female, n=3) ranging from 23-82 years of age (median age, 36.5 years). Prenatal tissue was sourced from three mid-fetal donors (male, n=1; female, n=2) ranging from 21-23.5 post-conception weeks (median age, 21 post-conception weeks). Where possible, tissue was sampled from four different brain regions from each donor; the dorsolateral prefrontal cortex (dlPFC or DFC), hippocampus (HIP), striatum (STR) and cerebellar cortex (CBC). The median post-mortem interval (PMI) was 15 h for adult specimens and 4 h for fetal specimens.

Chimpanzee (*Pan troglodytes*): Tissue samples were obtained from postmortem adult (age 23-31 years) chimpanzee brains (male, n=2; female, n=2) according to criteria previously described (*17*). All chimpanzees suffered sudden death with no prolonged agonal state and for reasons other than their participation in this study, and without any relation to the tissue used.

Rhesus macaque (*Macaca mulatta*): Tissue samples from the appropriate brain regions were dissected from postmortem adult (age 7-11 years) macaque brains (male, n=3; female, n=3) using criteria previously described (*17*). Rhesus macaques were euthanized for a previous study (*17*).

Rat (*Rattus norvegicus*): Animals were obtained from the University of Zagreb School of Medicine and with the approval of the ethics committee of the University of Zagreb, Faculty of Science (approval number: 251-58-10617-15-20). Whole brains were isolated from 2-month old Fisher 344 rats (male, n=3; female, n=3) and the appropriate brain regions were surgically microdissected. For the region corresponding to the primate dlPFC, the rat prefrontal cortex was dissected.

N-glycan release and labeling

An overview of workflow used in this study is provided in Fig S1. N-glycans were isolated from brain tissue, labelled with 2-aminobenzamide (2-AB), and prepared for chromatographic analysis as described previously (*14*), with minor modifications. Briefly, total protein content was isolated from brain tissue using chloroform/methanol extraction and denatured for 10 min at 95°C after which N-glycans were enzymatically released from glycoproteins using PNGase F (Promega, USA) over two overnight incubations at 37°C (5 units/incubation). The deglycosylation reactions were cleaned up using Amicon® Ultra 2 mL Centrifugal Filter Devices (Merck Millipore, Germany) according to the manufacturer's instructions and the purified N-glycans were labelled with 2-AB as previously reported (*42*). Free label and reducing agent were removed from the samples using a 0.2 µm GHP filter plate (Pall Corporation, USA). Glycans were eluted with ultra-pure water (2x 90 µL) and stored at –20°C until use.

<u>Ultra-performance liquid chromatography (UPLC)</u>

Fluorescently labeled N-glycans were separated by hydrophilic interaction liquid chromatography (HILIC) on an Acquity UPLC instrument (Waters, USA) as described previously (*14*) with excitation and emission wavelengths of 250 and 428 nm, respectively. The instrument was under the control of Empower 3 software, build 3471 (Waters). Labeled N-glycans were separated on a Waters BEH Glycan chromatography column (150 mm × 2.1 mm, 1.7 µm BEH particles) with 100 mM ammonium formate, pH 4.4, as solvent A and acetonitrile as solvent B. The samples were run at a flow rate of 0.561 mL/min using a linear gradient of 27-29.5% solvent A in the first 15 min and 29.5-38.7 % for the next 80 min. Samples were maintained at 5°C before injection, and the separation temperature was 25°C. Data processing was performed using an automatic processing method with a traditional integration algorithm, after which each chromatogram was manually corrected to maintain the same intervals of integration for all the samples. The chromatograms were thus integrated into 58 peaks (chromatographic peaks 1-58). The glycan abundance in each peak was expressed as a percentage of total integrated area. Chromatographic peaks were assigned glucose unit (GU) values based on their retention time relative to 2-AB-labelled glucose oligomers of known length belonging to the external dextran ladder standard, as previously described (*43*).

Matrix-assisted laser desorption/ionization time-of-flight/time-of-flight tandem mass spectrometry (MALDI TOF/TOF MS/MS)

The molecular masses of N-glycans and their monosaccharide compositions were determined by MALDI TOF/TOF MS/MS. Briefly, UPLC fractions corresponding to glycan peaks 1-58 were collected, dried, and resuspended in 2-5 µL of ultra-pure water depending on the length of the fraction. Linkage-specific ethyl esterification of N-glycans was performed to stabilize sialylated N-glycans and distinguish α(2-3)- and α(2-6)-linked sialic acids as previously described (*44*). Following ethyl esterification, samples were cleaned up using cotton HILIC solid phase extraction (*45*) and the N-glycans were dried, resuspended in 2 µL of ultra-pure water and spotted onto a MTP AnchorChip 384 BC MALDI target (Bruker Daltronics, Germany), mixed on plate with 1 µL of matrix solution (5 mg/mL 2,5-dihydroxybenzoic acid, 1 mM sodium hydroxide in 50% acetonitrile) and left to air dry. Recrystallization was performed by adding 0.2 µL of ethanol to each spot. Analyses were performed in reflectron positive (RP) mode on an UltraFlextreme MALDI-TOF-MS equipped with a Smartbeam-II laser and Flexcontrol 3.4 software Build 119 (Bruker Daltonics, Germany). The instrument was calibrated using a human plasma N-glycome standard. A 25 kV acceleration voltage was applied after a 140 ns extraction delay. A mass window of m/z 1000 to 5000 with suppression up to m/z 900 was used for brain N-glycan samples. For each spectrum, 10,000 laser shots were accumulated at a laser frequency of 2000 Hz, using a complete sample random walk with 200 shots per raster spot. Tandem mass spectrometry was performed on the most abundant peaks via laser-induced disassociation. All raw MS and MS/MS data are available from the GlycoPOST repository (*46*) under the following accession numbers: rat (GPST000311), macaque (GPST000309), chimpanzee (GPST000310), adult human (GPST000308), fetal human (GPST000312).

Annotation of chromatograms and assignment of proposed N-glycan structures to masses

The topological structure of N-glycans was deduced based on the convergence of evidence from several different sources, as previously described (*23*). Fig S2 shows the approach used for assigning N-glycan structures to masses. Briefly, the GlycoMod software tool (*47*) was used to obtain a list of theoretically possible monosaccharide compositions for each detected monoisotopic mass. The following fixed modifications were used: reducing end 2-AB, sodiated ($Na^+$). Since it has been shown that fucose is the only 6-deoxyhexose present in rat brain glycoproteins (*48*), 6-deoxyhexose values are reported as fucose. The list of potential compositions for each segment was additionally filtered by crosschecking the GU value of the segment against reference values in the GlycoBase 3.2.4 database (*49*) to eliminate N-glycans that would not be expected to elute at the given retention time (e.g. glycan fragments generated by in-source fragmentation). Information about topological structure and the sequence of monosaccharides in the carbohydrate chain was deduced from diagnostic ions present in MS/MS fragmentation spectra, where available, which indicate specific N-glycan structural features and thus aided in distinguishing between isobaric isomers. Finally, expert knowledge of the biosynthetic pathways involved in vertebrate N-glycan synthesis was coupled with evidence from the literature regarding the types of N-glycans present in brain tissue to provide an integrated approach to structural assignment. N-glycan structures were drawn using GlycoWorkbench (*50*) according to the standard symbol nomenclature for the representation of glycan structure proposed by the Consortium for Functional Glycomics (*51*).

Relative quantification of N-glycans and N-glycosylation traits

To make HILIC-UPLC measurements across samples comparable, normalization by total area was performed whereby the area of each of the 58 chromatographic peaks was divided by total area of the corresponding chromatogram. Batch correction was performed on normalized log-transformed measurements using linear mixed models (R package lme4), in which the technical source of variation was modeled as a random effect. Prior to statistical analyses, N-glycan variables were all log-transformed due to right skewness of their distributions. In addition to directly measured glycan structures (i.e. peak abundance data for single chromatographic peaks), derived traits were calculated from the HILIC-UPLC data. These derived traits group together common glycosylation features across different individual chromatographic peaks and, consequently, they should be more closely associated with specific enzymatic pathways responsible for N-glycan synthesis. Due to qualitative differences in the N-glycan content of chromatographic peaks, derived N-glycosylation traits were calculated individually for each species, brain region and developmental stage according to the formulas in Table S7.

Analyses of differences in N-glycan derived traits between sample groups were performed using the analysis of covariance (ANCOVA) model. When regional differences in N-glycosylation were analyzed within a single species, age and sex were included as additional covariates, while sex was included as a covariate when comparing N-glycomes across species. Correction for multiple testing was performed using the Bonferroni procedure.

Relative quantification of N-glycans within a chromatographic peak

For relative quantification of glycan peaks from MS spectra, absolute intensity values of the dominant isotopologue were manually extracted for each glycan composition in the spectrum and glycan abundance was expressed as a percentage of the cumulative intensity. Glycan structures were excluded if their relative abundance was <10% in all samples (i.e. across all species and brain regions for evolutionary comparisons or across all developmental stages and brain regions for developmental comparisons).

Primate bulk transcriptomics data analysis

Complementary primate gene expression data from age-matched individuals was obtained from publicly available databases: Non-Human Primate Brain Transcriptomes (www.sestanlab.org/resources), National Chimpanzee Brain Resource (www.chimpanzeebrain.org), Brainspan (*40*) (www.brainspan.org), and PsychENCODE (http://evolution.psychencode.org/#). For human samples, this corresponds to windows 4 and 9 for the mid-fetal and adult periods, respectively (*24*). Where possible, data from matching donors were used to complement the N-glycomic data and reduce noise introduced by inter-individual variability (5/6 adult human donors and 8/9 non-human primate donors were identical in both the N-glycomics and transcriptomics analyses, Table S1). We obtained log2 reads per kilobase million (RPKM) expression data from human, chimpanzee, and macaque RNA-sequencing (RNA-Seq) datasets for 47 glycogenes (Table S9). In the adult brain, initial clustering revealed separation by species rather than brain regions. We Z-scored expression values in each sample to express them as standard deviations from the mean expression of each sample. This data transformation rendered the expected clustering grouping first by region (particularly CBC and

STR), and secondly by species. Fold changes and P-values between fetal and adult were those determined in Li *et al*. 2018 using DESeq2.

Digital droplet polymerase chain reaction (ddPCR)

We employed droplet digital PCR (ddPCR) to reliably quantitate gene expression and thus validate the bulk RNA-Seq results. Briefly, 1 µg of total RNA was used for cDNA synthesis using SuperScript III First-strand synthesis Supermix (Invitrogen) and subsequently diluted with nuclease-free water. Custom gene-specific primers and probe for each gene of interest (Table S9) were designed using NCBI/Primer-BLAST (http://www.ncbi.nlm.nih.gov/tools/primer-blast/) and PrimerQuest tool (IDT). ddPCR was carried out using the Bio-Rad QX100 system. After each PCR reaction mixture consisting of ddPCR master mix and custom primers/probe set was partitioned into 15,000–20,000 droplets, parallel PCR amplification was carried out. Endpoint PCR signals were quantified, and Poisson statistics was applied to yield target copy number quantification of the sample. Two-color PCR reaction was utilized for the normalization of gene expression by the housekeeping gene *TBP*.

Primate single-nucleus RNA sequencing (snRNA-Seq) data analysis

Complementary primate dlPFC snRNA-Seq expression data was obtained from a previously published study (*22*). Here, the differential expression was performed between each pair of species in a given homologous cell cluster using the *FindMarkers* function in Seurat (*52*), with the significance evaluated by Wilcoxon Rank Sum test. Pie charts were used to summarize the expression patterns across species. We further clustered the glycogenes based on their expression patterns across cell clusters and species. Specifically, we calculated the average expression of each glycogene in each cell cluster and species using the *AverageExpression* function in Seurat followed by log-transformation. We then calculated the Spearman correlation coefficients between each pair of genes and defined the gene-gene distance by subtracting the coefficients from 1. The gene-gene distance matrix was passed to *hclust* function in R with the “ward.D2” algorithm to construct the dendrogram hierarchically organizing the glycogenes. For a global overview of the species divergence in glycogene expression, we measured the Pearson correlation coefficients of the average expression of glycogenes in a given homologous cell cluster between each pair of species, followed by subtracting from 1 to represent expression dissimilarity. The dissimilarities were visualized in Fig S5C.

Linear regression and quantile-quantile (Q-Q) plot

In each brain region, a linear model was performed for the glycans abundance against the evolutionary age using lm function in R. To evaluate the normality of the data across species, we obtained the standardized quantiles of the observed data via rstandard function in R, calculated the theoretical Gaussian quantiles, and plotted the results in dot plots (Q-Q plot) with dots colored by species.

Data analysis, vizualization and presentation

GraphPad Prism 8.4.3 was used for drawing graphs and the following basic statistical tests and analyses; t-tests, analysis of variance (ANOVA), mixed model analysis, and principal component analysis (PCA) with the exception of the PCA of the global N-glycome across human neurodevelopment which was performed using the analysis and visualization software BioVinci (version 1.1.5, r20181005). BioVinci was also used for t-distributed stochastic neighbor embedding (t-SNE) analysis and hierarchical clustering. The complete-linkage method and Euclidean distancing parameters were used for hierarchical clustering. GlyConnect Compozitor

(1.0.0) was used to construct the core brain N-glycan network (*53*) with the assumption that the core N-glycan structures are biosynthetically interrelated and are the products of a small number of N-glycan processing pathways that are dominant in the brain. Accordingly, the illustrated N-glycans depict the proposed structural configurations while also taking into account GU value, knowledge of N-glycan biosynthetic pathways, information from the literature, and MALDI TOF/TOF MS/MS fragmentation spectra (where available). Icons from BioRender were used for drawing figures.

Correlation of the N-glycome and glycotranscriptome within and among primate species

To avoid species/batch confounding effects in gene expression and peak abundance we calculated pairwise correlation matrices in glycogenes and peaks. Co-expression/co-abundance matrices were correlated between species using the Mantel test implemented in the ade4 R package. To test the significance of these correlations, we resampled groups of 47 genes 1000 times and obtained a distribution of Mantel's correlations between species. In matching samples, we also calculated the correlation between peaks and glycogenes independently in each species. In order to test how well the relationships between peaks and glycogenes were conserved, we then calculated the correlation of these Pearson's coefficients between species.

We further analyzed the relationship between glycogenes and peaks in human development using Sparse Partial Least Squares (sPLS) implemented in the mixOmics R package (*54*). We used two components to calculate the correlation structure between the abundance of glycan peaks and the expression of glycogenes measured in humans. We visualized relationships between glycogenes and peaks in a network using igraph package in R, removing unconnected nodes, and edges with correlations smaller than 0.6.Data were analyzed and visualized using R programming language (version 3.0.1).

Calculation of the glycogene specificity score

For each human brain region and time period (fetal and adult) we encoded the putative involvement between each glycogene and the dominant N-glycan structure in each individual N-glycan peak with a binary entry (1/0 for evidence/no-evidence) based on common knowledge of N-glycan biosynthetic pathways (*55*) in conjunction with available information regarding enzyme activity (*56*). For each glycogene we calculated a specificity score describing how specific the relationship is between the glycogene and a certain N-glycan peak, considering the involvement of the glycogene with the rest of the peaks. The score was calculated as:

$$S_z = \frac{\sum_i^n z - x_i}{n - 1}$$

where $S_z$ is the specificity score of a particular glycogene-peak relationship, Z is the evidence for functional relationship glycogene-peak and $x_i$ all possible (n) relationships.

### Supplementary Text

Structural features of Euarchontoglires brain N-glycomes

Our detailed structural analysis of brain N-glycans revealed a number of striking features that are consistent with the brain-specific pattern of N-glycosylation that has been well-described (*12*, *57–61*). In agreement with previous studies, we report that the Euarchontoglires brain N-glycome is characterized by high levels of core and antennary fucose, bisecting GlcNAc, terminal α(2-3)-linked Neu5Ac often forming the sialyl-Lewis$^x$ epitope {Neu5Acα(2-3)Galβ(1-4)[Fucα(1-3)]GlcNAc}, Neu5Ac α(2-6)-linked to GlcNAc forming the branched disialyl Lewis C motif (Fig S4), and the absence of *N-glyc*olylneuraminic acid (Neu5Gc). Several of the detected N-glycans carried covalent modifications such as sulfate groups (Fig S5), which were found on complex N-glycans (e.g. FA3SulfateG1 in chromatographic peak 10), or phosphate groups (Fig S6), which were present on oligomannose N-glycans (e.g. M7Phos in chromatographic peak 27). While both of these modifications have been reported to occur relatively more frequently on brain N-glycans than in other tissue types (*61–64*), in this study they were generally found on low-abundance structures. Interestingly, our data show that tri- and tetrasialylated N-glycans homogeneously adorned with α(2-6)-linked Neu5Ac do not occur on brain N-glycans (Fig S18). Given that we did not detect such structures in any brain region of any species, not in any region of the developing brain (Fig. S45), we hypothesize that the absence of such N-glycans is a conserved feature of the Euarchontoglires brain N-glycome, possibly indicating that there are biosynthetic constraints that regulate the processing of sialylated N-glycans in neural cells.

Gradient of increasing abundance of complex N-glycans from the hindbrain to the cortex

When brain regions were compared within a species using an ANCOVA, the differences were found to be statistically significant only in rats, likely due to their low inter-individual variability (i.e. genetic homogeneity along with identical age, living environment, and PMI). Since it is known that many N-glycosylation traits have a high degree of heritability (*65*) and that the proportion of oligomannose N-glycans in the brain increases with age (*12*), we reasoned that the higher inter-individual variability in age and genetic background in the primate cohorts may hamper the ability of such a test to detect trends at the whole group level, especially in the human cohort, which had the broadest age spread as well as the largest variability in PMI (Fig S3, Table S1). To mitigate this, we employed mixed model analysis to investigate this anatomical trend at the level of individual subjects and found that in all species there was an overall nominally significant linear trend of increasing complex N-glycan content from the CBC, through the STR, HIP and dlPFC (Fig S15). The strength of the linear trend was commensurate with the homogeneity of the cohort (Fig S3). Consequently, the more robust linear trends in rats and macaques ($P < 0.0001$) withstood correction for multiple testing using the Bonferroni method while those in humans ($P = 0.03$) and chimpanzees ($P = 0.003$) did not. Nevertheless, these data support the hypothesis that a conserved neuroanatomical trend in N-glycome complexity exists which may be partially obscured by inter-individual variability in the primate cohorts.

Taxon-specific differences in Euarchontoglires brain N-glycosylation

In addition to broad evolutionary trends, we also observed clear taxon-specific differences in brain N-glycosylation (Fig S10, S11B, and S14). Statistically significant differences are summarized in Fig 3H. When compared to rats, primates displayed several key differences in the way N-glycans are processed in the brain, namely; (i) a greater abundance of N-glycans modified with bisecting GlcNAc throughout the brain, (ii) a lower abundance of the M6 oligomannose structure throughout the brain, (iii) increased abundance of tetrasialylated L1E3 variants in the forebrain (discussed below), and (iv) higher levels of core fucosylation, primarily in the HIP. Notable changes specific to the hominids include; (i) a lower abundance of hybrid N-glycans in the dlPFC and CBC, and (ii) a greater proportion of N-glycans carrying solely α(2-6)-linked Neu5Ac in the CBC. Features specific to the human brain N-glycome were predominantly related to differences in the linkage of Neu5Ac and include; (i) a lower abundance of N-glycans carrying exclusively α(2-3)-linked Neu5Ac in the HIP, (ii) a greater abundance of N-glycans carrying exclusively α(2-6)-linked Neu5Ac in the dlPFC and HIP, (iii) a greater abundance of N-glycans carrying both α(2-3)- and α(2-6)-linked Neu5Ac in the STR, (iv), a greater abundance of neutral complex N-glycans in the HIP, and (v) increased galactosylation density in the CBC.

Primate-specific Neu5Ac usage pattern

Although tetrasialylated L1E3 variants [i.e. carrying one lactonized α(2-3)-linked Neu5Ac residue (L1) and three esterified α(2-6)-linked Neu5Ac residues (E3)] were relatively abundant in primate forebrain regions, these variants were detected only as minor structures in rat brains (Fig S18).

Weak correlation between glycogene expression and N-glycosylation in primate brains

To investigate the extent to which glycogene expression determines the final N-glycosylation profile, and therefore whether it can be used to predict the N-glycome, we explored the relationship between the two datasets. Overall, the global correlation between glycogene expression and the N-glycome across all matched samples is modest (r=0.42, P=0.01, 0.51

quantile; Fig S21A), and disappears completely if the CBC is excluded (r=0.34, data not shown), indicating that the highly conserved CBC is driving this weak global correlation. However, this does not preclude a more nuanced direct correlation between the two datasets. Therefore, we also calculated the Mantel's correlation between N-glycan peaks and glycogenes among groups of samples in the PCA space using the averaged Euclidean distance of the first ten principal components (PCs) by species and brain region, obtaining a higher correlation value (r = 0.73 when using all brain regions and r = 0.43 when excluding the CBC; Fig S21B-C), albeit non-significant in the permutations. Using individual samples (instead of averaging by species and brain regions) yielded a much lower N-glycome-glycogene correlation in the PCA (r = 0.5). We further investigated the relationship between the two datasets using individual matched samples and the number of PCs in each data modality which maximized Mantel's correlation: PC3 for glycogenes and PC4 for the N-glycome, which resulted in a higher correlation of r=0.603 placing it in the 0.85 quantile (Fig S21D-E).

We then searched for associations between individual glycogenes and chromatographic peaks in humans using sparse partial least squares (sPLS) regression to measure covariance in an unsupervised manner (Fig S21F-G). This enabled us to find highly correlated N-glycan peak/glycogene pairs from which a proposed regulatory network could be constructed (Fig S21H-I). Though some of the associations were contrary to basic glycobiology theory and thus likely to be purely coincidental, by manually curating the network using common knowledge of N-glycan biosynthetic pathways (*55*) in conjunction with available information regarding enzyme activity (*56*) we obtained a putative functional network outlining the enzymes predicted to be involved in the synthesis of particular N-glycan structures in human brains (Fig S21J). Of course, this model makes certain assumptions and the identified associations need to be validated experimentally. Nevertheless, our analysis demonstrates that this could be a useful approach for investigating the intricate relationship between glycosylation enzymes and glycan products in a complex *in vivo* biological system and thus complements the work of others in the field (*15*, *66*).

Developmentally-regulated N-glycans in the human brain

We identified numerous N-glycan peaks and structures that were specifically enriched in either the fetal or the adult brain (Fig S37-39, Table S3). The majority of these developmentally-regulated N-glycans were adult-specific indicating a greater diversity of structures in the mature brain. These developmentally-regulated oligosaccharides were sometimes present brain-wide, such as the adult-specific FA4F4G4 structure in Peak 54a (H7N6F5, 3244.20 *m/z*) and the fetal-specific FA2F1G2S[3]1 structure in Peak 29 (H5N4F2L1, 2348.85 *m/z*). The composition of the latter corresponds to an N-glycan that is enriched in the young postnatal human prefrontal cortex (*13*), though additional structural analysis will be required to determine whether they are equivalent N-glycans. Other times, the developmentally-regulated oligosaccharides were region-specific, such as the two adult-specific complex N-glycans in Peak 35 (H6N5F2L1, 2713.98 *m/z* and H5N7F3, 2831.06 *m/z*) that were detected only in the adult CBC, as well as the fetal-specific N-glycan in Peak 38 (H5N4F1L2E1, 2795.00 *m/z*), which was specifically absent from the mid-fetal CBC.

Relative decrease in bigalactosylated N-glycans throughout human brain development

We observed a specific reduction in the proportion of bigalactosylated N-glycan structures in the adult human brain (Fig S43B). We hypothesize that this curious redistribution of galactosylation density is due to the increased prevalence of truncated N-glycans in the adult brain. In the rodent brain, these structures are generated via the action of HEXB, an enzyme that is more highly expressed in adults than in developing animals and which trims biantennary N-glycans to produce monoantennary N-glycans which are either monogalactosylated or agalatosylated (*26*, *67*).

Changes in sialylation occurring during human neurodevelopment

Strikingly, we discovered that there are some combinations of Neu5Ac that do not occur on N-glycans of the human brain. Namely, although we detected tri- and tetrasialylated N-glycans that were homogeneously adorned with α(2-3)-linked Neu5Ac, we did not detect their α(2-6)-linked Neu5Ac counterparts (Fig S45). Other combinations appear to be developmentally-regulated. For example, while the trisialylated L2E1 variants [i.e. carrying two lactonized α(2-3)-linked Neu5Ac residues (L2) and one esterified α(2-6)-linked Neu5Ac residue (E1)] are the most common trisialylated variants in the fetal forebrain regions, these were only detected as minor structures in the adult brain. In contrast, the tetrasialylated L1E3 variants are abundant in the adult forebrain regions, yet are minor structures in the fetal brain.

Glycogenes whose expression is globally up-regulated in the fetal human brain

The glycogenes whose expression was most highly enriched throughout all regions of the fetal brain were *ST8SIA2* and *ST8SIA4*, which encode the two enzymes exclusively responsible for the synthesis of polysialic acid (PSA) chains on glycoproteins (*68*). Their developmental enrichment mirrors findings from rodent studies which have demonstrated that these polysialyltransferases, as well as PSA itself, are highly expressed in the developing rodent brain, where they promote plasticity and cell migration (*69*–*72*). Other genes whose expression was enriched throughout the fetal brain include *MGAT4B*, *B4GALT2*, *B4GALT5*, and *ST6GAL2*. In agreement with our data, the expression of *B4GALT2* was previously shown to be enriched in the fetal human brain by northern blot (*73*). Increased fetal expression of the two galactosyltransferases, *B4GALT2* and *B4GALT5*, may account for the higher levels of galactosylation observed in the developing brain. The expression of *ST6GAL2*, which encodes an enzyme that preferentially transfers Neu5Ac to a GalNAcβ(1-4)GlcNAc (LacdiNAc) acceptor via α(2-6)-linkage (*56*), was enriched in the developing brain. Its reduced expression in the mature brain may contribute to the shift towards greater usage of the α(2-3)-linkage observed in adults.

<u>Glycogenes whose expression is globally up-regulated in the adult human brain</u>

Glycogenes that were widely upregulated in the adult brain include *MAN1C1*, *MAN2A2*, *B4GALT6*, *ST6GALNAC6*, and *HEXB*. In support of our data, northern blot analysis of human brain tissue has previously shown that *B4GALT6* is more highly expressed in adulthood than during fetal development (*73*). Similarly, increased expression of *HEXB* has also been reported in the adult rodent brain (*26*). The increased adult expression of the *MAN1C1* and *MAN2A2* mannosidases might be expected to result in an increase in the abundance of the M5 oligomannose structure and complex N-glycans, respectively, which is not entirely reflected in the adult brain N-glycome. However, these are processing steps that are catalyzed by multiple enzymes and their redundancy likely accounts for the lack of a clear effect on the glycosylation phenotype.

<u>Glycogenes exhibiting region-specific developmental regulation in the human brain</u>

Expression of *B4GALT1* and *FUT9* was enriched specifically in adult CBC, while *B3GALT2* was upregulated only in the fetal CBC. There were even several instances in which glycogenes exhibited a converse temporal expression trajectory in the CBC compared to other regions, e.g.

the enrichment of *MAN2A1* and *CHST2* expression in the fetal and adult CBC, respectively. Other examples of region-specific developmental regulation include the enrichment of *FUT7* and *ST6GALNAC2* expression in the fetal dlPFC, as well as the enrichment of *ST8SIA5* expression in the fetal STR. The significance of these regional differences in glycogene expression is unknown and they do not seem to be reflected in the overall glycosylation trends of the respective brain regions. It is possible that they affect only a small number of specific N-glycans and thus their effects are undetectable at the macro level. Alternatively, additional factors, such as enzyme redundancy or substrate availability, may dampen the impact of altered glycogene expression on the N-glycome.

<u>Deducing the involvement of glycogenes in the generation of specific N-glycan structures via multi-omic correlation</u>

Based on the dominant N-glycan structure in each chromatographic peak of the human brain N-glycome, we used common knowledge of N-glycan biosynthetic pathways (*55*) in conjunction with available information regarding enzyme activity (*56*) to generate a specificity score for each glycogene/peak pair that describes the expected involvement of that enzyme in generating the major N-glycan structure in that particular peak. The specificity score ranges from 1, if the glycogene is only involved in the generation of that particular peak, to -1, when the glycogene is involved in the generation of all peaks except for that particular peak. This score was then integrated with data about glycogene/peak pairs whose expression levels changed in the same direction throughout development.

Using this approach, we were able to infer direct relationships between the expression of certain glycogenes and the abundance of specific N-glycan structures (Fig S48). For example, increased

*HEXB* expression in adults was associated with an increased abundance of Peak 5. This supports the hypothesis that HEXB-dependent conversion of the biantennary bisected FA2B N-structure (H3N5F1, Peak 7, 1808.62 *m/z*) to the truncated bisected FBA1 structure (H3N4F1, Peak 5, 1605.60 *m/z*) is the main factor leading to the increased abundance of Peak 5 in the mature brain (*23*, *67*). Similarly, increased expression of *ST8SIA3* and *ST8SIA5* in the adult dlPFC correlated with increased abundance of Peak 54 from which we infer that these two enzymes from the α(2-8)-sialyltransferase family are involved in the production of the putative Neu5Acα(2-8)Neu5Acα(2-3)Gal disialic motif present on the dominant N-glycan of that chromatographic peak. Of course, this model makes certain assumptions and the identified associations need to be validated experimentally as some of them may be coincidental. Nevertheless, our analysis demonstrates that this could be a useful approach for identifying candidate glycogenes involved in the synthesis of specific N-glycans and, hence, generating hypotheses for future experiments.

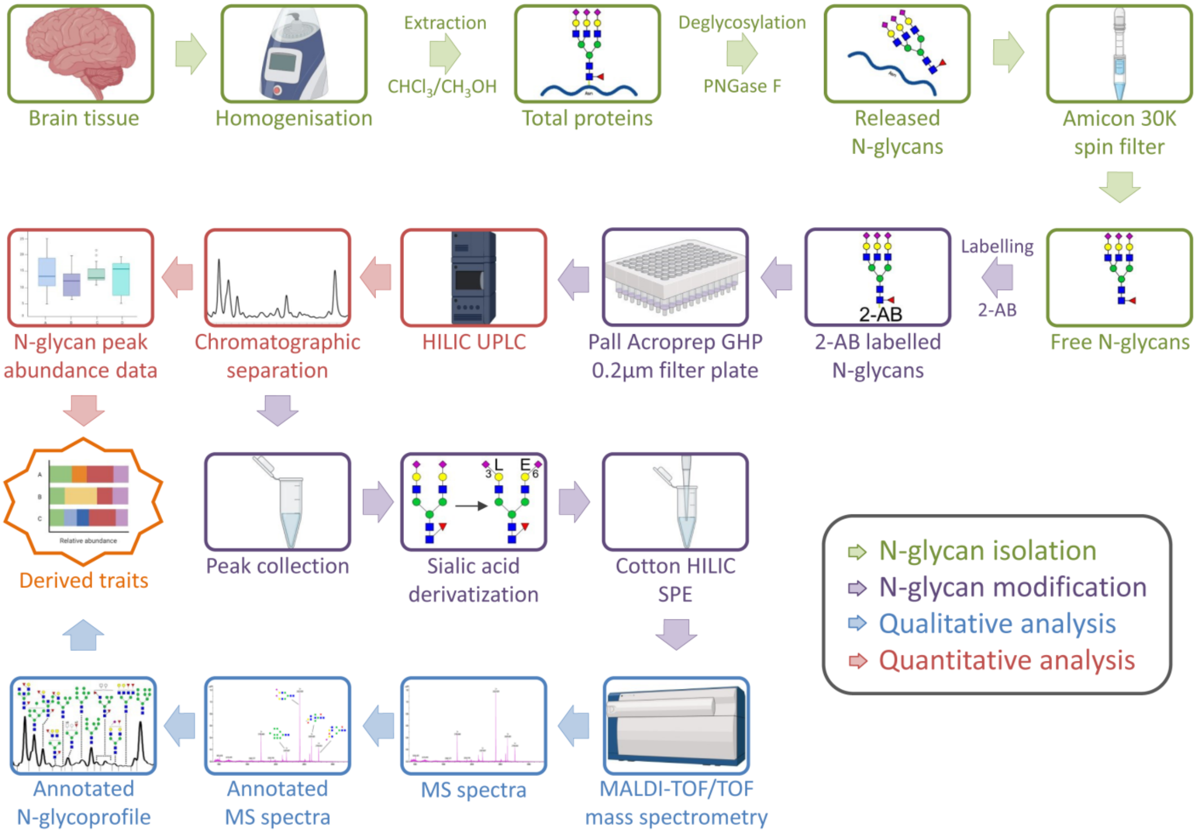


**Fig. S1. Study workflow.** Brain tissue was homogenized using ceramic beads and total protein was extracted from the homogenate using chloroform/methanol extraction. N-glycans were enzymatically released from glycoproteins using PNGase F and purified by size exclusion using a centrifugal filter with a 30 kDa molecular weight cut-off. Purified N-glycans were labelled with the fluorophore 2-AB at the reducing end via reductive amination which was followed by solid phase extraction (SPE) using a 0.2 µm filter plate coupled to a vacuum manifold to remove excess dye. N-glycans in the complex mixture were separated using HILIC UPLC and chromatographic fractions corresponding to N-glycan peaks were collected and dried. Ethyl esterification was performed to enable differentiation of α(2-3)- and α(2-6)-linked Neu5Ac and this was followed by cotton HILIC SPE. N-glycans from individual chromatographic fractions were then analyzed by MALDI TOF/TOF MS/MS and the obtained MS and fragmentation spectra were manually annotated.

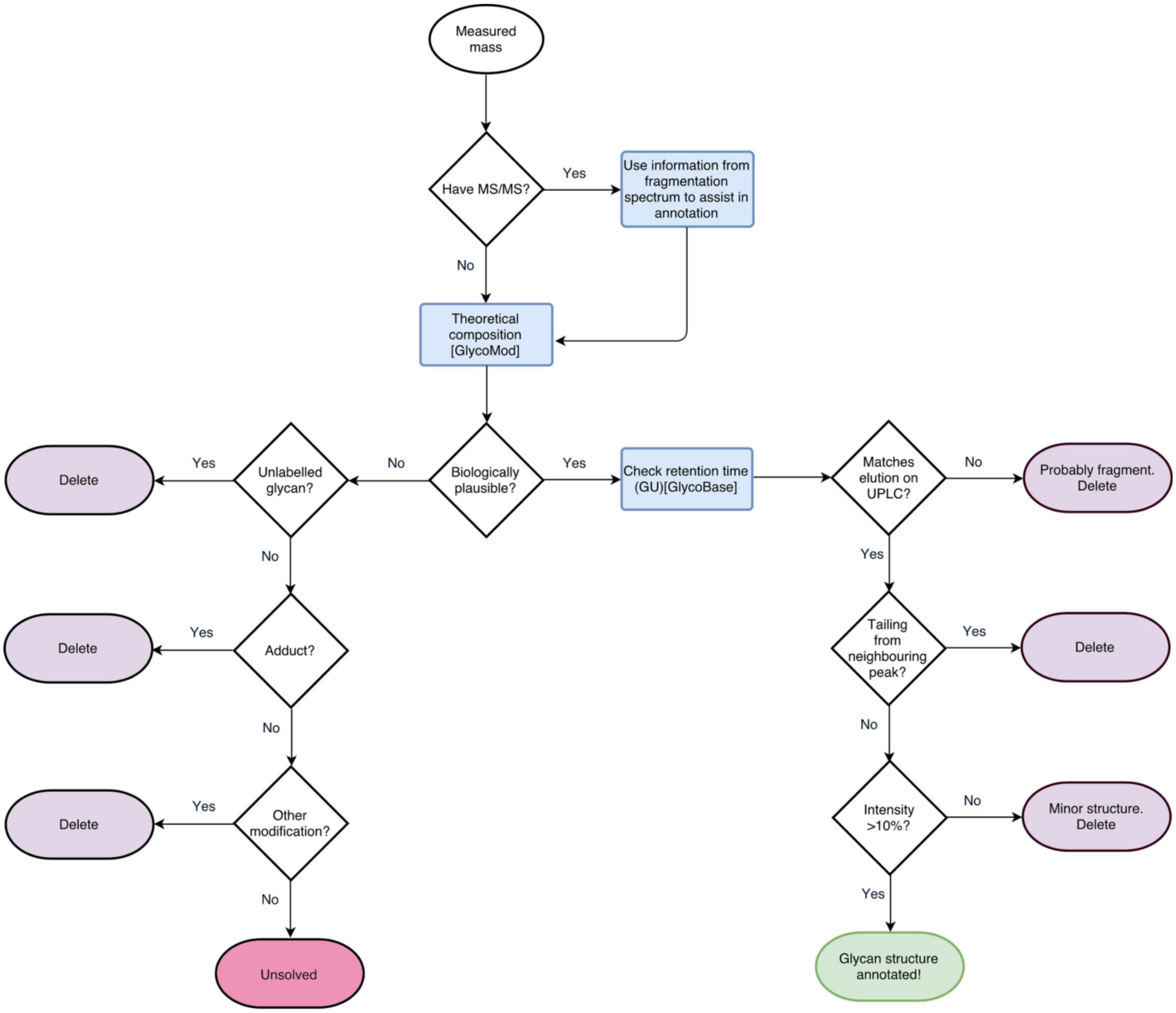


**Fig. S2. Approach used for assigning N-glycan structures to masses.** Flow-diagram illustrating the integrative approach used for N-glycan structural assignment.

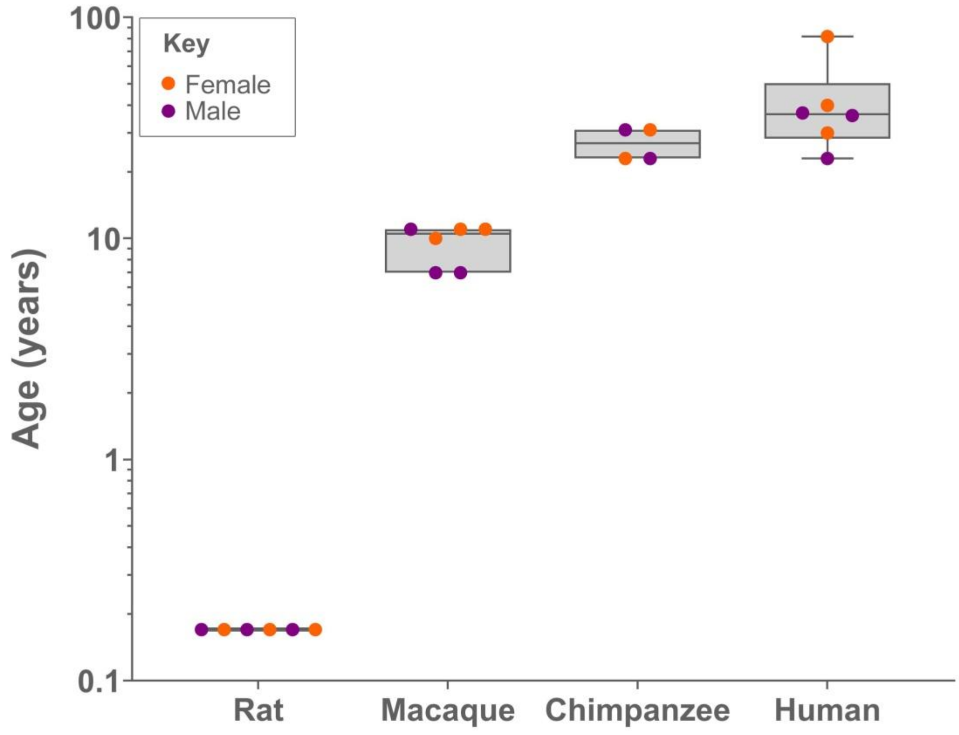


**Fig. S3. Age spread of samples in each species.** Note the logarithmic scale on the y-axis.

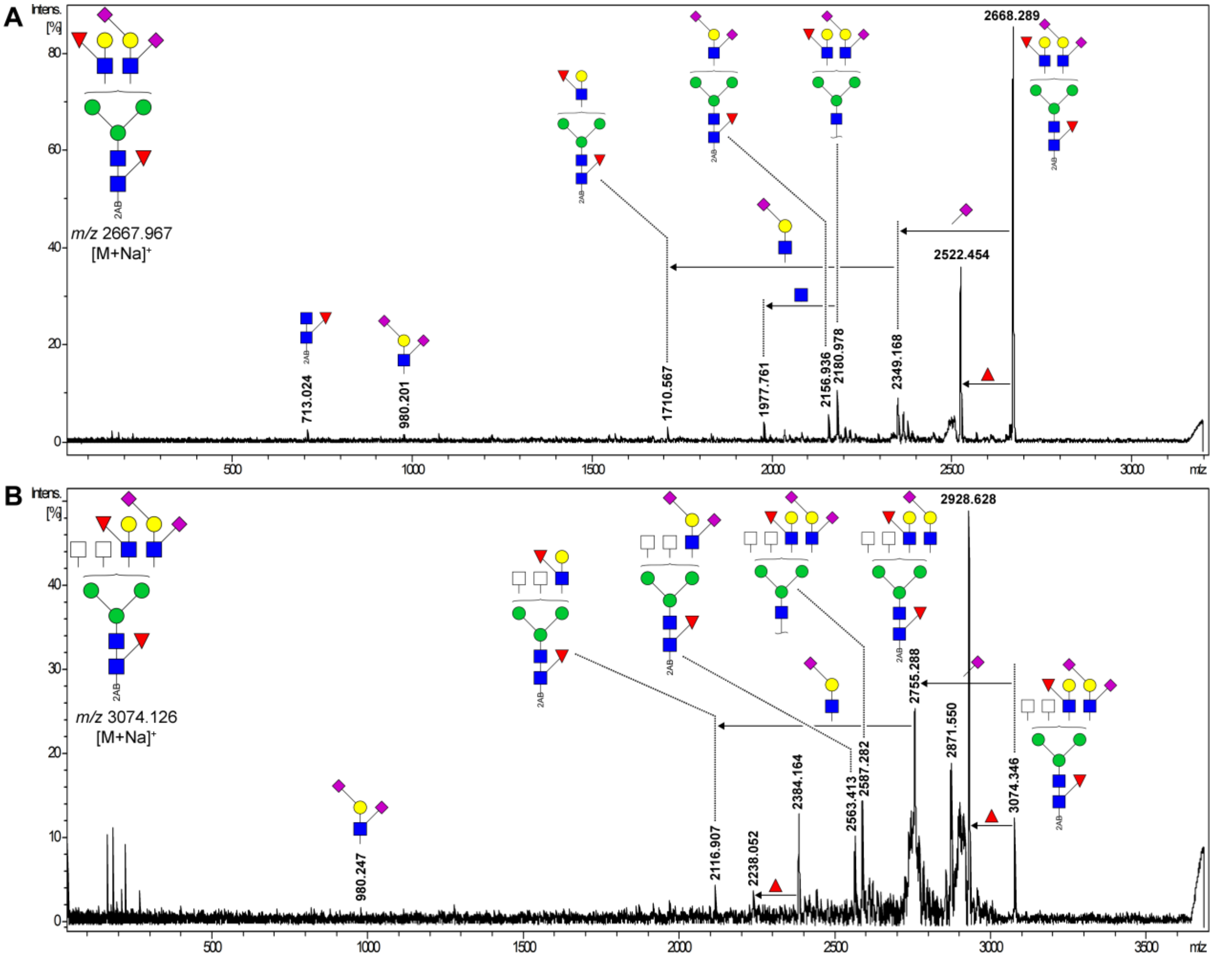


**Fig. S4. MS/MS spectra of N-glycans carrying the di-sialyl Lewis C epitope found in the mammalian brain.** The proposed structure of the precursor N-glycan and the theoretical mass of the sodiated ion are shown in the upper left corner. **(A)** Annotated MS/MS spectrum of an N-glycan ion (m/z 2668.289) detected in the human dorsolateral prefrontal cortex. This N-glycan was detected in chromatographic peak 37 and has the monosaccharide composition H5N4F2E1L1. Diagnostic fragments for core fucose (m/z 713.024, fucosylated chitobiose core labelled with 2-AB) and the di-sialyl Lewis C epitope (m/z 980.201, an esterified and a lactonized Neu5Ac on the same antenna) are observed. Other isobaric configurations, such as an Neu5Acα(2-8)- Neu5Acα(2-6)-Gal disialic linkage, are excluded by the presence of the fragment at m/z 2349.168 which is formed by the loss of a single esterified Neu5Ac – this would not be possible if the α(2-6)-linked Neu5Ac was further substituted with a lactonized Neu5Ac. It is known that antennary fucosylation does not occur on the antenna carrying the di-sialyl Lewis C epitope (*61*). **(B)** Annotated MS/MS spectrum of an N-glycan ion (m/z 3074.346) detected in the rat striatum. This N-glycan was detected in chromatographic peak 38 and has the monosaccharide composition H5N6F2E1L1. A diagnostic fragment for the di-sialyl Lewis C epitope is observed at m/z 980.247. As in (A), a disialic linkage is excluded by the presence of the fragment at m/z 2755.288 which is formed by the loss of a single esterified. x-axis – m/z; y-axis – Percentage of maximum intensity. H – hexose; N – N-acetylhexosamine; F - deoxyhexose; E – esterified Neu5Ac; L – lactonized Neu5Ac. Spectra were processed using the ‘Baseline Subtraction’ and ‘Smoothing’ functions available in the Compass DataAnalysis™ software package (Bruker Daltonics).

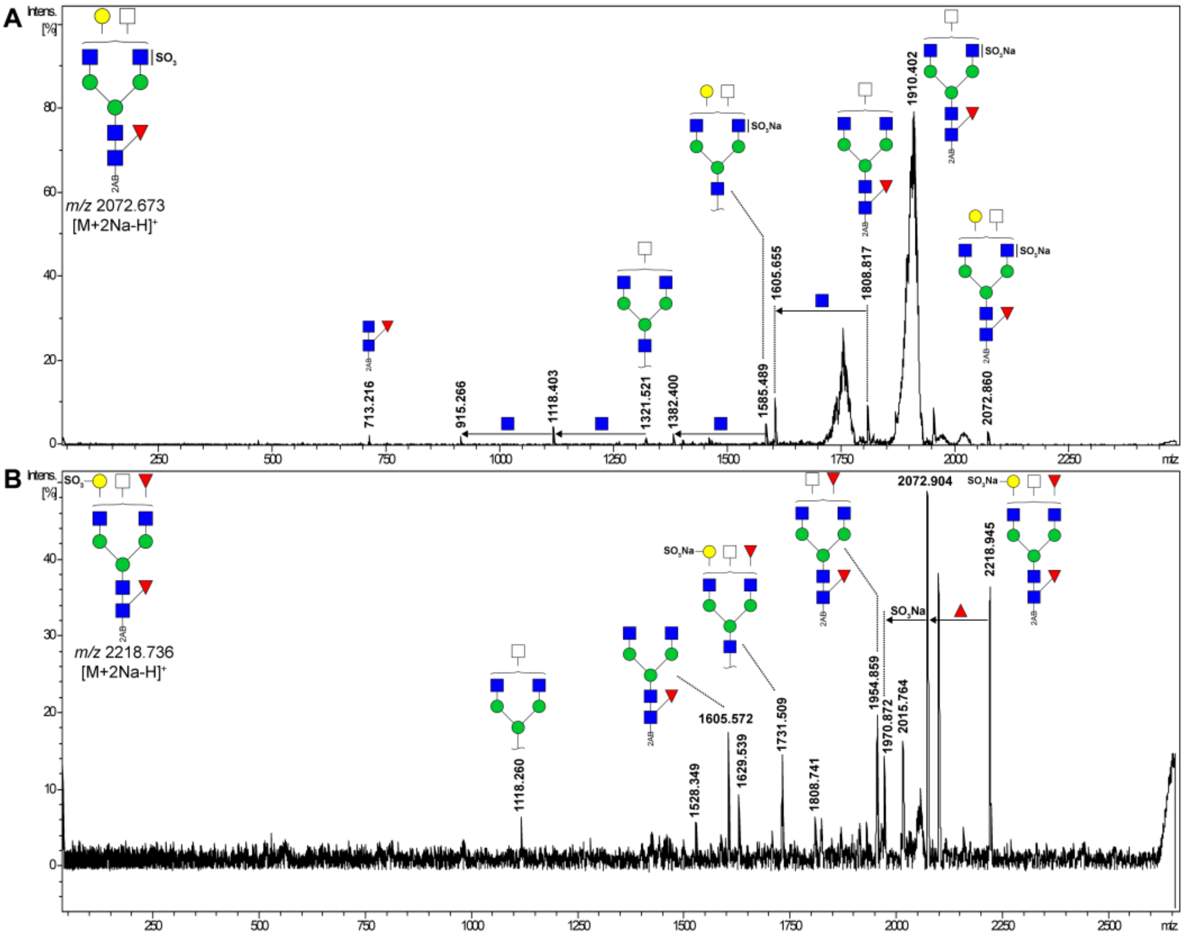


**Fig. S5. MS/MS spectra of sulfated N-glycans found in the mammalian brain.** The proposed composition of the precursor N-glycan and the theoretical mass of the sodiated ion are shown in the upper left corner. **(A)** Annotated MS/MS spectrum of an N-glycan ion (m/z 2072.860) detected in the rat hippocampus. This N-glycan was detected in chromatographic peak 10. The fragmentation pattern shows that the N-glycan is of the complex type (H4N5F1) and is modified by a sodiated phosphate group ($H^+ \rightarrow Na^+$ neutral exchange). The presence of a diagnostic fragment at m/z 1910.402 that lacks a hexose but has retained the sulfate group indicates that the sulfate group is attached to an antennary N-acetylglucosamine. A diagnostic fragment for core fucose is observed at m/z 713.216 (fucosylated chitobiose core labelled with 2-AB). **(B)** Annotated MS/MS spectrum of an N-glycan ion (m/z 2218.945) detected in the human striatum. This N-glycan was detected in chromatographic peak 16 but was not reported in Table

S3 since the intensity of the signal was <10% of the base peak intensity. The fragmentation pattern shows that the N-glycan is of the complex type (H4N5F2) and is modified by a sodiated phosphate group ($H^+ \rightarrow Na^+$ neutral exchange). The presence of a fragment at m/z 1954.859 that is missing both a hexose and the sulfate group suggests that the sulfate group is attached to an antennary galactose while the presence of a fragment at m/z 1731.509 that is missing the reducing end GlcNAc and a fucose suggests that the N-glycan is core fucosylated. x-axis – m/z; y-axis - Percentage of maximum intensity. H – hexose; N – N-acetylhexosamine; F - deoxyhexose. Spectra were processed using the ‘Baseline Subtraction’ and ‘Smoothing’ functions available in the Compass DataAnalysis™ software package (Bruker Daltonics).

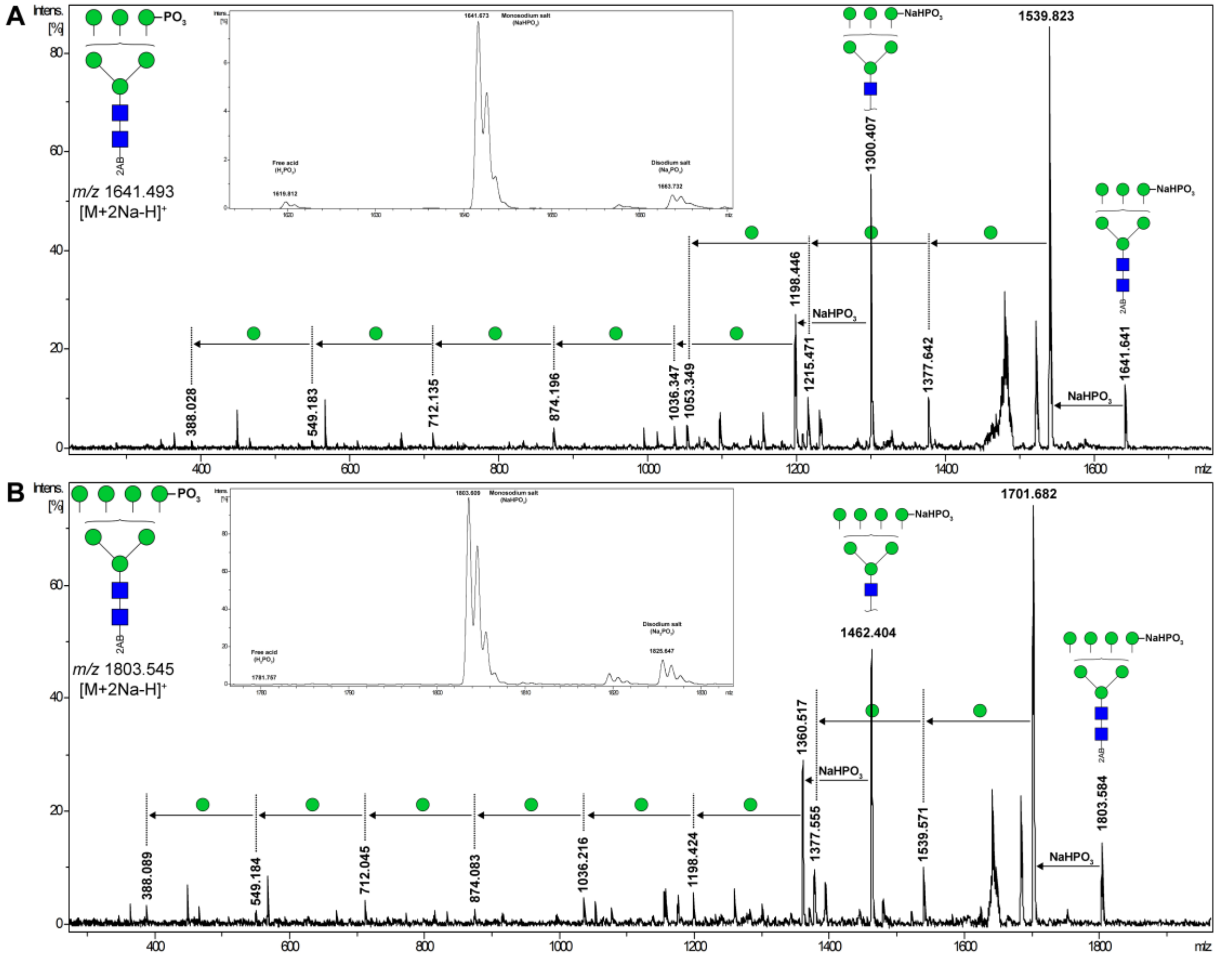


**Fig. S6. MS/MS spectra of phosphorylated N-glycans found in the human brain.** The proposed composition of the precursor N-glycan and the theoretical mass of the sodiated ion are shown in the upper left corner. **(A)** Annotated MS/MS spectrum of an N-glycan ion (m/z 1641.641) detected in the human striatum. This N-glycan was detected in chromatographic peak 19 but was not reported in Table S3 since the intensity of the signal was <10% of the base peak intensity. The fragmentation pattern shows that the N-glycan is of the oligomannose type (M6) and is modified by a mono-sodiated phosphate group ($H^+ \rightarrow Na^+$ neutral exchange). The inset shows a portion of the MS spectrum from which the MS/MS spectrum was obtained. The presence of signals corresponding to the free acid (m/z 1619.812) as well as the disodium salt (m/z 1663.732) confirm that the precursor ion contains a phosphate group rather than a sulfate group (*74*). **(B)** Annotated MS/MS spectrum of an N-glycan ion (m/z 1803.584) detected in the human striatum. This N-glycan was detected in chromatographic peak 27. The fragmentation pattern shows that the N-glycan is of the oligomannose type (M7) and is modified by a mono-sodiated phosphate group ($H^+$→$Na^+$ neutral exchange). The inset shows a portion of the MS spectrum from which the MS/MS spectrum was obtained. The presence of signals corresponding to the free acid (m/z 1781.757, not visible at this magnification) as well as the disodium salt (m/z 1825.647) confirm that the precursor ion contains a phosphate group rather than a sulfate group (*74*). x-axis – m/z; y-axis – Percentage of maximum intensity. Spectra were processed using the ‘Baseline Subtraction’ and ‘Smoothing’ functions available in the Compass DataAnalysis™ software package (Bruker Daltonics).

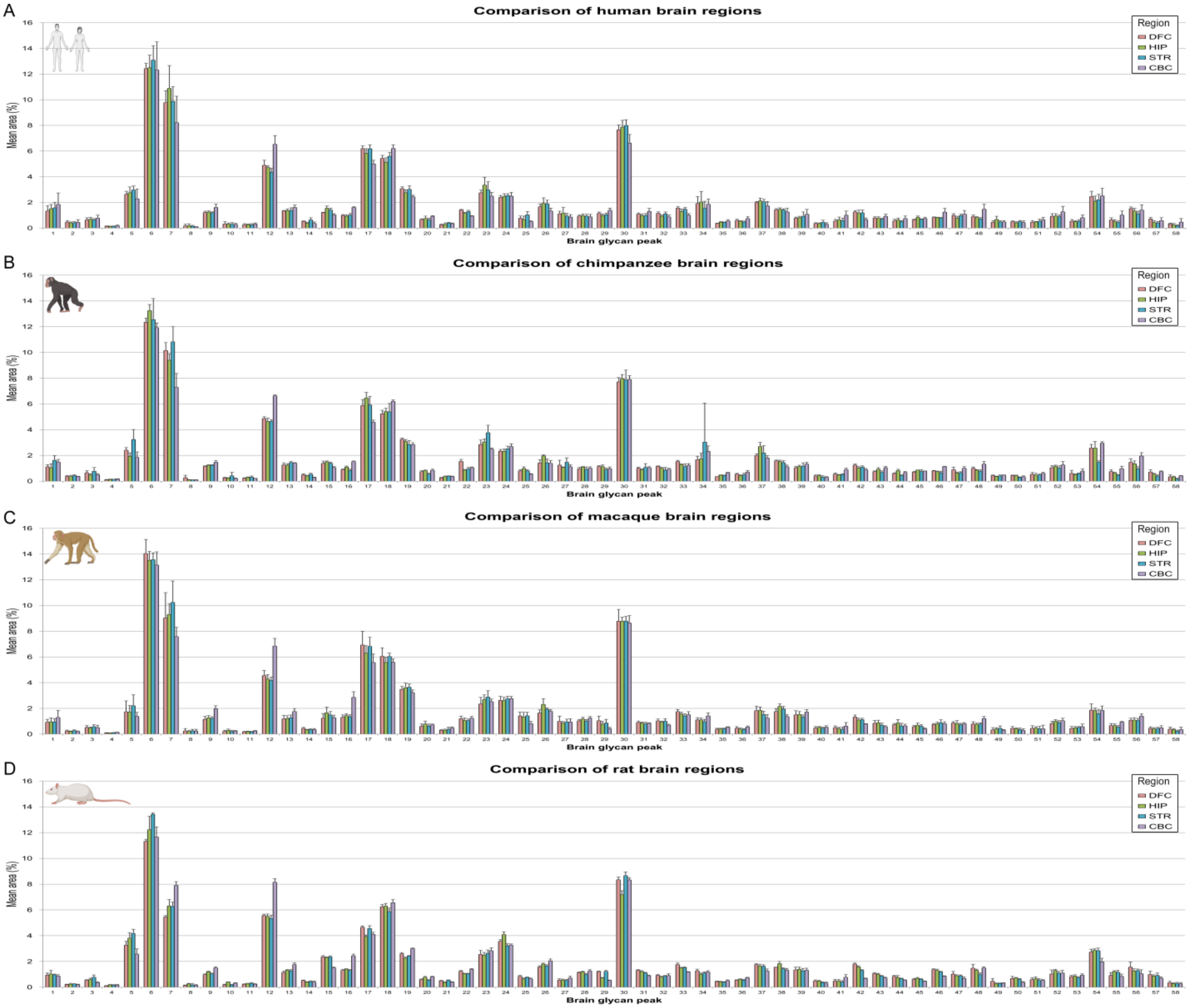


**Fig. S7. Intra-species comparison of N-glycan peak abundance in different brain regions.** Normalized mean relative abundance of each N-glycan peak in the N-glycome of the **(A)** human, **(B)** chimpanzee, **(C)** macaque, and **(D)** rat brain. Abundance is expressed as a percentage of the total integrated chromatographic area. DFC – dorsolateral prefrontal cortex; HIP – hippocampus; STR – striatum; CBC – cerebellar cortex. Error bars show the standard deviation. For statistically significant differences in peak area across brain regions see Table S5.

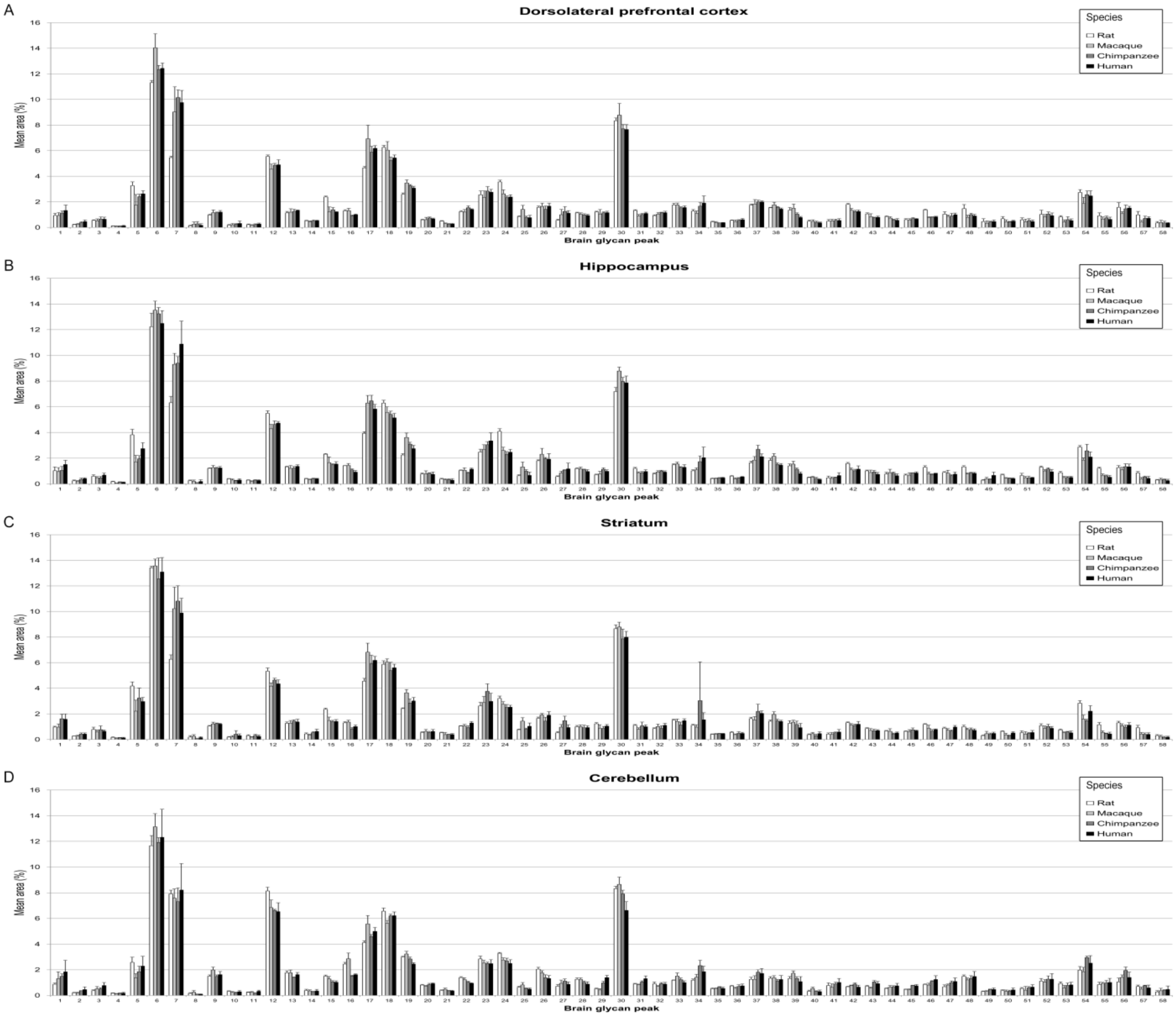


**Fig. S8. Inter-species comparison of the abundance of individual N-glycan peaks across homologous brain regions.** Normalized mean relative abundance of each N-glycan peak in the N-glycome of the **(A)** dorsolateral prefrontal cortex, **(B)** hippocampus, **(C)** striatum, and **(D)** cerebellar cortex. Abundance is expressed as a percentage of the total integrated chromatographic area. Error bars show the standard deviation. For statistically significant differences in peak area between species see Table S6.

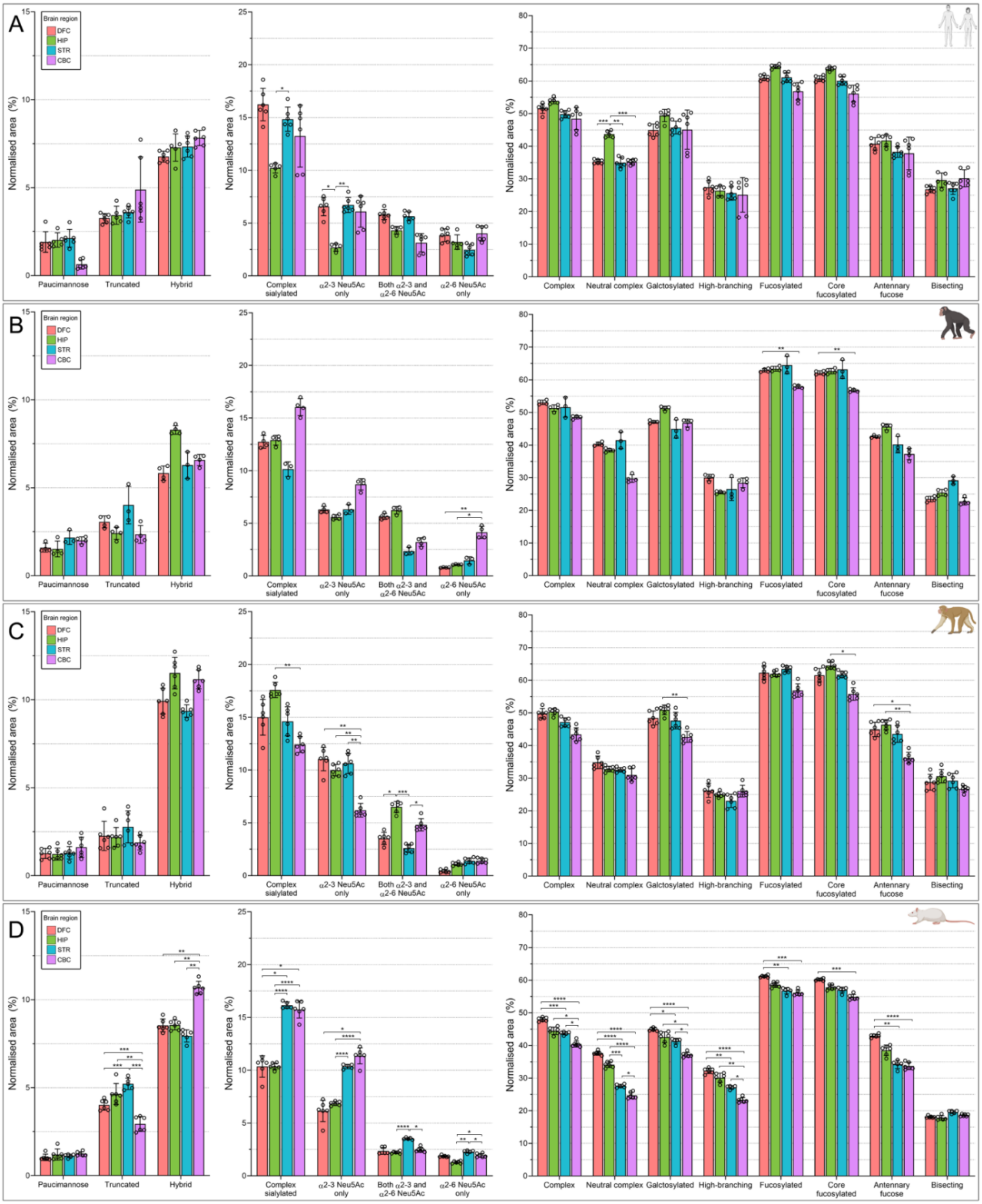


**Fig. S9. Intra-species comparison of N-glycan derived traits in different brain regions.** Relative abundance of each derived trait in the N-glycome of the **(A)** human, **(B)** chimpanzee, **(C)** macaque, and **(D)** rat brain. Abundance is expressed as the summed area of all chromatographic peaks in which the dominant N-glycan structure exhibits that particular trait as a percentage of the total chromatographic area (see Table S7 for derived trait formulas). Statistical significance between brain regions was tested using ANCOVA with correction for multiple testing using the Bonferroni procedure (Table S8). Adjusted P-values: * 0.05 > P > 0.01, ** 0.01 > P 0.001, *** 0.001 P > 0.0001, **** P < 0.0001. DFC – dorsolateral prefrontal cortex; HIP – hippocampus; STR – striatum; CBC – cerebellar cortex. Error bars show the standard deviation.

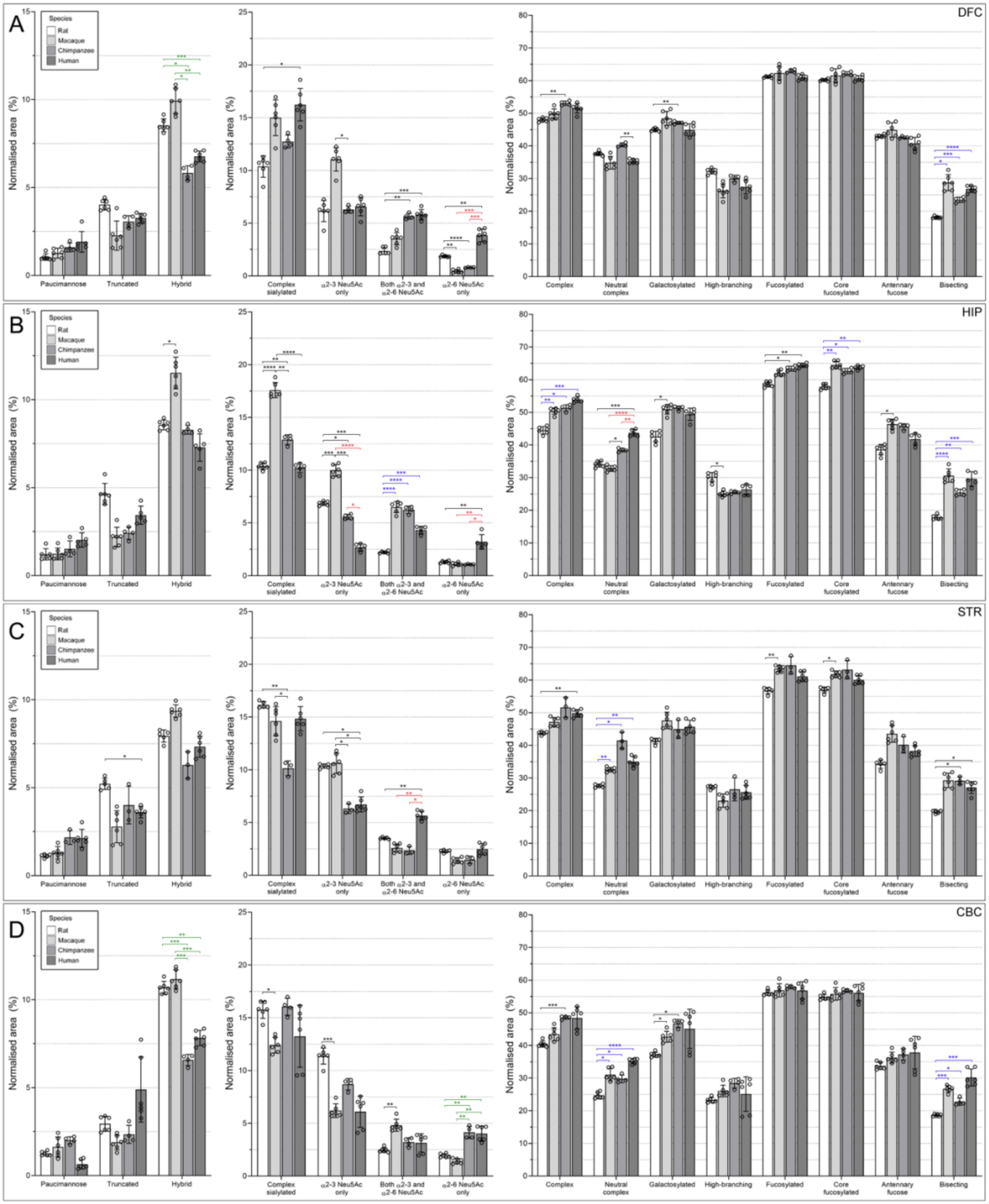


**Fig. S10. Inter-species comparison of the relative abundance of N-glycan derived traits in the brain N-glycome by region.** Relative abundance of each derived trait in the brain N-glycome of the **(A)** dorsolateral prefrontal cortex, **(B)** hippocampus, **(C)** striatum, and **(D)**

cerebellar cortex. Abundance is expressed as the summed area of all chromatographic peaks in which the dominant N-glycan structure exhibits that particular trait as a percentage of the total chromatographic area (see Table S7 for derived trait formulas). Statistical significance between species was tested using ANCOVA with correction for multiple testing using the Bonferroni procedure (Table S8). Adjusted P-values: * $0.05 > P > 0.01$, ** $0.01 > P\ 0.001$, *** $0.001\ P > 0.0001$, **** $P < 0.0001$. Statistically significant phylogenetic differences specific to certain taxa are color coded for clarity: blue – rats vs all primates; green – hominids vs non-hominids; red – humans vs both non-human primates (i.e. chimpanzees and macaques).

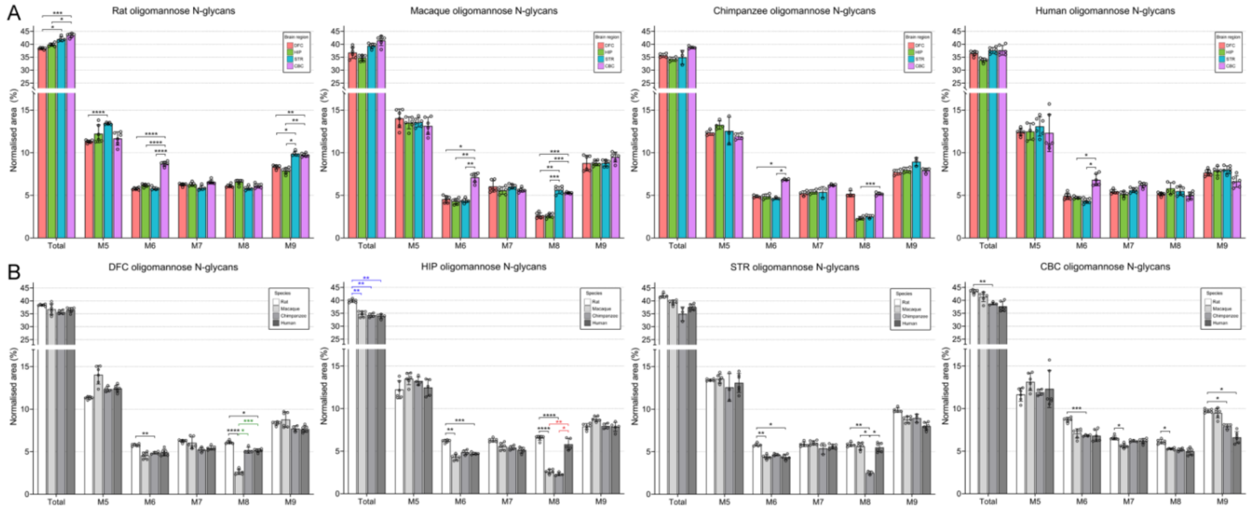


**Fig. S11. Breakdown of oligomannose N-glycan abundance in the brain N-glycome by species and brain regions.** Total oligomannose N-glycan content, as well as the abundance of each of the oligomannose subtypes, is presented as **(A)** an inter-regional comparison within each species, and **(B)** an inter-species evolutionary comparison for each brain region. In addition to all of the subtypes displayed, total oligomannose content also includes the core fucosylated M5 structure (FM5) and the M9 structure with an additional glucose residue (M9+Glc). Statistical significance between groups was tested using ANCOVA with correction for multiple testing using the Bonferroni procedure (Table S8). Adjusted P-values: * 0.05 > P > 0.01, ** 0.01 > P 0.001, *** 0.001 P > 0.0001, **** P < 0.0001. Statistically significant phylogenetic differences specific to certain taxa are color coded for clarity: blue – rats vs all primates; green - hominids vs non-hominids; red – humans vs both non-human primates (i.e. chimpanzees and macaques). DFC – dorsolateral prefrontal cortex; HIP – hippocampus; STR – striatum; CBC – cerebellar cortex.

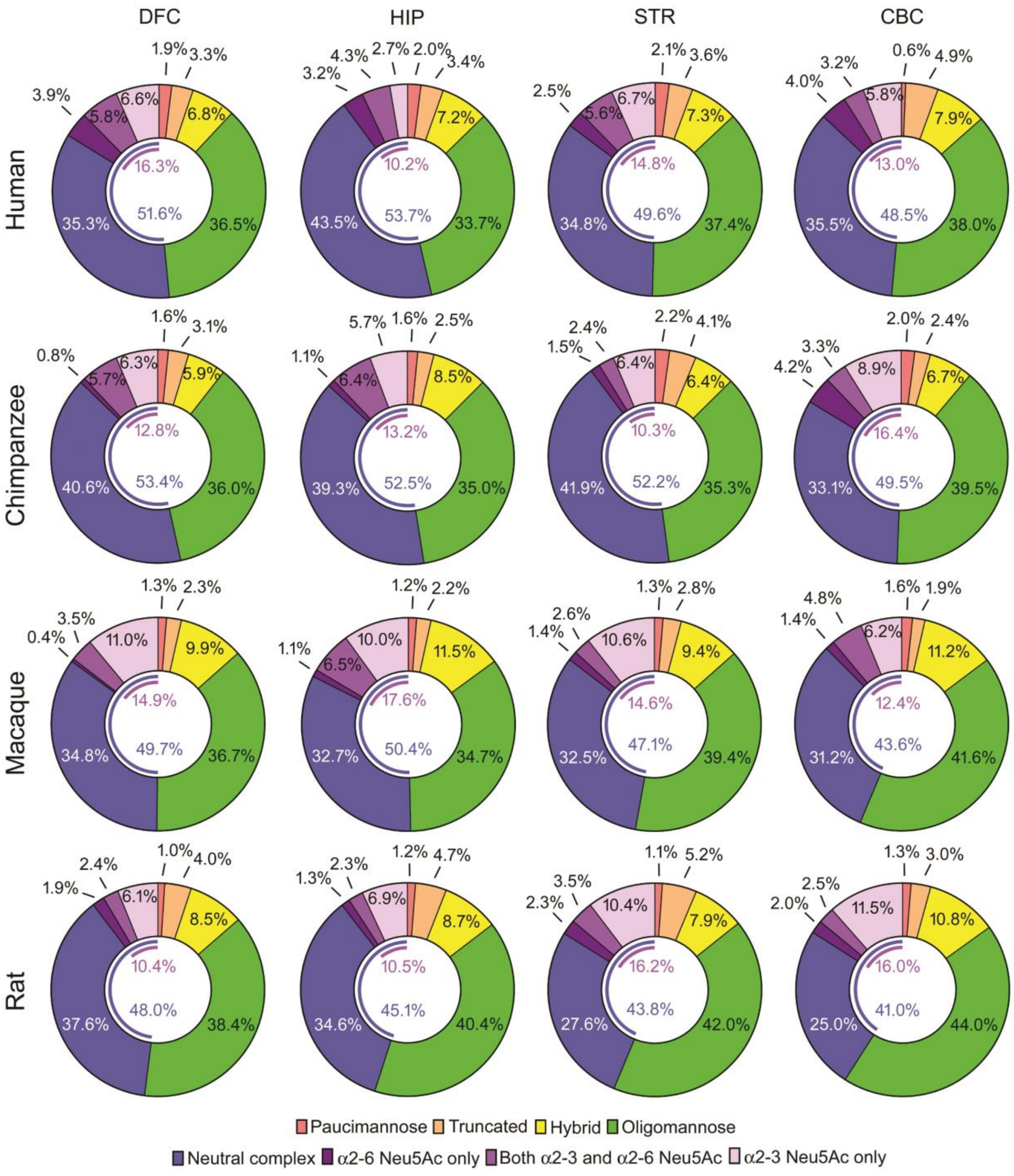


**Fig. S12. Total N-glycome composition by N-glycan type for each group.** Donut charts are aligned as a matrix with rows defining the species and columns defining the brain region. Derived traits were calculated from UPLC N-glycan peak abundance data and were used to determine the abundance of each category of N-glycans within the total N-glycome (expressed as a percentage of the total N-glycome by chromatographic area). Mean values for each group are displayed. The blue arc in the donut hole indicates the percentage of total complex N-glycans (i.e. the sum of both neutral and sialylated complex N-glycans) while the purple arc indicates the percentage of total sialylated complex N-glycans (i.e. the sum of all complex N-glycans containing Neu5Ac). DFC – dorsolateral prefrontal cortex; HIP – hippocampus; STR – striatum; CBC – cerebellar cortex.

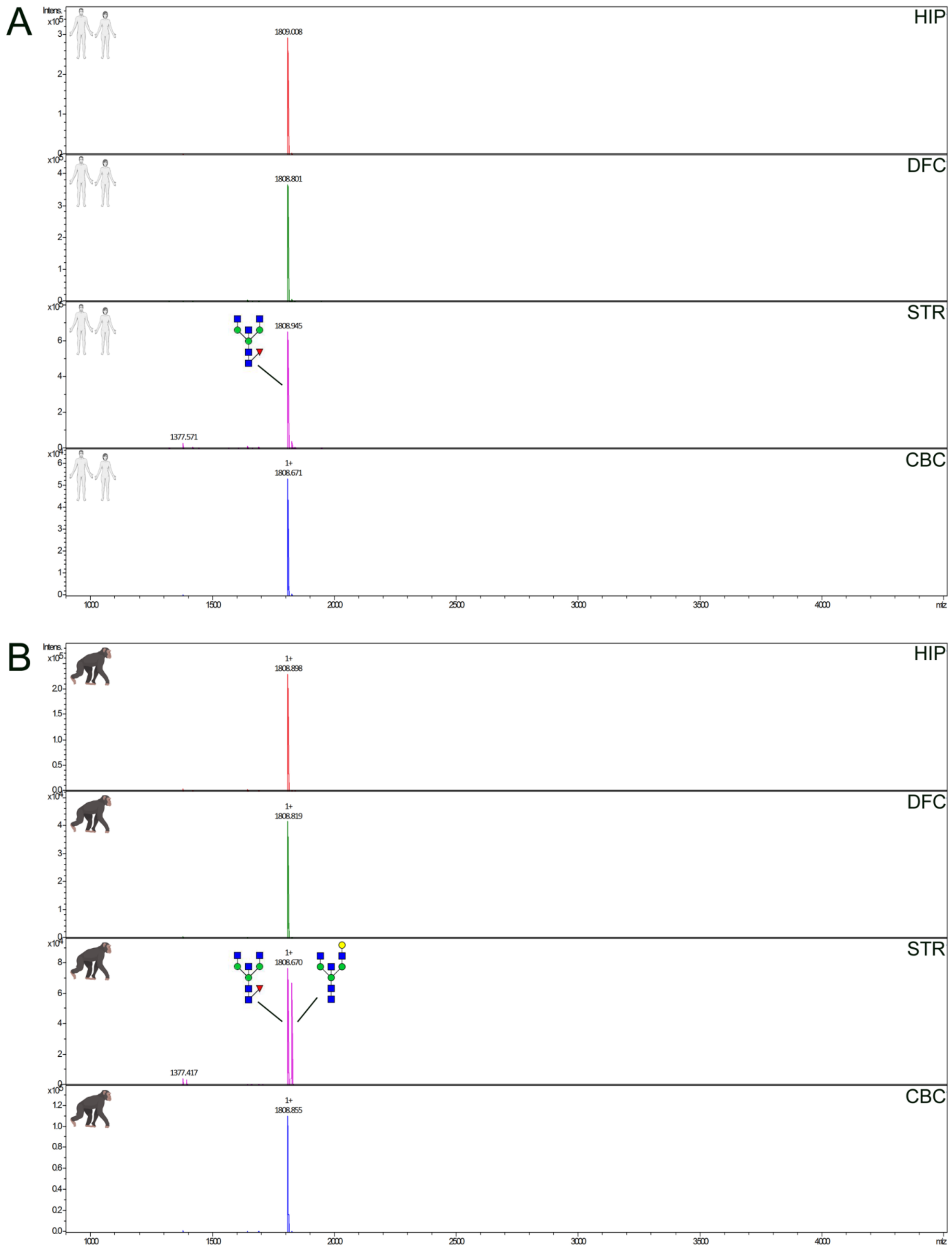


**Fig. S13. MS spectra illustrating region- and species-specific qualitative differences present in equivalent chromatographic peaks.** Although the N-glycan content was generally consistent across corresponding chromatographic peaks in different species and brain regions, this was not always the case. **(A)** MS spectra of chromatographic peak 7 for each of the human brain regions. **(B)** MS spectra for the corresponding N-glycan peak in the chimpanzee brain regions. The dominant ion in all instances can be attributed to the complex N-glycan with monosaccharide composition H3N5F1 (theoretical mass $[M{:}2\text{-}AB+Na^{+}]^{+}$ = 1808.682 Da). However, the chimpanzee striatum additionally contains a secondary N-glycan with the monosaccharide composition H4N5 (theoretical mass $[M{:}2\text{-}AB+Na^{+}]^{+}$ = 1824.677 Da), that is not present in the other brain regions, nor in any of the human brain regions. This N-glycan was also detected in the macaque striatum as a minor structure, but was not detected in the rat brain (data not shown). x-axis – m/z; z-axis – Absolute intensity. HIP – hippocampus; DFC – dorsolateral prefrontal cortex; STR – striatum; CBC – cerebellar cortex. H – hexose; N – N-acetylhexosamine; F - deoxyhexose.

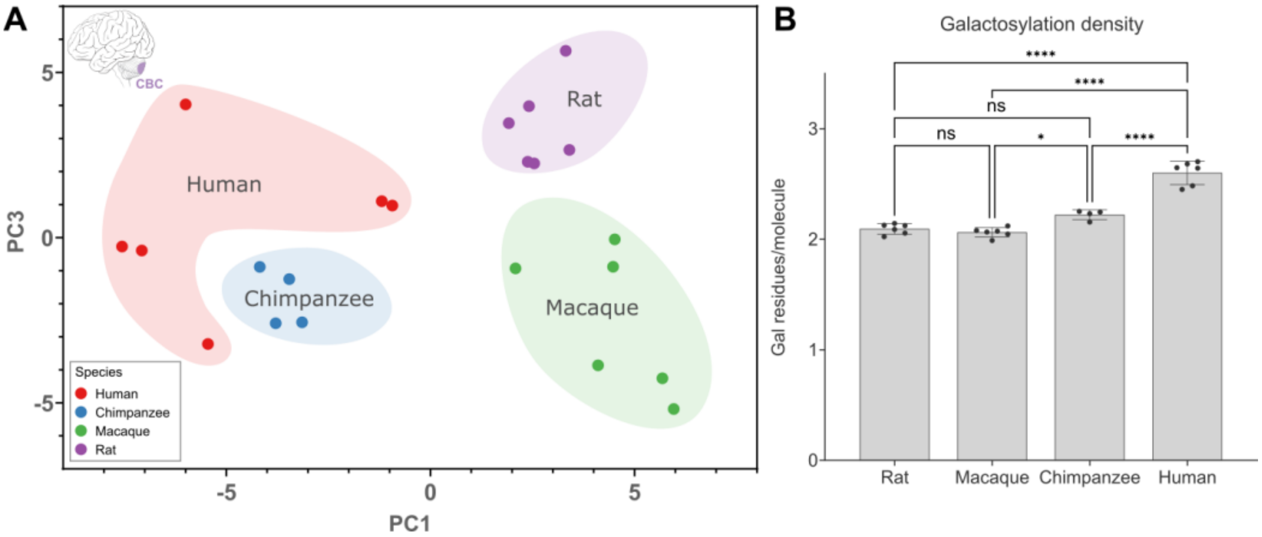


**Fig. S14. Phylogenetic divergence and human-specific features of the CBC N-glycome. (A)** PCA scatter plot of Euarchontoglires CBC total N-glycomes. Clustering according to species is evident indicating taxon-specific differences in N-glycosylation of CBC glycoproteins. The x-axis and y-axis display the first (PC1) and third (PC3) principal components, respectively. **(B)** The human CBC has the highest galactosylation density (expressed as the mean number of galactose residues per galactosylated N-glycan) of all species. Statistical significance between groups was tested using one-way ANOVA with correction for multiple testing using the Bonferroni procedure. Adjusted P-values: * $0.05 > P > 0.01$, **** $P < 0.0001$. Other taxon-specific differences in CBC N-glycosylation can be seen in Fig S10D.

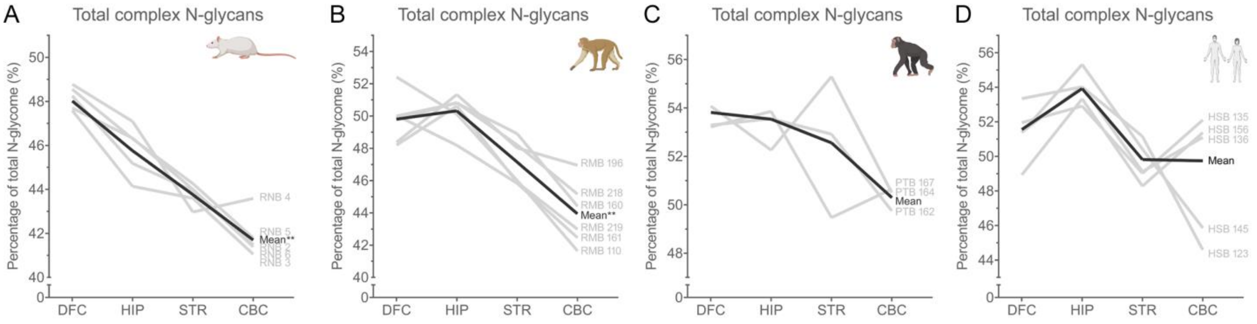


**Fig. S15. Anatomical variation in complex N-glycan abundance in various Euarchontoglires brain N-glycomes.** Total complex N-glycan content in the **(A)** rat, **(B)** macaque, **(C)** chimpanzee, and **(D)** human brain N-glycome according to brain region. Gray lines show the complex N-glycan content across different brain regions of individual animals or donors (labels shown on the right). The black line shows the group mean. Only subjects with data for all brain regions are plotted. Data were analyzed using mixed model analysis and a test for linear trend across brain regions (in the order shown from left to right) was performed with correction for multiple testing using the Bonferroni procedure. Adjusted P-values: ** P < 0.01. DFC – dorsolateral prefrontal cortex; HIP – hippocampus; STR – striatum; CBC – cerebellar cortex.

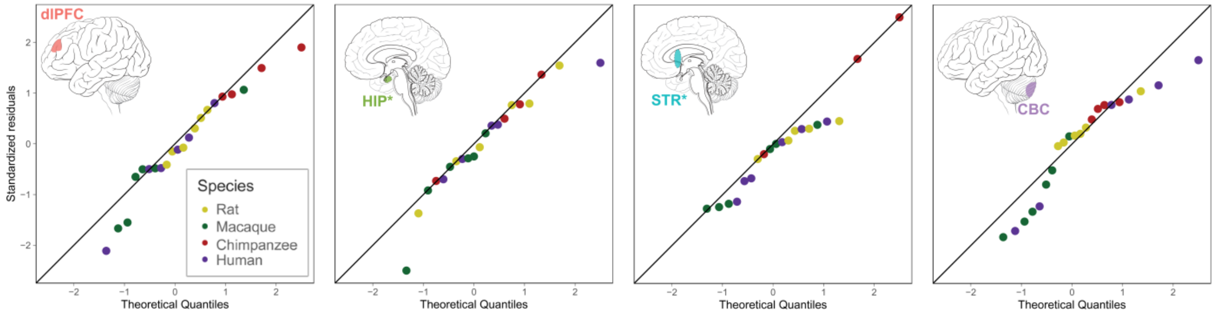


**Fig. S16. The evolutionary change in Euarchontoglires brain N-glycome complexity has been steady.** A quantile-quantile (QQ) plot shows the standardized residuals of the actual abundance of complex N-glycans (y-axis) against the theoretical Gaussian quantiles (x-axis) and confirms the data normality across species. This indicates that the rate of change in the abundance of complex N-glycans did not deviate from the rate that would be expected given the evolutionary distance between species and, therefore, that this change has been gradual without major jumps in any taxa. dlPFC – dorsolateral prefrontal cortex; HIP – hippocampus; STR - striatum; CBC – cerebellar cortex. *Denotes that regions are internal and not visible from the view shown.

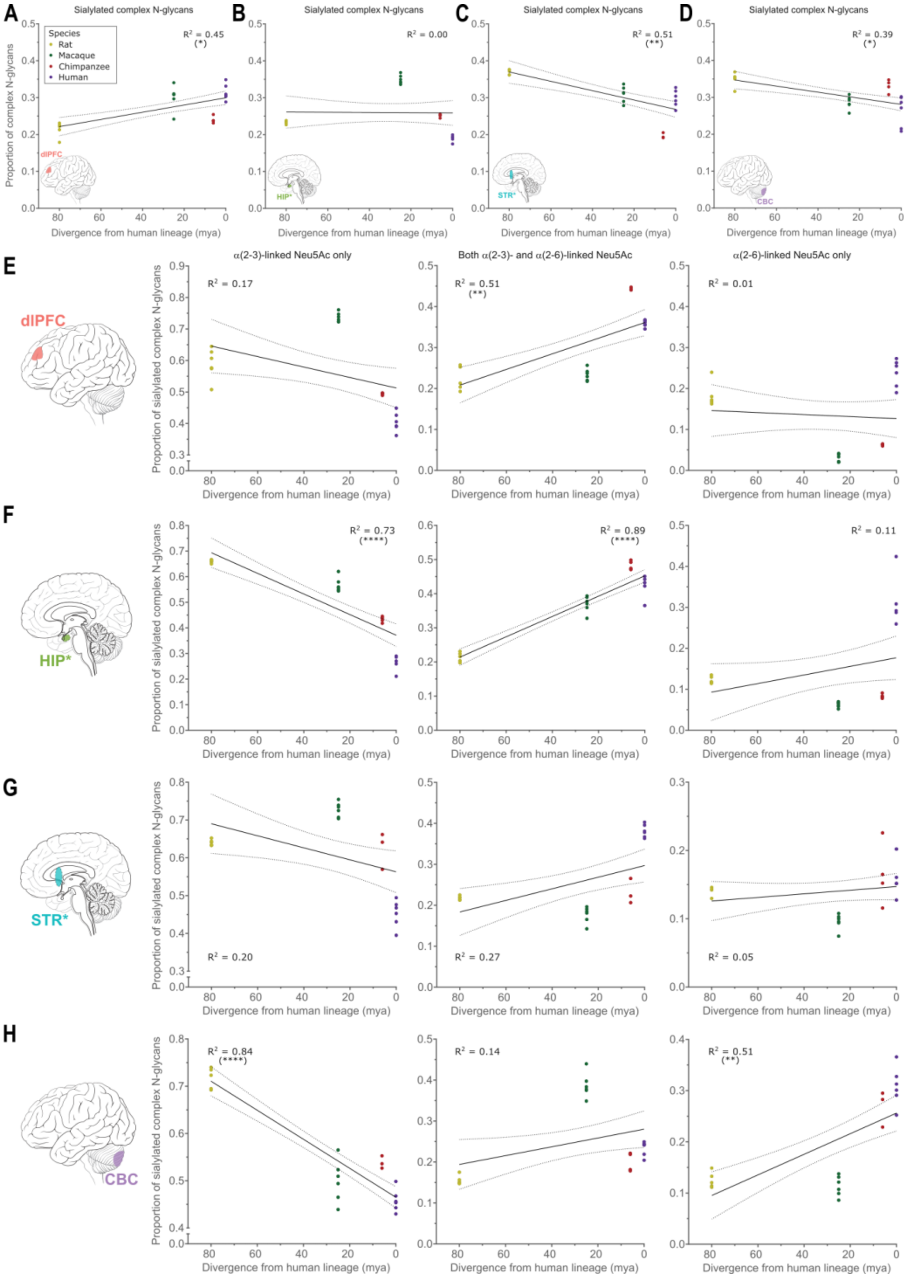


**Fig. S17. Phylogenetic trends in sialylation frequency and linkage distribution.** Sialylation frequency in the (**A**) dlPFC, (**B**) HIP, (**C**) STR, and (**D**) CBC of various Euarchontoglires mammals. Neu5Ac linkage distribution in the (**E**) dlPFC, (**F**) HIP, (**G**) STR, and (**H**) CBC. The plots show the proportion of sialylated complex N-glycans carrying only α(2-3)-linked Neu5Ac, both α(2-3)- and α(2-6)-linked Neu5Ac, and only α(2-6)-linked Neu5Ac from left to right. Data were analyzed using linear regression (90% confidence intervals and $R^2$ values are shown).

Asterisks in parentheses indicate lines whose slopes were significantly non-zero after correction for multiple testing using the Bonferroni procedure. Adjusted P-values: (*) $0.05>P\geq0.01$, (**) $0.01>P\geq0.001$, (****) $P<0.0001$. *Denotes that regions are internal and not visible from the view shown. dlFC – dorsolateral prefrontal cortex; HIP – hippocampus; STR – striatum; CBC – cerebellar cortex; mya – millions of years ago.

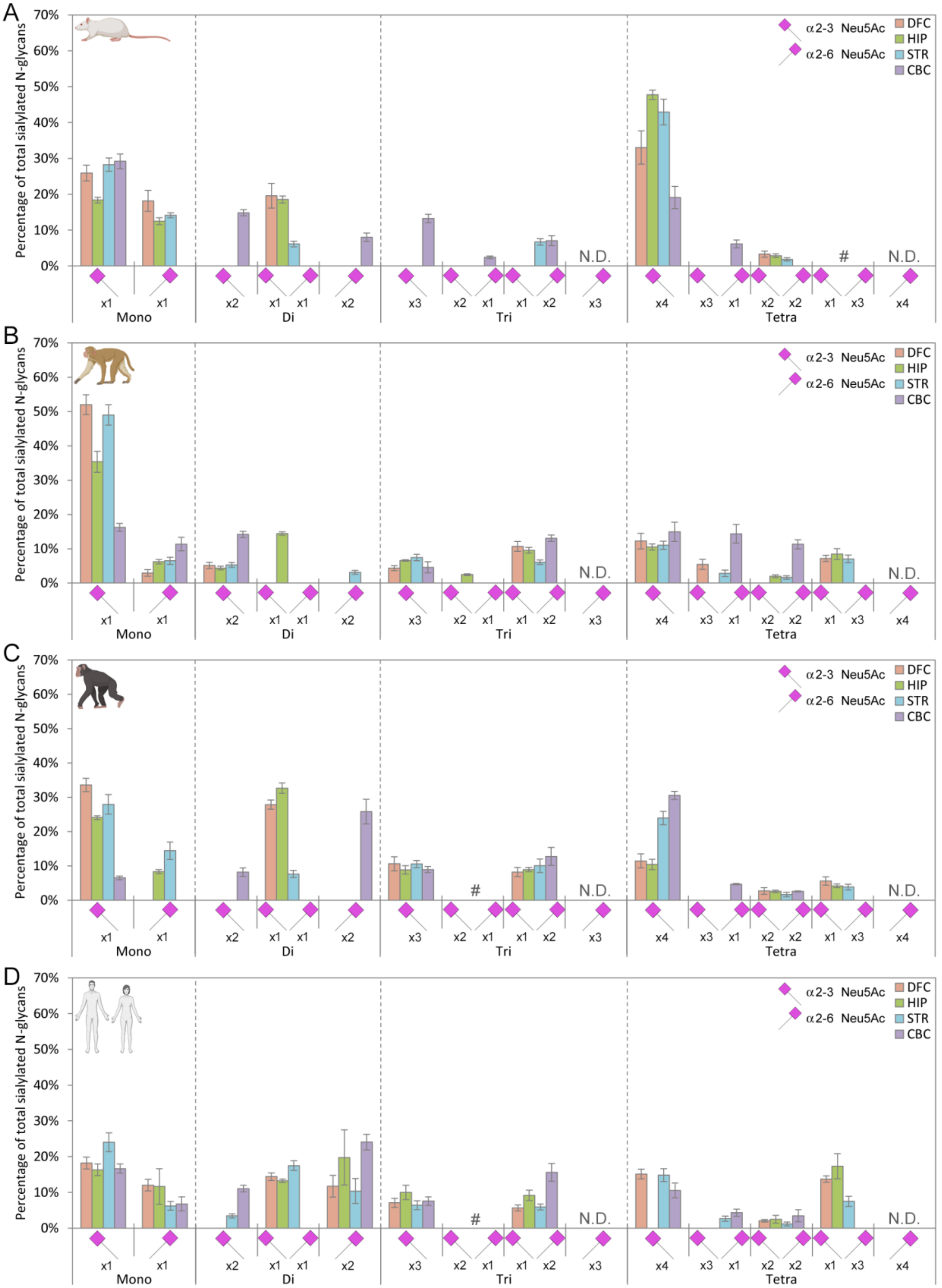
A
B
C
D
Percentage of total sialylated N-glycans
70%
60%
50%
40%
30%
20%
10%
0%
α2-3 Neu5Ac
α2-6 Neu5Ac
DFC
HIP
STR
CBC
N.D.
#
x1
x2
x3
x4
Mono
Di
Tri
Tetra

**Fig. S18. Overview of Neu5Ac linkage on complex N-glycans of the brain in various Euarchontoglires species.** Distribution of α(2-3)- and α(2-6)-linked Neu5Ac on complex N-glycans in the adult **(A)** rat, **(B)** macaque, **(C)** chimpanzee, and **(D)** human brain by region. All possible Neu5Ac linkage combinations are depicted for mono-, di-, tri-, and tetrasialylated complex N-glycans. Error bars show the standard deviation. ND – not detected; # – detected as minor structures. All data are expressed as a percentage of total sialylated complex N-glycans. DFC – dorsolateral prefrontal cortex; HIP – hippocampus; STR – striatum; CBC – cerebellar cortex.

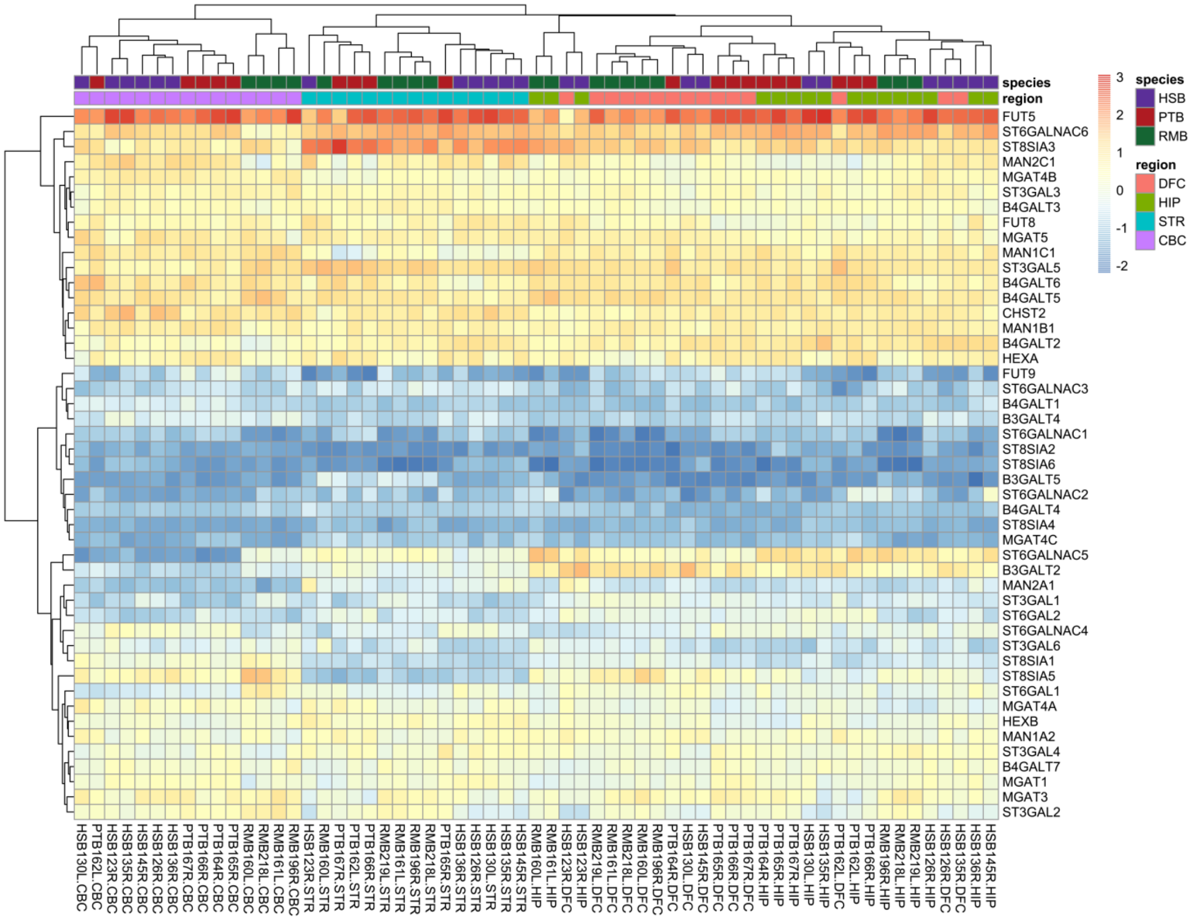


**Fig. S19. Clustering of primate brain samples according to their glycogene expression profiles.** The heatmap shows scaled glycogene expression for all of the primate brain samples. The dendrograms show clustering of samples (top) and glycogenes (left). Notably, clustering of samples is driven primarily by brain region and secondarily by species. Z-score sample-wise normalization was used to eliminate species batch effect in the RNA-Seq expression dataset. Species: HSB – *Homo sapiens* brain; PTB – *Pan troglodytes* brain; RMB – *Macaca mulatta* brain. Brain regions: DFC – dorsolateral prefrontal cortex; HIP – hippocampus; STR – striatum; CBC – cerebellar cortex.

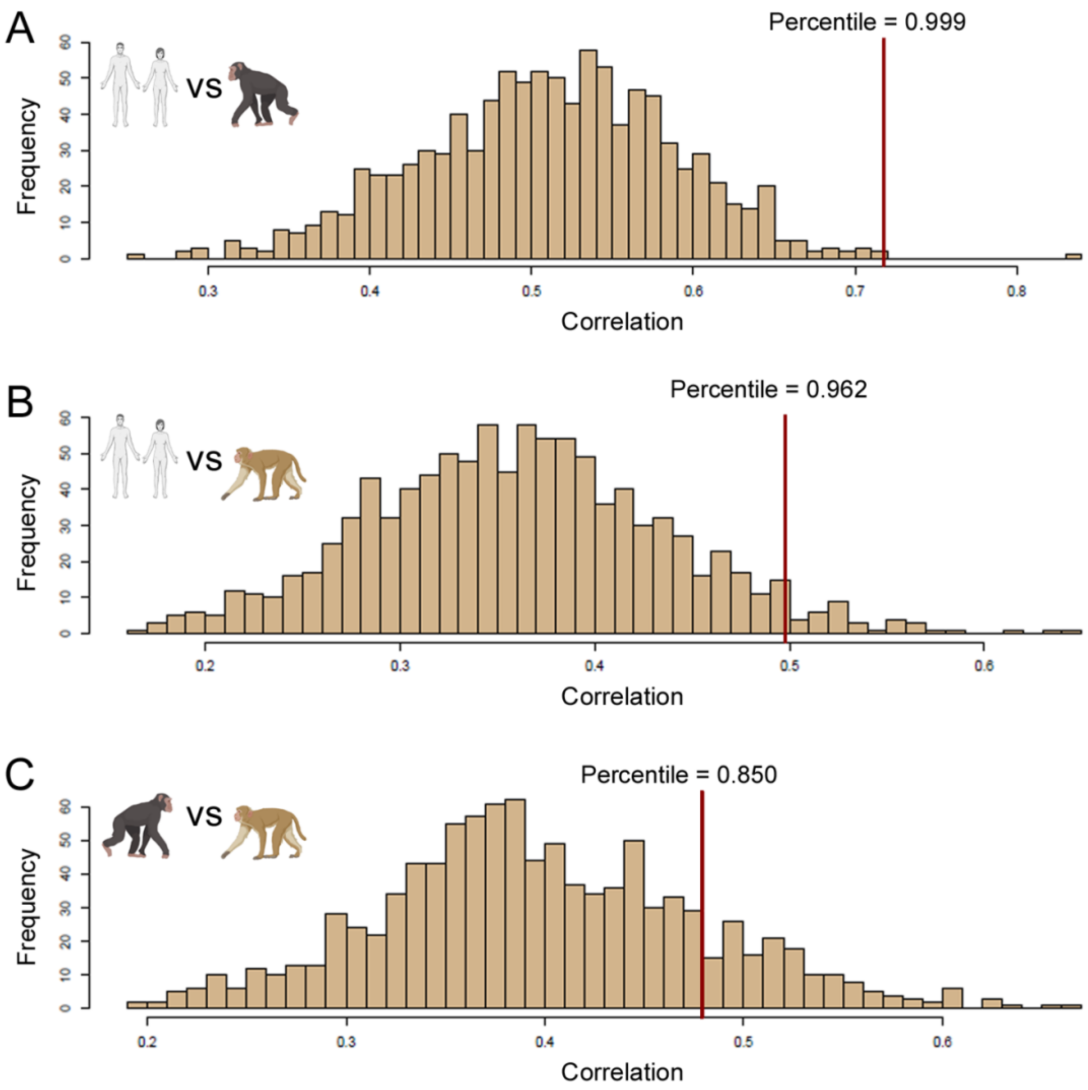


**Fig. S20. Expression of glycogenes is very highly conserved in the primate brain.** Cross-species glycogene correlation was compared with the correlation of 47 genes randomly selected from the respective genomes. Histograms showing the distribution of 1000 resamplings for the **(A)** human vs chimpanzee (glycogene correlation = 0.72; quantile = 0.999), **(B)** human vs macaque (glycogene correlation = 0.50; quantile = 0.962), and **(C)** chimpanzee vs macaque (glycogene correlation = 0.48; quantile = 0.850) comparisons. The x-axis displays correlation while the y-axis shows frequency. The vertical red lines show the position (quantile) of the glycogene correlations.

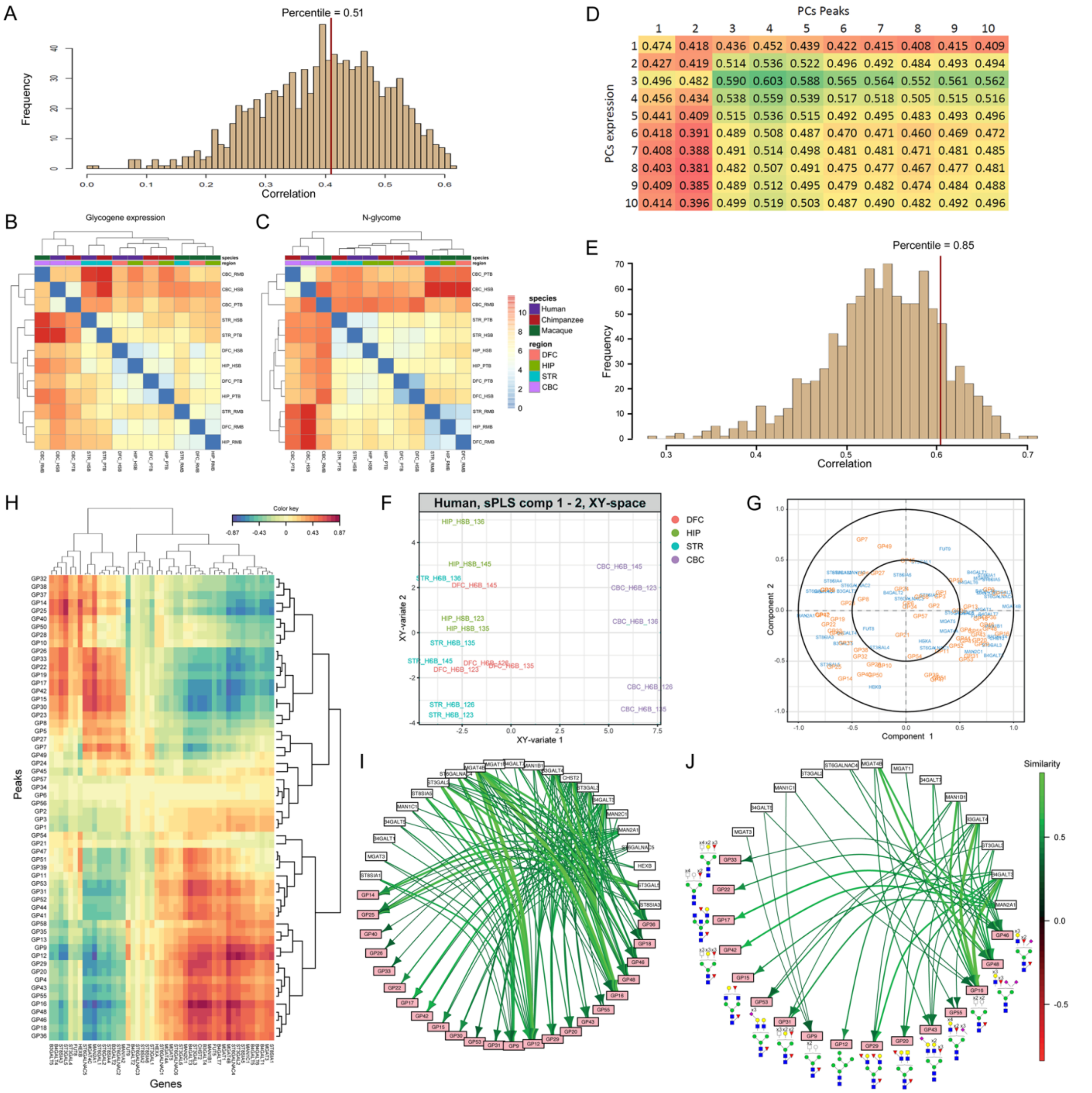


**Fig. S21. Correlation of glycogene expression with N-glycosylation in the primate brain. (A)** Global correlation between the expression of 47 random genes and the N-glycome across all matched samples. Histogram showing the distribution of 1,000 resamplings of 47 random genes. The vertical red line shows the position (quantile) of the glycogene correlation (r=0.42, quantile=0.51). The correlation between each of the primate brain samples was determined with regard to **(B)** glycogene expression, and **(C)** the N-glycome. Following PCA, distance matrices and dendrograms were generated by summarizing the first ten principal components (PCs) by

Euclidean distances of the mean PC values per region and species. The correlation between the two distance matrices, determined using a Mantel correlogram, is r=0.73 (P=0.01 for 100 permutations) when using all brain regions. This correlation value is not significantly higher than would be achieved by resampling a set of 47 random genes 1000 times and is mainly driven by the uniqueness of CBC in terms of both the transcriptome and the glycome (r=0.43 when excluding the CBC, data not shown). **(D)** Optimum PCs selected for correlation: PC3 for glycogenes (rows) and PC4 for N-glycans (columns). **(E)** Global correlation between the expression of 47 random genes and the N-glycome across all matched samples after optimized PCA. Histogram showing the distribution of 1000 resamplings of 47 random genes. The vertical red line shows the position (quantile) of the optimized glycogene correlation (r=0.603, quantile=0.85). **(F)** Glycogenes and N-glycan peaks represented in the average space of the two selected informative sPLS components. **(G)** Correlation circle plot showing the results of sPLS analysis in humans. Glycogenes are written in blue text, N-glycan peaks are written in orange text. **(H)** Heatmap showing the scalar product between pairs of glycogenes and N-glycan peaks representing their correlation structure derived from the sPLS analysis in humans. **(I)** Putative bipartite network of N-glycosylation in the human brain indicating the enzymes predicted to be involved in the synthesis of particular N-glycan peaks. **(J)** Curated network obtained by manual elimination of biologically implausible associations based on common knowledge of N-glycan biosynthetic pathways (*55*) in conjunction with available information regarding enzyme activity (*56*). The major N-glycan structure in each of the peaks is illustrated. Species: HSB – *Homo sapiens* brain; PTB – *Pan troglodytes* brain; RMB – *Macaca mulatta* brain. Brain regions: DFC – dorsolateral prefrontal cortex; HIP – hippocampus; STR – striatum; CBC – cerebellar cortex.

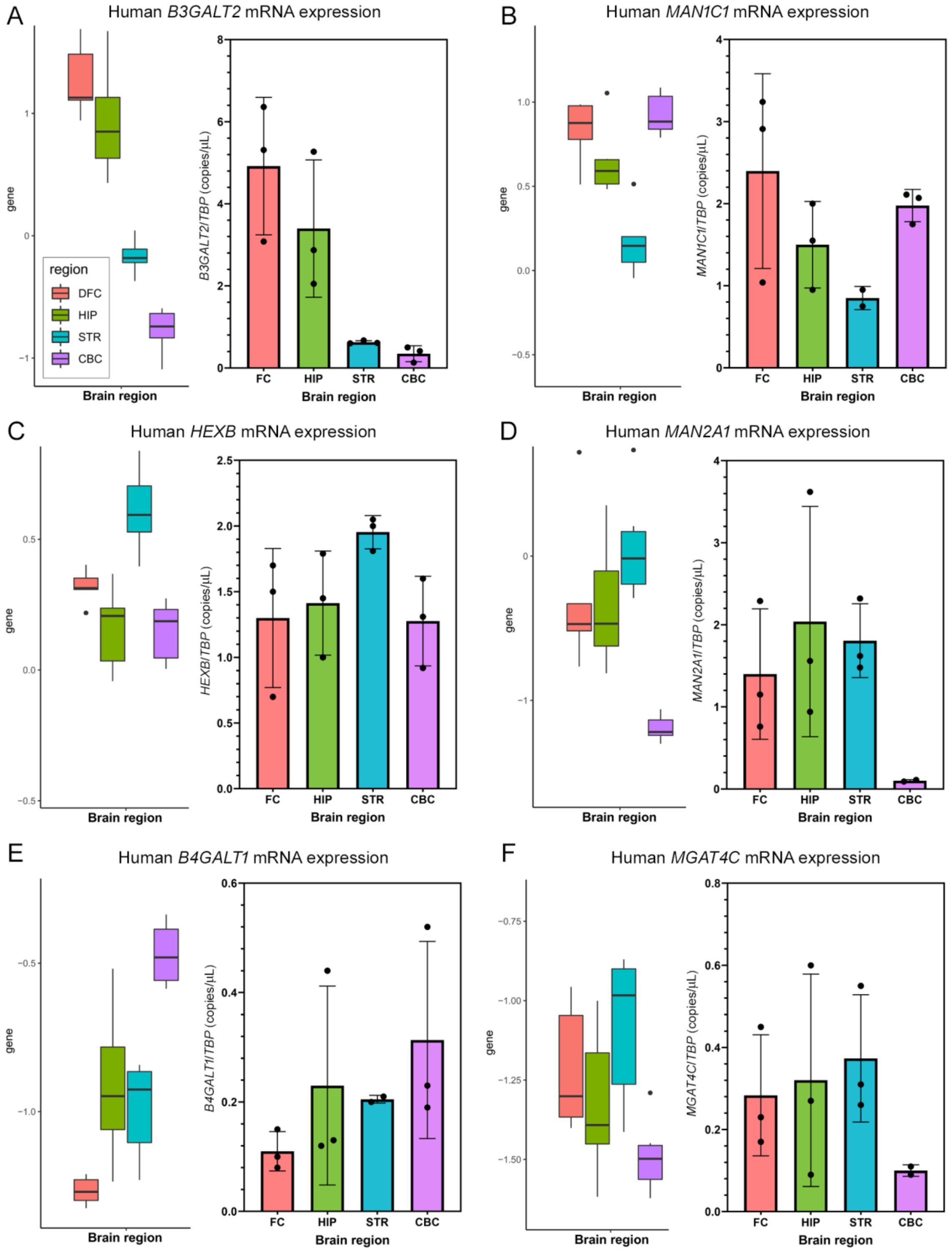


**Fig. S22. Validation of glycogene mRNA expression in the human brain.** Bulk RNA-Seq expression data was validated using ddPCR in each of the four regions of the human brain for the following glycogenes: **(A)** *B3GALT2* (Group I), **(B)** *MAN1C1* (Group III), **(C)** *HEXB* (Group

IV), **(D)** *MAN2A1* (Group IV), **(E)** *B4GALT1* (Group VI), and **(F)** *MGAT4C* (Group V). Left panels show bulk RNA-Seq expression data. The y-axes show scaled expression of logRPKM values within samples and among the 47 glycogenes. Key for region color coding is shown in (A). Right panels show normalized mRNA expression for the corresponding gene as quantified by ddPCR. Expression is normalized to the reference gene *TBP* and expressed as copies of template/μL. Error bars show standard deviation. DFC – dorsolateral prefrontal cortex; FC - frontal cortex; HIP – hippocampus; STR – striatum; CBC – cerebellar cortex.

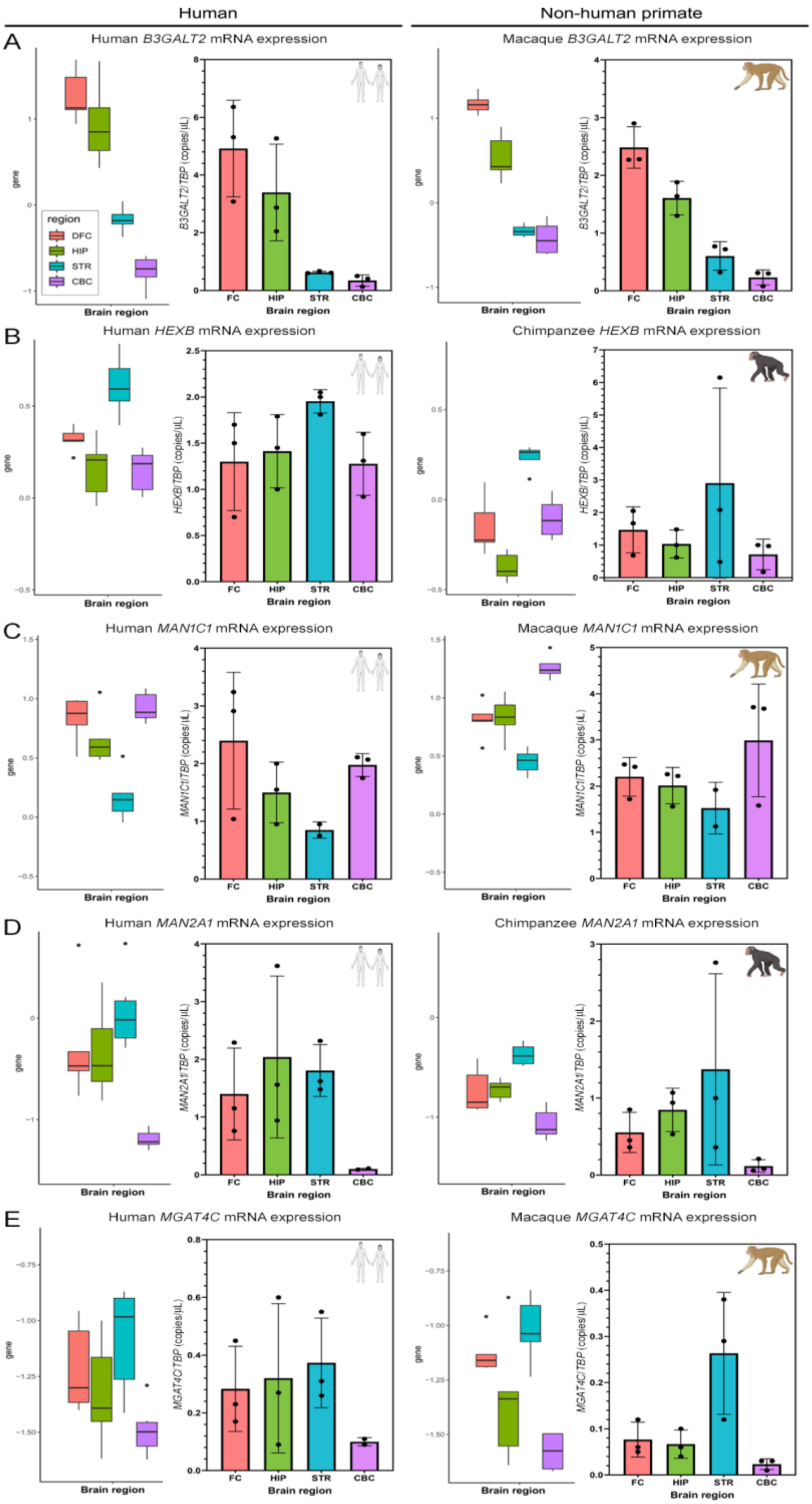

Human
Non-human primate
A
Human B3GALT2 mRNA expression
Macaque B3GALT2 mRNA expression
region
DFC
HIP
STR
CBC
B3GALT2/TBP (copies/uL)
B
Human HEXB mRNA expression
Chimpanzee HEXB mRNA expression
HEXB/TBP (copies/uL)
C
Human MAN1C1 mRNA expression
Macaque MAN1C1 mRNA expression
MAN1C1/TBP (copies/uL)
D
Human MAN2A1 mRNA expression
Chimpanzee MAN2A1 mRNA expression
MAN2A1/TBP (copies/uL)
E
Human MGAT4C mRNA expression
Macaque MGAT4C mRNA expression
MGAT4C/TBP (copies/uL)
gene
FC
HIP
STR
CBC
Brain region

**Fig. S23. Glycogenes exhibiting an evolutionarily conserved spatial expression pattern in the primate brain.** Bulk RNA-Seq expression data was validated using ddPCR in each of the four brain regions for the following glycogenes: **(A)** *B3GALT2* (Group I), **(B)** *HEXB* (Group IV), **(C)** *MAN1C1* (Group III), **(D)** *MAN2A1* (Group V), and **(E)** *MGAT4C* (Group V). The left column shows expression of human glycogenes while the right column shows expression of the orthologous genes in non-human primates (either chimpanzee or macaque). In each column, the left panels show bulk RNA-Seq expression data. The y-axes show scaled expression of logRPKM values within samples and among the 47 glycogenes. Key for region color coding is shown in (A). Right panels show normalized mRNA expression for the corresponding gene as quantified by ddPCR. Expression is normalized to the reference gene *TBP* and expressed as copies of template/µL. Error bars show standard deviation. DFC – dorsolateral prefrontal cortex; FC – frontal cortex; HIP – hippocampus; STR – striatum; CBC – cerebellar cortex.

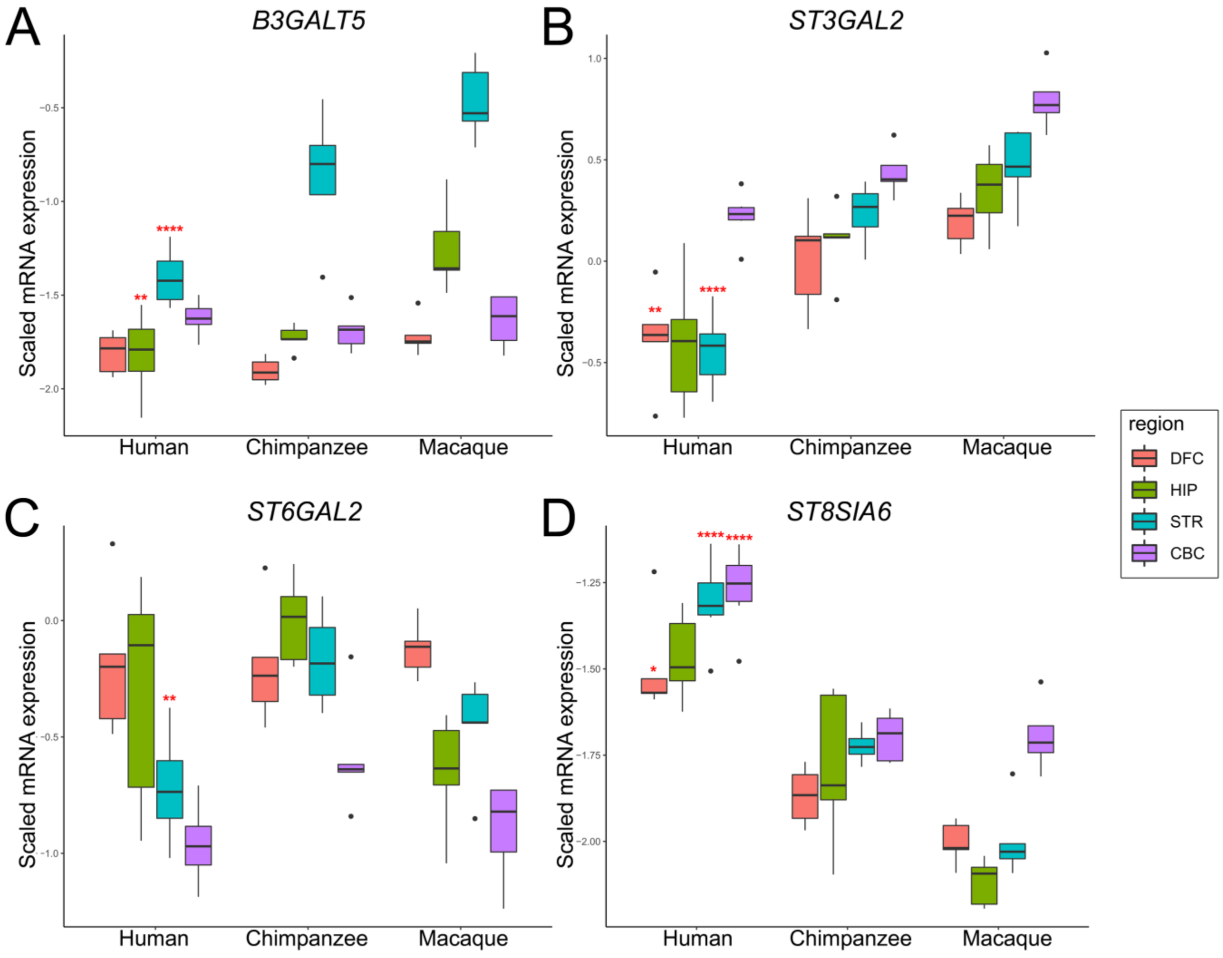


**Fig. S24. Glycogenes with human-specific spatial expression patterns in the brain.** Bulk RNA-Seq expression data for glycogenes that are differentially expressed in humans compared to both non-human primate species in at least one brain region: **(A)** *B3GALT5* (Group IV), **(B)** *ST3GAL2* (Group VI), **(C)** *ST6GAL2* (Group I), and **(D)** *ST8SIA6*. Y-axes show scaled expression of logRPKM values within samples and among the 47 glycogenes. Brain regions in which expression is significantly different in humans compared to both chimpanzees and macaques are denoted by red asterisks: * 0.05 > P > 0.01, ** 0.01 > P 0.001, *** 0.001 P > 0.0001, **** P < 0.0001. P-values as previously determined (*17*).

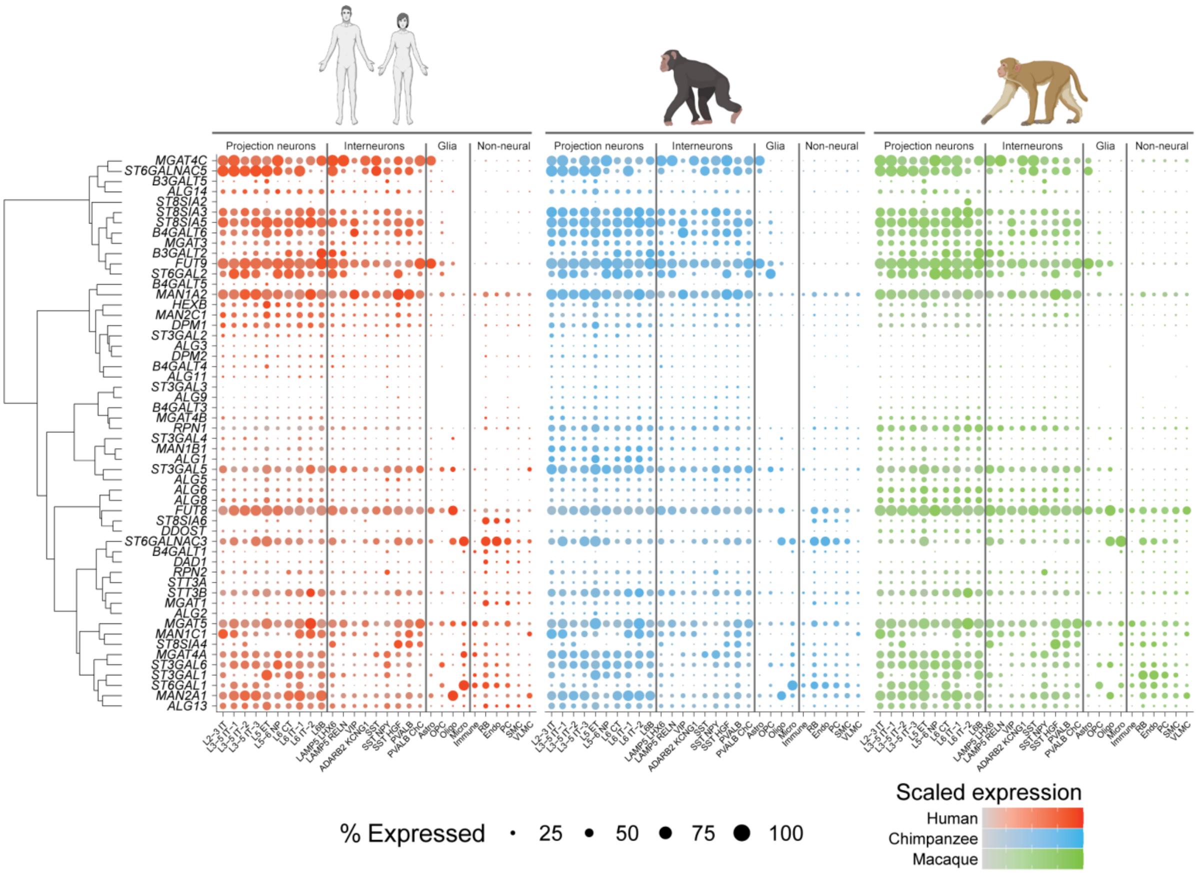


**Fig. S25. Phylogenetic differences in the cell-type distribution of glycogene expression in the primate neocortex.** Expression of glycogenes in the dorsolateral prefrontal cortex of adult humans, chimpanzees and macaques. The sizes and color gradients of the dots represent the expression ratio and the scaled average expression, respectively. Cell type abbreviations: L, layer; IT, intratelencephalic neurons; ET, extratelencephalic neurons; NP, near-projecting; CT, cortical-thalamic projection neurons; ChC, chandelier cells; Astro, astrocytes; OPC, oligodendrocyte precursor cells; Oligo, oligodendrocytes; Micro, microglia; Immune, immune cells; RB, red blood lineage cells; Endo, endothelial cells; SMC, smooth muscle cells; VLMC, vascular leptomeningeal cells.

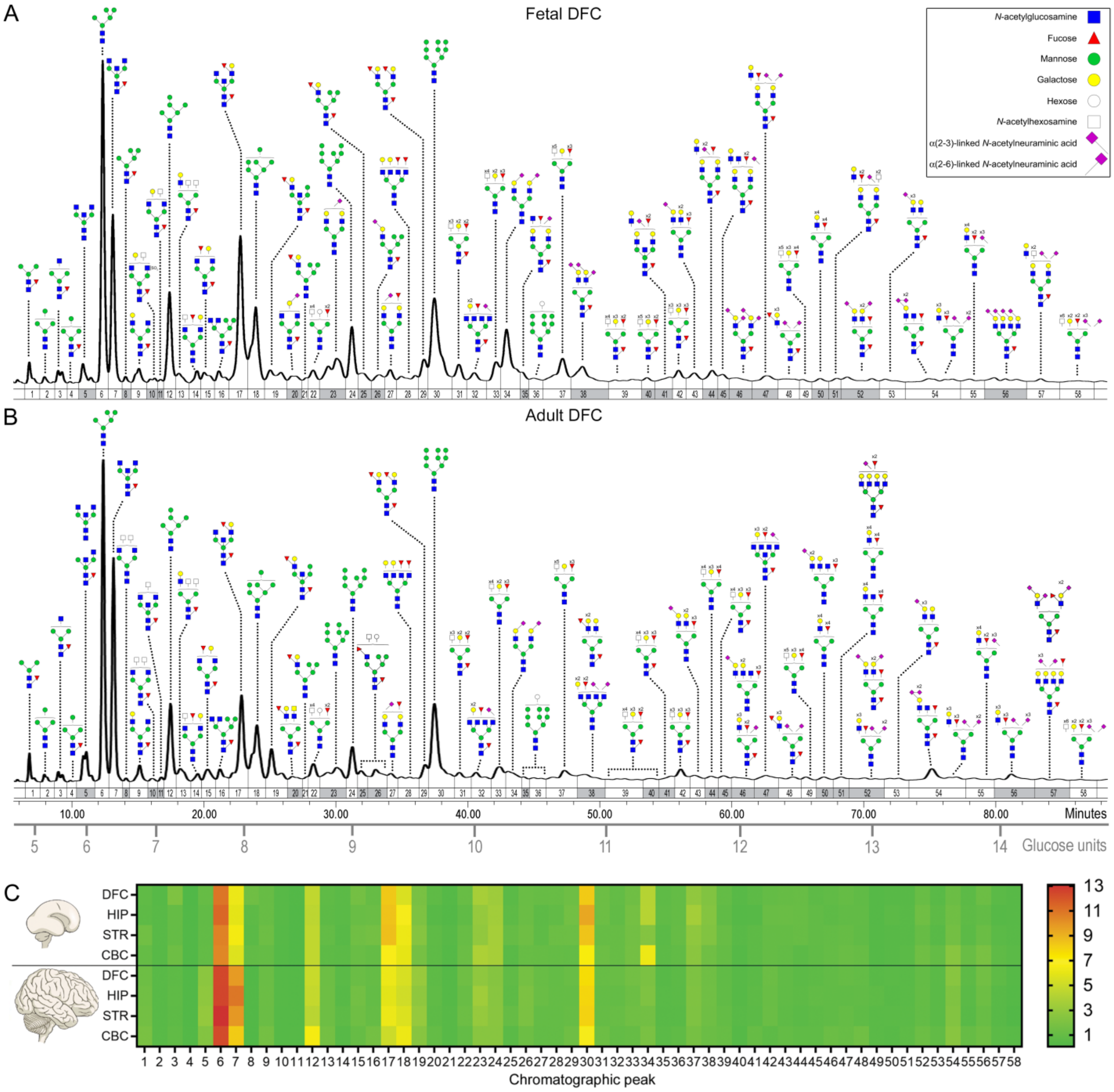


**Fig. S26. The mid-fetal brain has already acquired the archetypal "brain type" N-glycosylation profile by post-conception week 21.** Representative N-glycoprofiles are shown for the **(A)** fetal and **(B)** adult dorsolateral prefrontal cortex. N-glycoprofiles from all brain regions were divided into 58 N-glycan peaks. Integration boundaries are shown as vertical lines. The x-axis displays analyte retention time (minutes), while relative fluorescence at 428 nm is shown in the y-dimension. Glucose units (GUs), as determined from the elution position of 2-AB-labelled glucose oligomers prepared from a standard dextran hydrolysate ladder, are shown below. The chromatogram has been annotated with proposed N-glycan structures. Only the major N-glycan species in each peak are shown (see Table S3 for complete list). Monosaccharide composition was determined by accurate mass MALDI TOF MS, while proposed structural configurations were deduced based on GU value, knowledge of N-glycan biosynthetic pathways, information from the literature, and MALDI TOF/TOF MS/MS fragmentation spectra (where available). The key in the top right corner shows the symbolic representation of monosaccharides used to draw N-glycan structures as proposed by the Consortium for Functional Glycomics (CFG). Analyte peak labels in which the dominant N-glycan structure differs between the fetal and adult DFC are shaded grey to illustrate the qualitative differences between the N-glycomes. **(C)** Mean N-glycan peak abundance in the developing and adult human brain. The heatmap shows the normalized mean relative abundance of each N-glycan chromatographic peak in the brain N-glycome of the fetal (above the horizontal line) and adult brain (below the horizontal line) for each brain region. Abundance is expressed as a percentage of the total integrated chromatographic area. DFC – dorsolateral prefrontal cortex; HIP – hippocampus; STR – striatum; CBC – cerebellar cortex.

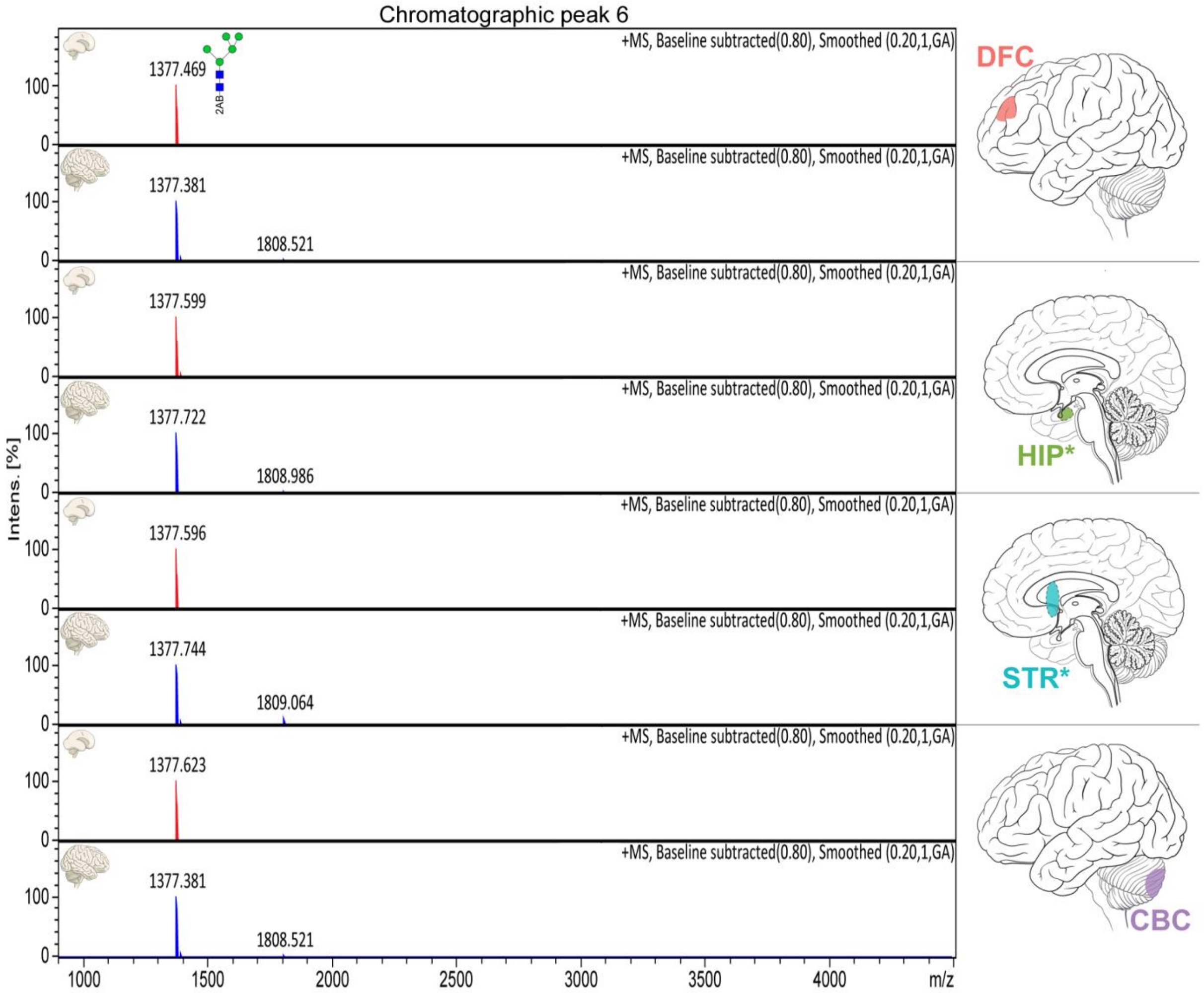


**Fig S27. N-glycan content of chromatographic Peak 6.** MALDI-TOF MS spectra of Peak 6 are displayed for fetal (red) and adult (blue) brain samples. x-axis - mass-to-charge ratio (*m/z*); y-axis - relative ion intensity (% of the base peak). Monoisotopic masses of $[M{:}2\text{-}AB+Na^{+}]^{+}$ ions are displayed above their corresponding ion peaks which are annotated with proposed N-glycan structures. Spectral acquisition and processing parameters are indicated in the top right corner of each spectrum. Brain region is indicated on the right. Icons depict adult brain regions; corresponding nascent structures were sampled in fetal brains. *Denotes that regions are internal and not visible from the view shown. DFC – dorsolateral prefrontal cortex; HIP – hippocampus; STR – striatum; CBC – cerebellar cortex.

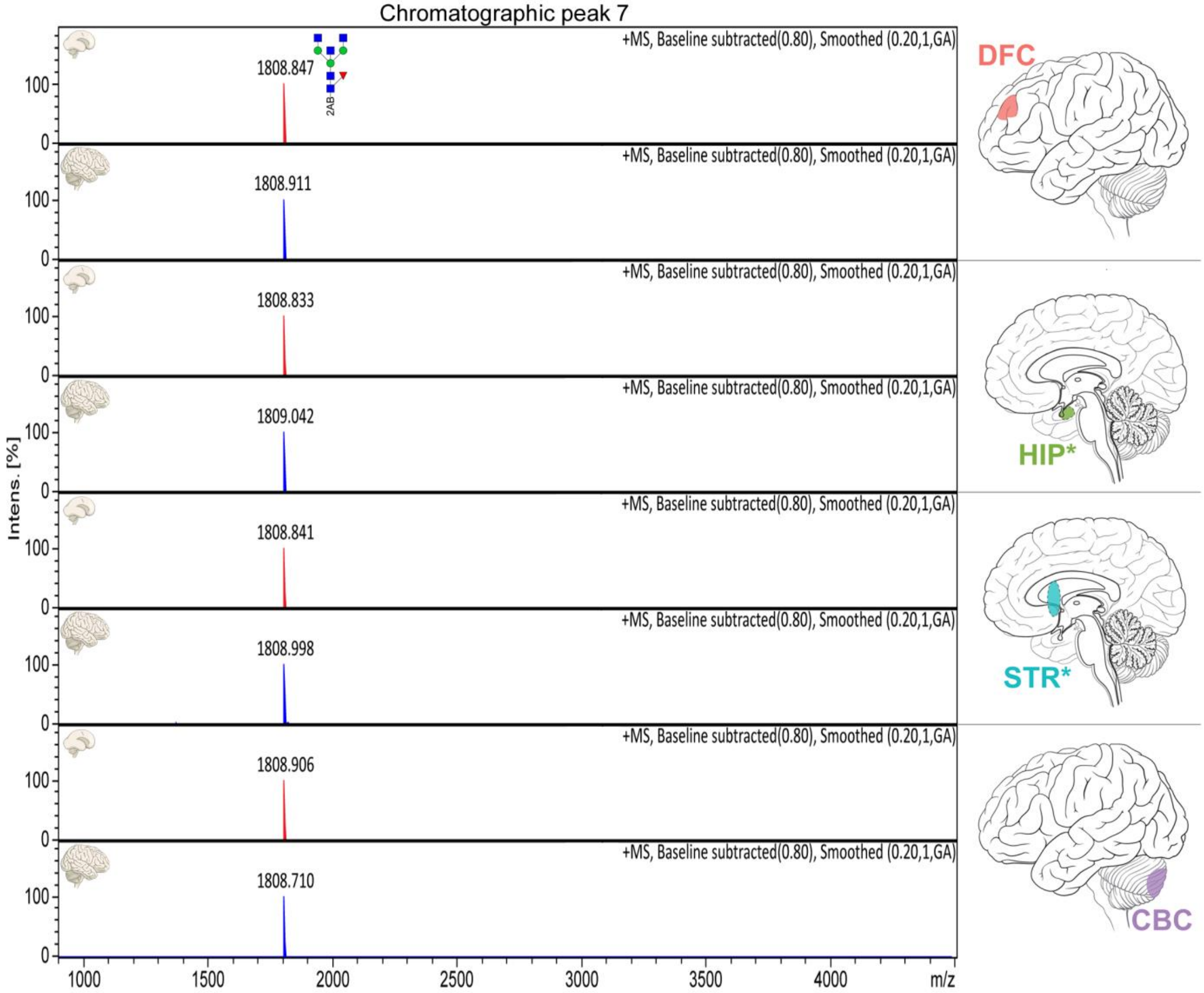


**Fig 28. N-glycan content of chromatographic Peak 7.** MALDI-TOF MS spectra of Peak 7 are displayed for fetal (red) and adult (blue) brain samples. x-axis - mass-to-charge ratio (*m/z*); y-axis - relative ion intensity (% of the base peak). Monoisotopic masses of $[M{:}2\text{-}AB{+}Na^{+}]^{+}$ ions are displayed above their corresponding ion peaks which are annotated with proposed N-glycan structures. Spectral acquisition and processing parameters are indicated in the top right corner of each spectrum. Brain region is indicated on the right. Icons depict adult brain regions; corresponding nascent structures were sampled in fetal brains. *Denotes that regions are internal and not visible from the view shown. DFC – dorsolateral prefrontal cortex; HIP – hippocampus; STR – striatum; CBC – cerebellar cortex.

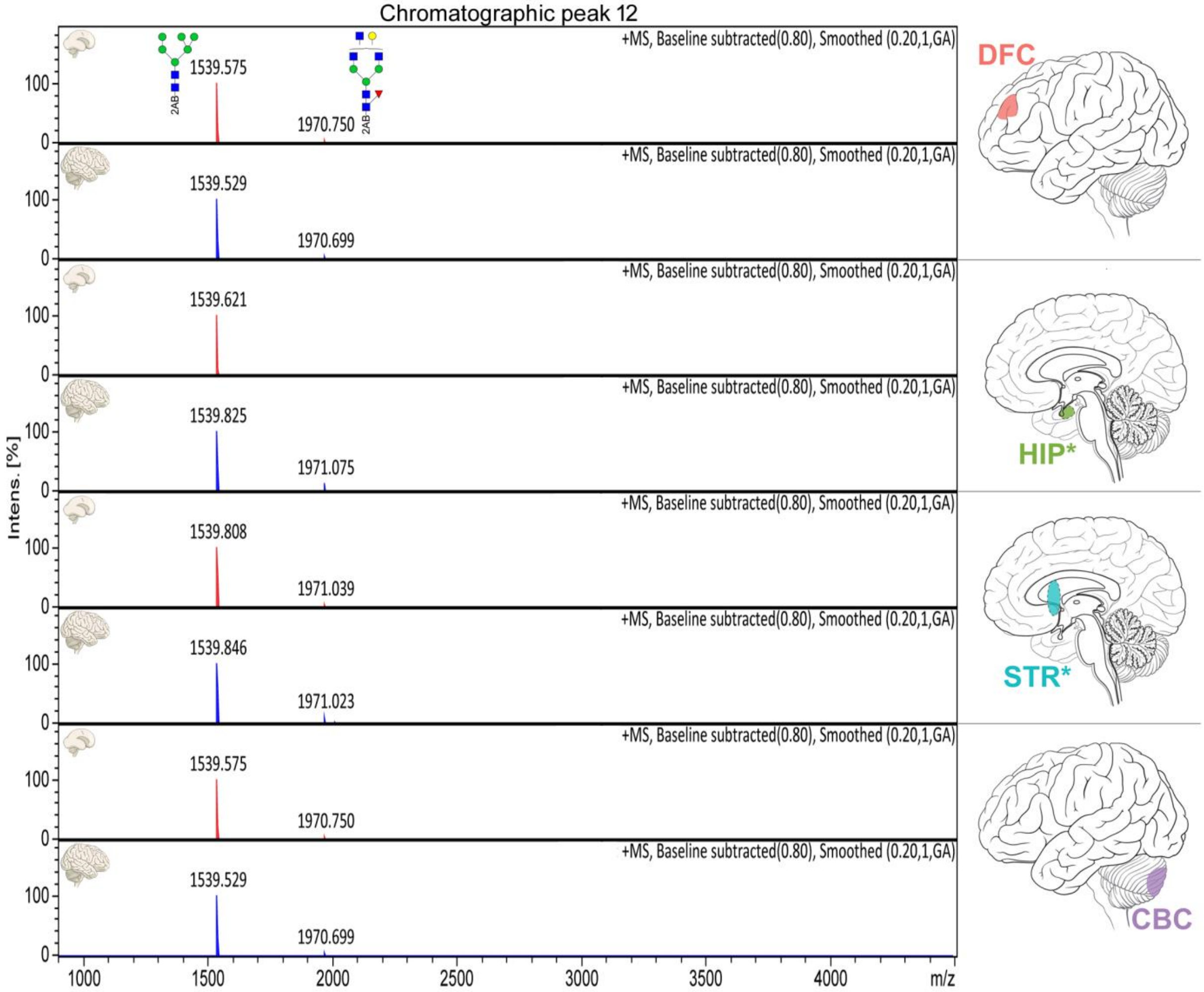


**Fig 29. N-glycan content of chromatographic Peak 12.** MALDI-TOF MS spectra of Peak 12 are displayed for fetal (red) and adult (blue) brain samples. x-axis - mass-to-charge ratio (*m/z*); y-axis - relative ion intensity (% of the base peak). Monoisotopic masses of $[M{:}2\text{-}AB+Na^{+}]^{+}$ ions are displayed above their corresponding ion peaks which are annotated with proposed N-glycan structures. Spectral acquisition and processing parameters are indicated in the top right corner of each spectrum. Brain region is indicated on the right. Icons depict adult brain regions; corresponding nascent structures were sampled in fetal brains. *Denotes that regions are internal and not visible from the view shown. DFC – dorsolateral prefrontal cortex; HIP – hippocampus; STR – striatum; CBC – cerebellar cortex.

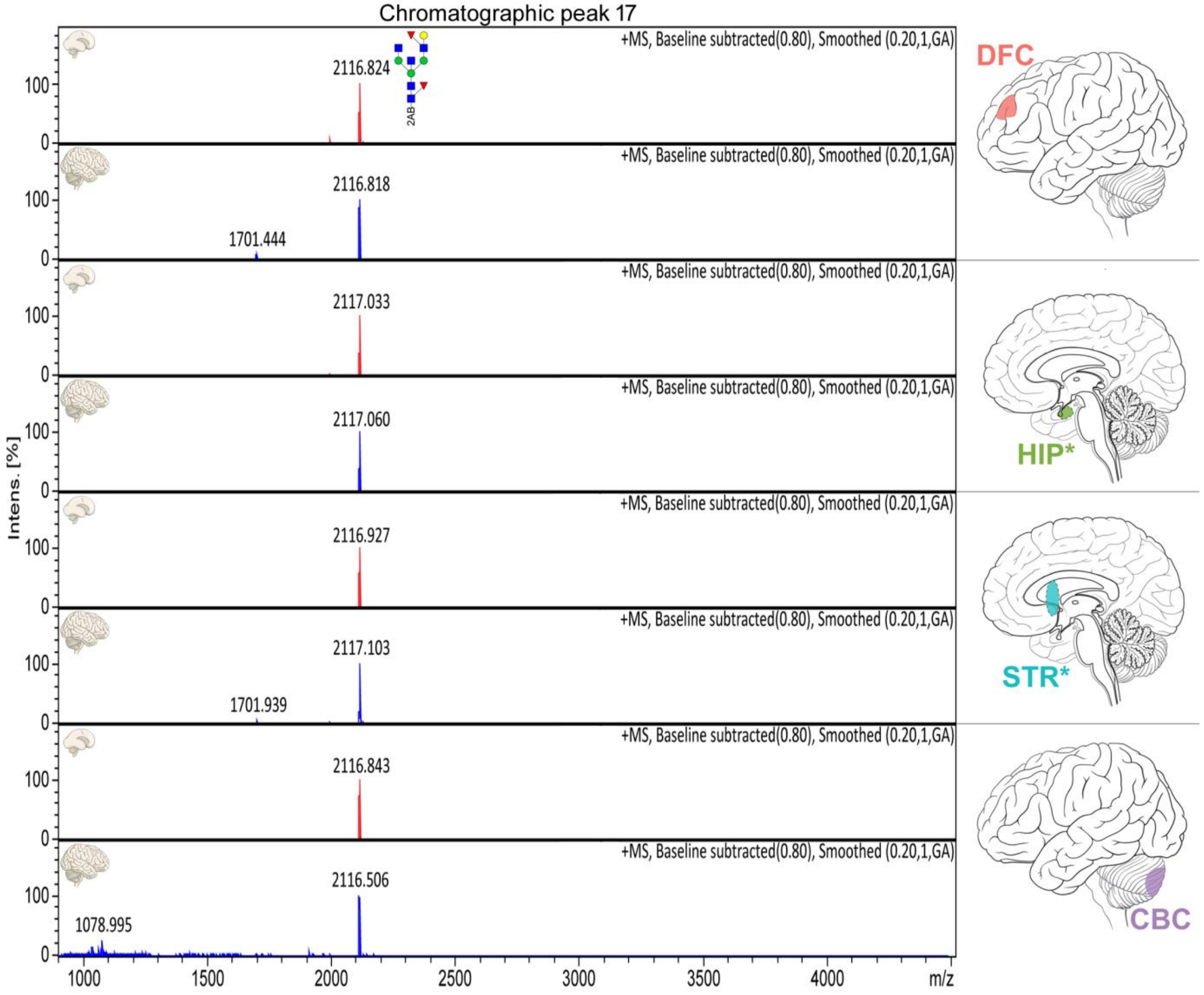


**Fig S30. N-glycan content of chromatographic Peak 17.** MALDI-TOF MS spectra of Peak 17 are displayed for fetal (red) and adult (blue) brain samples. x-axis - mass-to-charge ratio (*m/z*); y-axis - relative ion intensity (% of the base peak). Monoisotopic masses of $[M{:}2\text{-}AB{+}Na^{+}]^{+}$ ions are displayed above their corresponding ion peaks which are annotated with proposed N-glycan structures. Spectral acquisition and processing parameters are indicated in the top right corner of each spectrum. Brain region is indicated on the right. Icons depict adult brain regions; corresponding nascent structures were sampled in fetal brains. *Denotes that regions are internal and not visible from the view shown. DFC – dorsolateral prefrontal cortex; HIP – hippocampus; STR – striatum; CBC – cerebellar cortex.

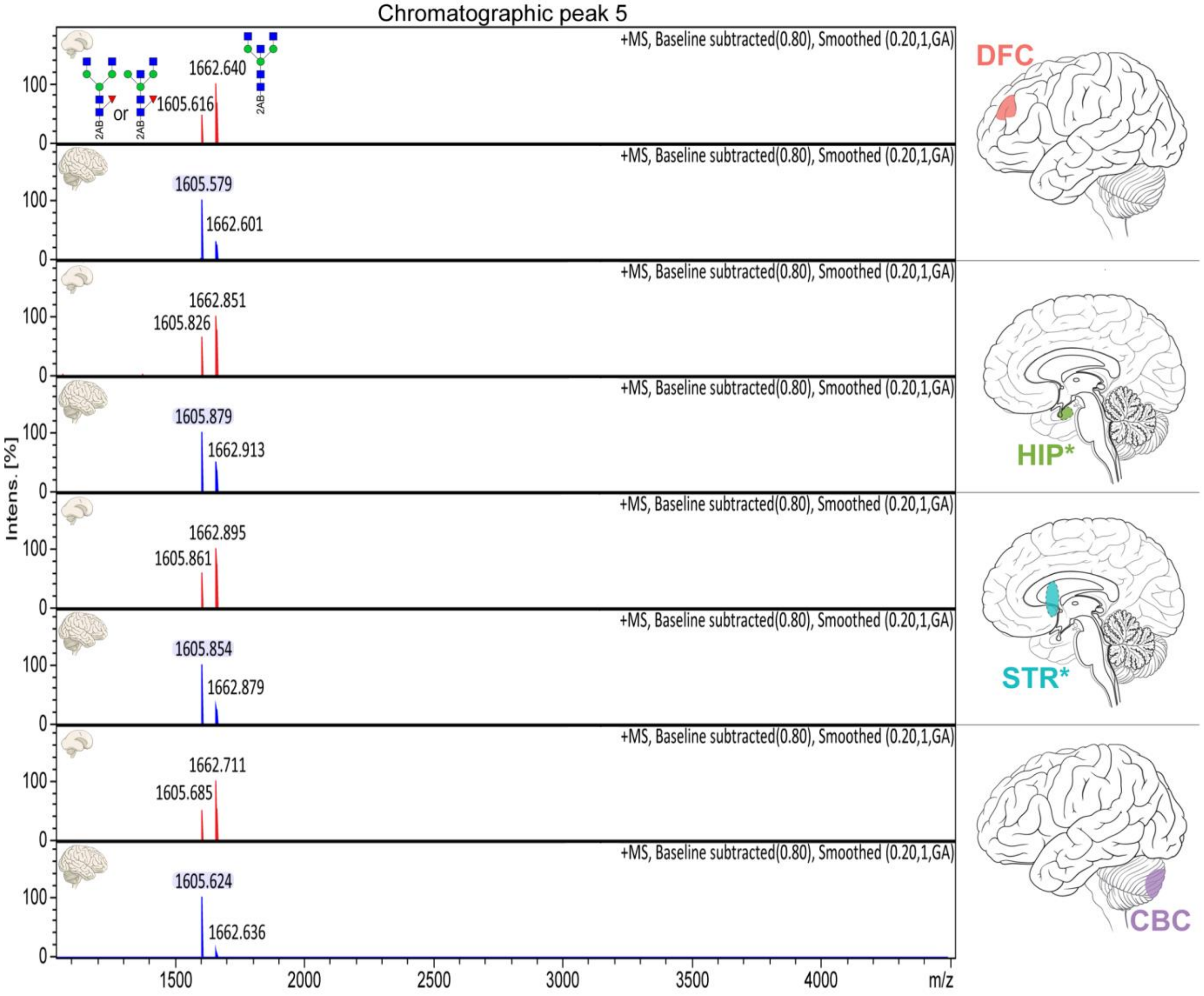


**Fig. S31. Widespread differences in N-glycan peak content between the fetal and adult brain.** MALDI-TOF MS spectra of Peak 5 are displayed for fetal (red) and adult (blue) brain samples. x-axis - mass-to-charge ratio (*m/z*); y-axis - relative ion intensity (% of the base peak). Monoisotopic masses of $[M{:}2\text{-}AB+Na^{+}]^{+}$ ions are displayed above their corresponding ion peaks which are annotated with proposed N-glycan structures. Switching of the major N-glycan species with age was observed in all brain regions; though only a minor signal in the fetal brain, the ion at 1605.60 *m/z* (shaded blue) becomes the dominant ion signal the mature brain. Spectral acquisition and processing parameters are indicated in the top right corner of each spectrum. Brain region is indicated on the right. Icons depict adult brain regions; corresponding nascent structures were sampled in fetal brains. *Denotes that regions are internal and not visible from the view shown. DFC – dorsolateral prefrontal cortex; HIP – hippocampus; STR – striatum; CBC – cerebellar cortex.

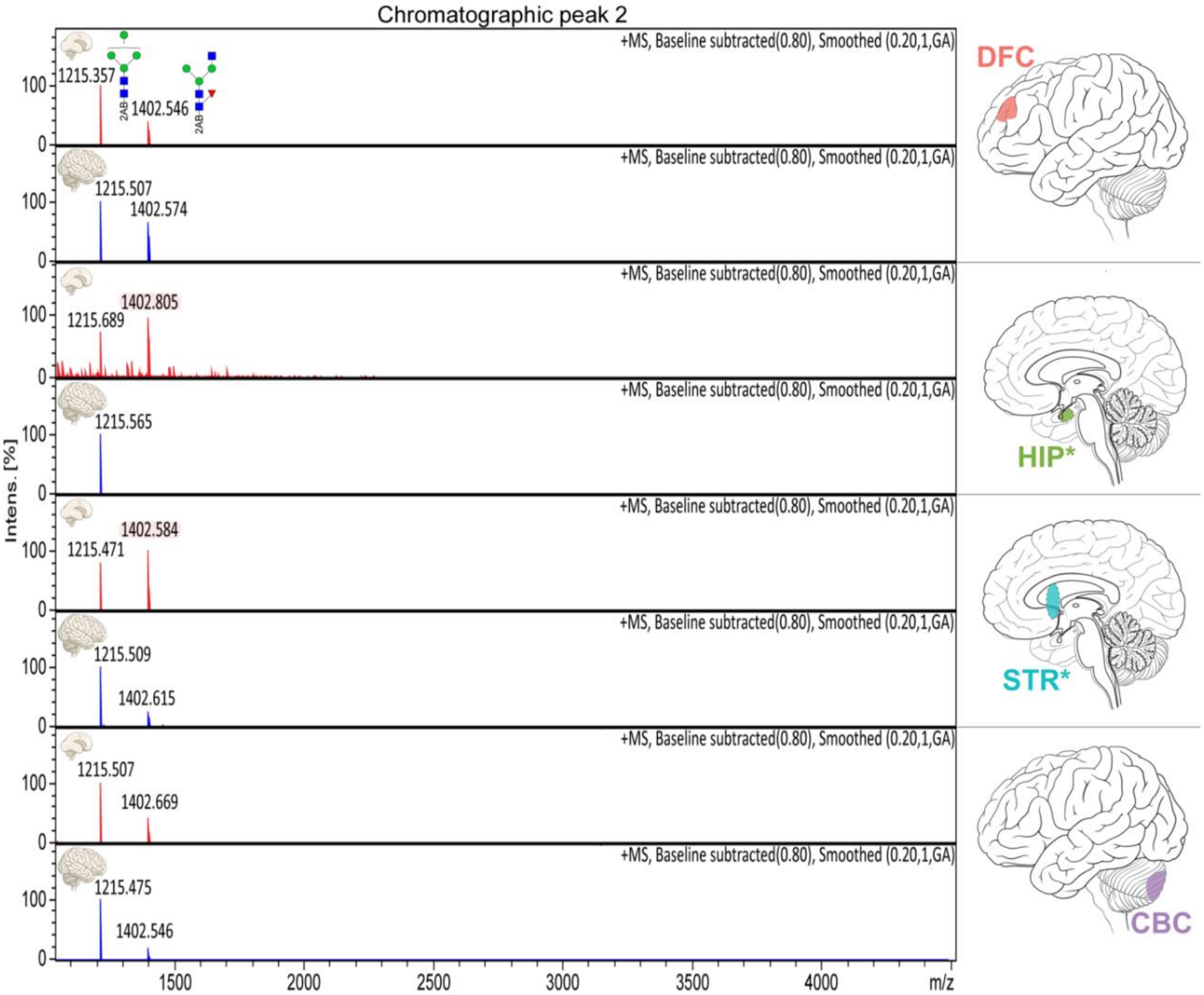


**Fig S32. Region-specific differences in the N-glycan content of Peak 2 between the fetal and adult brain** MALDI-TOF MS spectra of Peak 2 are displayed for fetal (red) and adult (blue) brain samples. x-axis - mass-to-charge ratio (*m/z*); y-axis - relative ion intensity (% of the base peak). Monoisotopic masses of $[M{:}2\text{-}AB+Na^{+}]^{+}$ ions are displayed above their corresponding ion peaks which are annotated with proposed N-glycan structures. Region-specific switching of the major N-glycan species was observed in the HIP and STR where the ion at 1402.52 *m/z* (shaded red) is the dominant ion signal the developing brain. Spectral acquisition and processing parameters are indicated in the top right corner of each spectrum. Brain region is indicated on the right. Icons depict adult brain regions; corresponding nascent structures were sampled in fetal brains. *Denotes that regions are internal and not visible from the view shown.

DFC - dorsolateral prefrontal cortex; HIP – hippocampus; STR – striatum; CBC – cerebellar cortex.

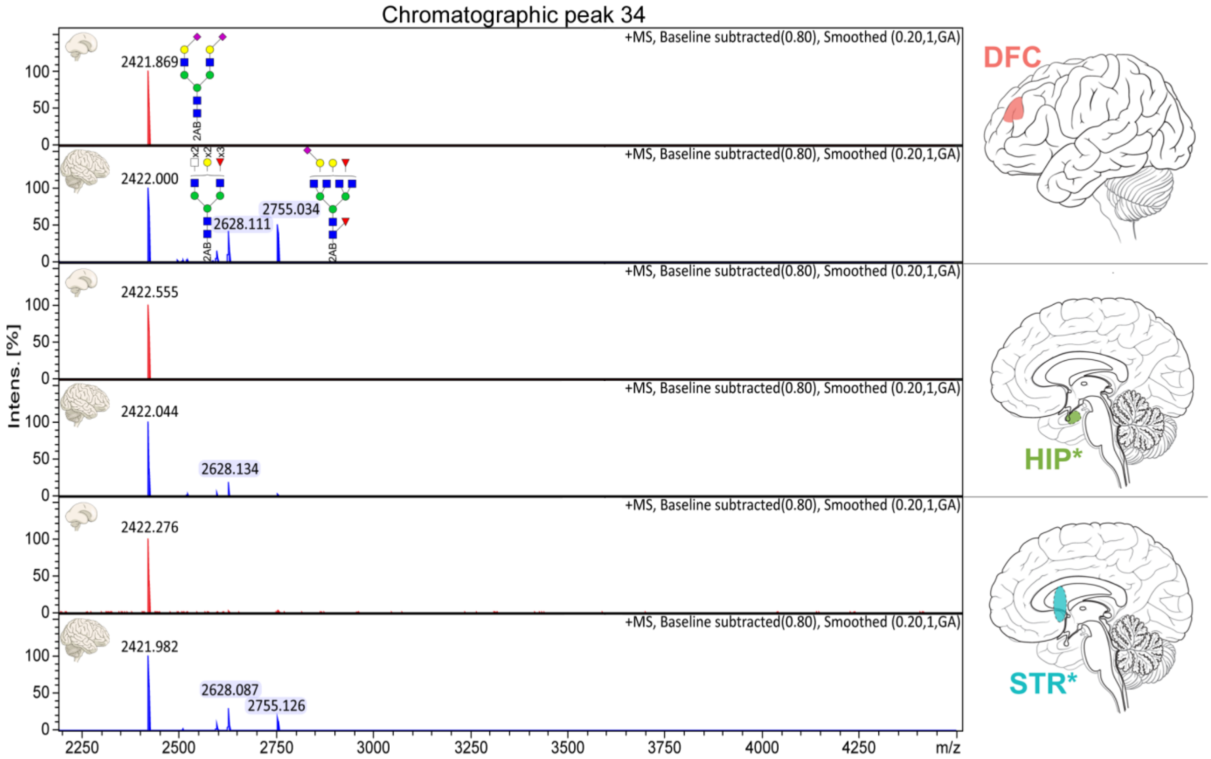


**Fig S33. Region-specific differences in the N-glycan content of Peak 34 between the fetal and adult brain.** Mass spectra recorded upon MALDI-TOF MS analysis of Peak 34 are displayed for fetal (red) and adult (blue) brain samples. Mass-to-charge ratio (*m/z*) is plotted on the x-axis and relative ion intensity, expressed as a percentage of the base peak, is plotted on the y-axis. Monoisotopic masses of singly charged $[M{:}2\text{-}AB{+}Na^{+}]^{+}$ ions are displayed above their corresponding ion peaks which have also been annotated with proposed N-glycan structures. Peak labels of N-glycans specific to the adult brain are shaded in blue. Spectral acquisition and processing parameters are indicated in the top right corner of each spectrum; spectra were acquired in positive ion mode (+MS) and spectra were processed using the default baseline subtraction and smooth functions available in the Data Analysis software. The brain region is indicated by the icon on the right. Icons depict adult brain regions; corresponding nascent structures were sampled in fetal brains. *Denotes that brain regions are internal and not visible from the view shown. DFC – dorsolateral prefrontal cortex; HIP – hippocampus; STR – striatum.

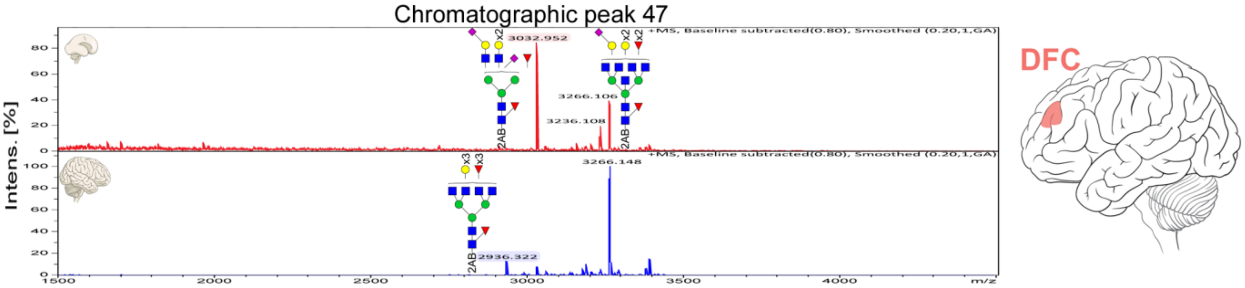


**Fig. S34. Differences in the N-glycan content of Peak 47 between the fetal and adult dorsolateral prefrontal cortex.** Mass spectra recorded upon MALDI-TOF MS analysis of Peak 47 are displayed for fetal (red) and adult (blue) DFC samples. Mass-to-charge ratio (*m/z*) is plotted on the x-axis and relative ion intensity, expressed as a percentage of the base peak, is plotted on the y-axis. In some instances, in-source fragmentation led to large base peaks having low *m/z* values (not visible). Monoisotopic masses of singly charged $[M{:}2AB+Na^{+}]^{+}$ ions are displayed above their corresponding ion peaks which have also been annotated with proposed N-glycan structures. Note the disparity in N-glycan content between age groups. Peak labels of N-glycans specific to the fetal brain are shaded in red, while N-glycans specific to the adult brain are shaded in blue. Spectral acquisition and processing parameters are indicated in the top right corner of each spectrum; spectra were acquired in positive ion mode (+MS) and spectra were processed using the default baseline subtraction and smooth functions available in the Data Analysis software. The icon depicts the adult DFC; the corresponding nascent structure was sampled in fetal brains.

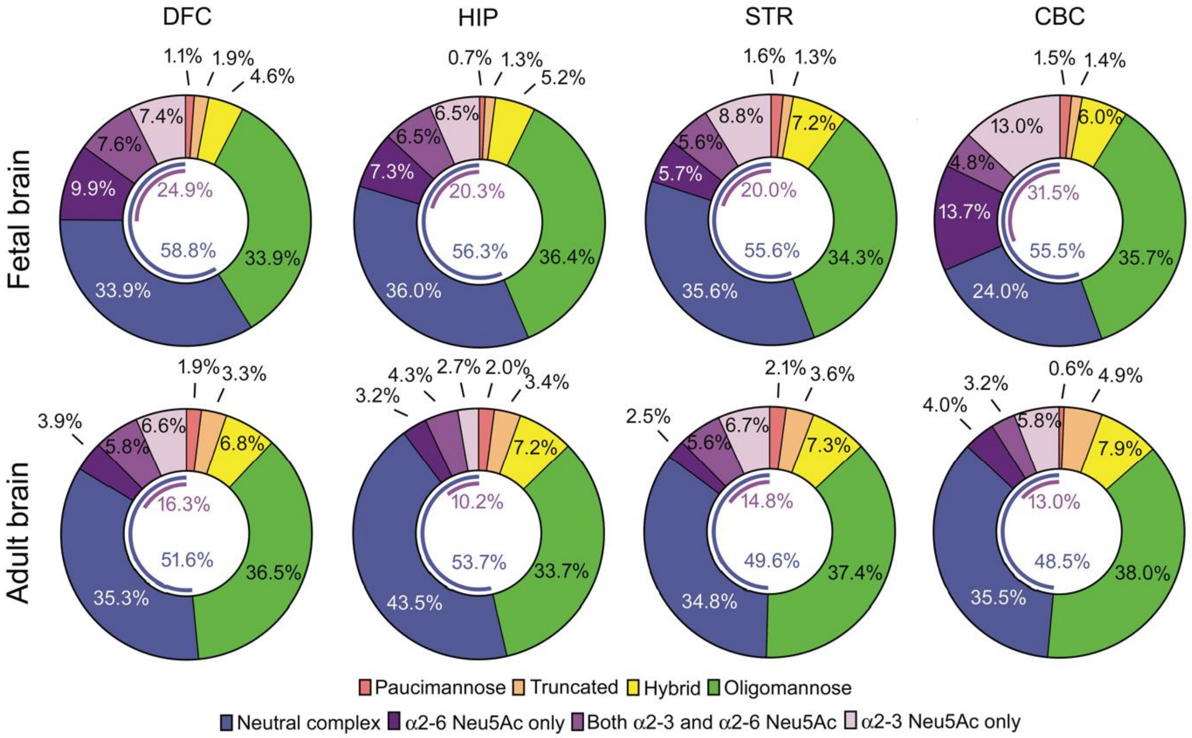


**Fig. S35. Overview of total N-glycome composition by N-glycan type for each group.** Donut charts are aligned as a matrix with rows defining the developmental stage and columns defining the brain region. Derived traits were calculated from UPLC N-glycan peak abundance data and were used to determine the abundance of each category of N-glycans (expressed as a percentage of the total N-glycome by chromatographic area). Mean values for each group are displayed. The blue arc in the donut hole indicates the percentage of total complex N-glycans (i.e. the sum of neutral and sialylated N-glycans). The purple arc in the donut hole indicates the percentage of sialylated complex N-glycans (i.e. the sum of all complex N-glycans containing Neu5Ac). DFC – dorsolateral prefrontal cortex; HIP – hippocampus; STR – striatum; CBC – cerebellar cortex.

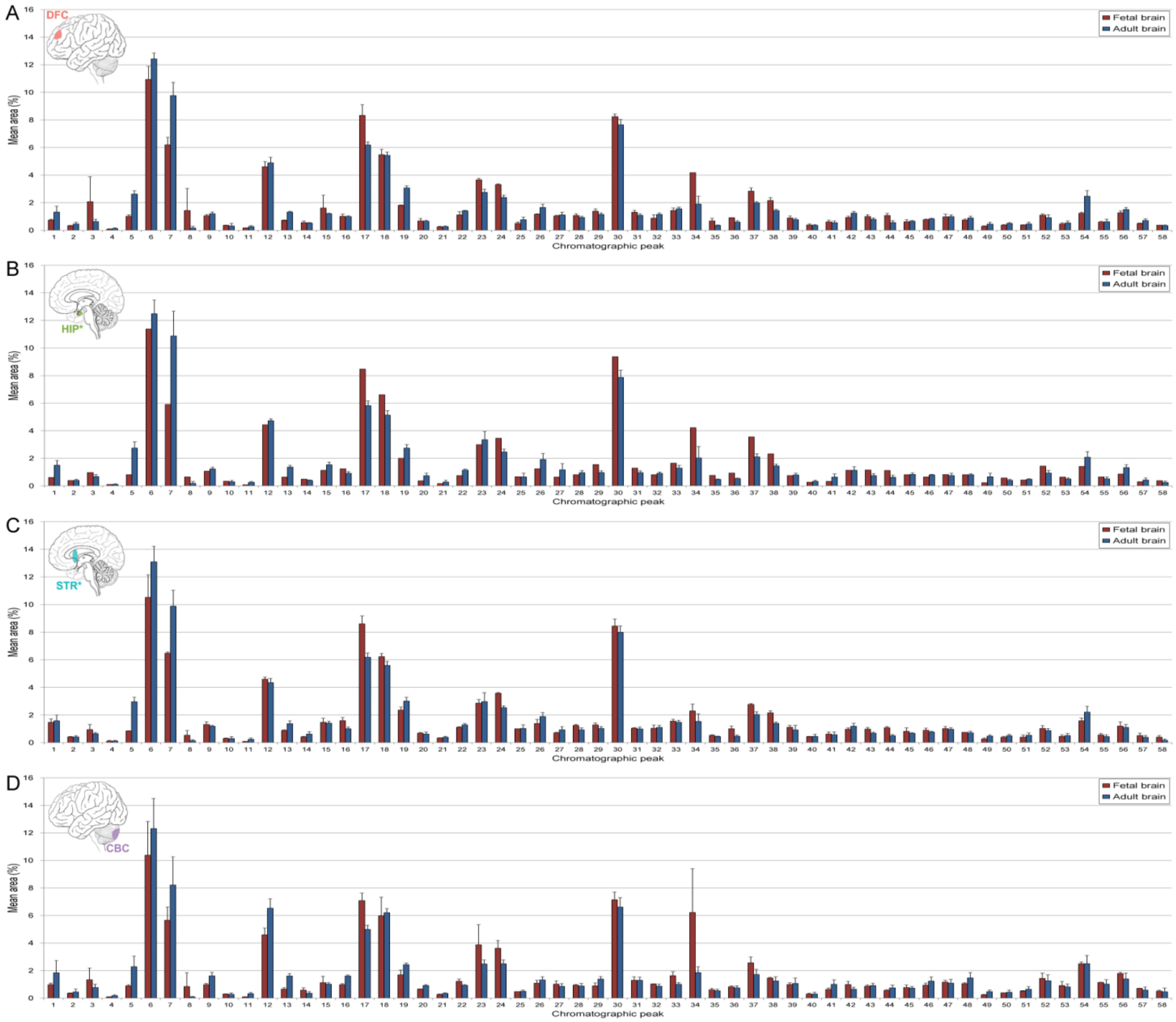


**Fig. S36. N-glycan peak abundance in the developing and adult human brain.** Normalized mean relative abundance of each N-glycan chromatographic peak in the brain N-glycome of the **(A)** dorsolateral prefrontal cortex, **(B)** hippocampus, **(C)** striatum, and **(D)** cerebellar cortex is shown for the fetal brain (red bars) and adult brain (blue bars). Abundance is expressed as a percentage of the total integrated chromatographic area. Error bars show the standard deviation. Statistical significance between age groups was assessed using the Wilcoxon-Mann-Whitney test but no statistically significant differences were found after P-values were adjusted for multiple testing (likely due to small sample size). Icons depict adult brain regions; corresponding nascent structures were sampled in fetal brains. *Denotes that brain regions are internal and not visible from the view shown.

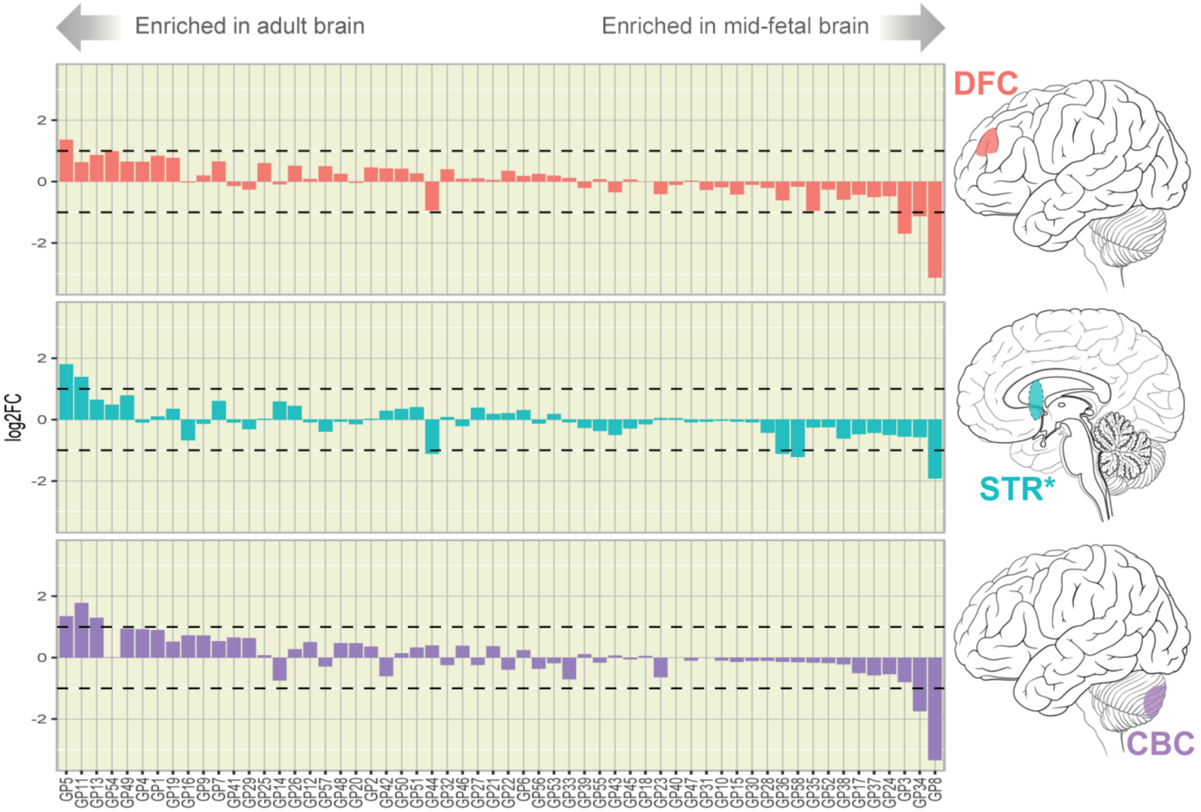


**Fig. S37. Temporal dynamics of N-glycan peak abundance in the human brain during neurodevelopment.** Each column corresponds to one chromatographic peak with peak labels displayed on the bottom. The y-axis shows the binary logarithm of the fold change in peak abundance between the adult and fetal brain. Positive fold change indicates peak abundance is greater in the adult brain, while negative fold change indicates that it is greater in the fetal brain. Statistical significance was assessed using the Wilcoxon-Mann-Whitney test. No statistically significant differences were found after P-values were adjusted for multiple testing (likely due to small sample size). The brain region is indicated by the icon on the right. Hippocampal samples were not analyzed due to an insufficient number of fetal samples. Icons depict adult brain regions; corresponding nascent structures were sampled in fetal brains. *Denotes that brain regions are internal and not visible from the view shown. DFC – dorsolateral prefrontal cortex; STR – striatum; CBC – cerebellar cortex.

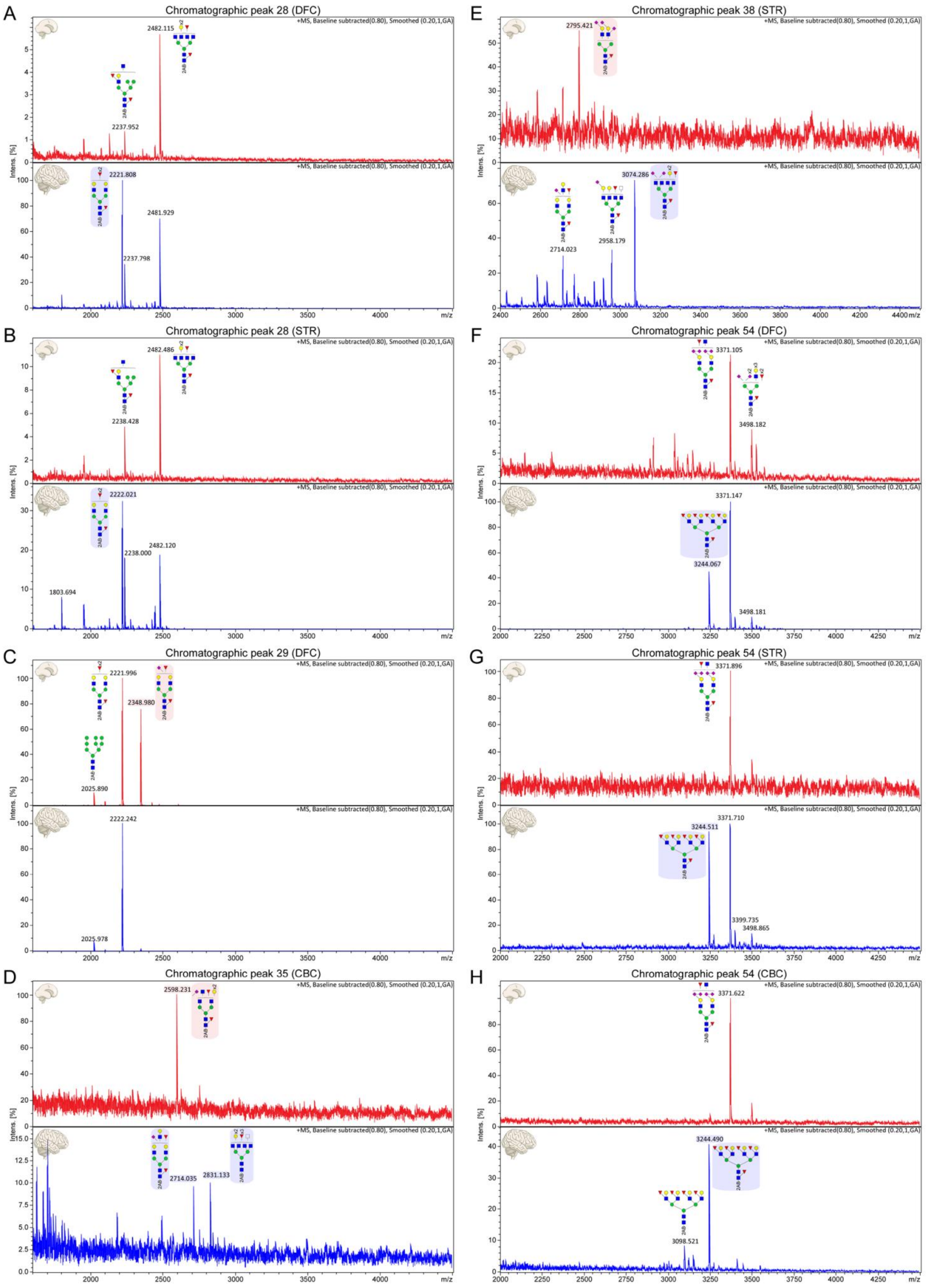
A
Chromatographic peak 28 (DFC)
2482.115
2237.952
2221.808
2481.929
2237.798
B
Chromatographic peak 28 (STR)
2482.486
2238.428
2222.021
2482.120
2238.000
1803.694
C
Chromatographic peak 29 (DFC)
2221.996
2348.980
2025.890
2222.242
2025.978
D
Chromatographic peak 35 (CBC)
2598.231
2714.035
2831.133
E
Chromatographic peak 38 (STR)
2795.421
3074.286
2714.023
2958.179
F
Chromatographic peak 54 (DFC)
3371.105
3498.182
3371.147
3244.067
3498.181
G
Chromatographic peak 54 (STR)
3371.896
3244.511
3371.710
3399.735
3498.865
H
Chromatographic peak 54 (CBC)
3371.622
3244.490
3098.521
Intens. [%]
m/z

**Fig. S38. Identification of N-glycans specific to either the developing or adult human brain.** MALDI-TOF MS was used to investigate qualitative differences in N-glycan peak content between the fetal brain (red spectra, top) and the adult brain (blue spectra, bottom): **(A)** Peak 28, DFC; **(B)** Peak 28, STR; **(C)** Peak 29, DFC; **(D)** Peak 35, CBC; **(E)** Peak 38, STR; **(F)** Peak 54a, DFC; **(G)** Peak 54a, STR; **(H)** Peak 54a, CBC. N-glycans specific to the fetal brain are shaded in red, while N-glycans specific to the adult brain are shaded in blue. Mass-to-charge ratio (*m/z*) is plotted on the x-axis and relative ion intensity, expressed as a percentage of the base peak, is plotted on the y-axis. In some instances, in-source fragmentation led to large base peaks having low *m/z* values (not visible). Monoisotopic masses of singly charged $[M{:}2AB+Na^+]^+$ ions are displayed above their corresponding ion peaks which have also been annotated with proposed N-glycan structures. Spectral acquisition and processing parameters are indicated in the top right corner of each spectrum; spectra were acquired in positive ion mode (+MS) and spectra were processed using the default baseline subtraction and smooth functions available in the Data Analysis software. DFC – dorsolateral prefrontal cortex; STR – striatum; CBC – cerebellar cortex.

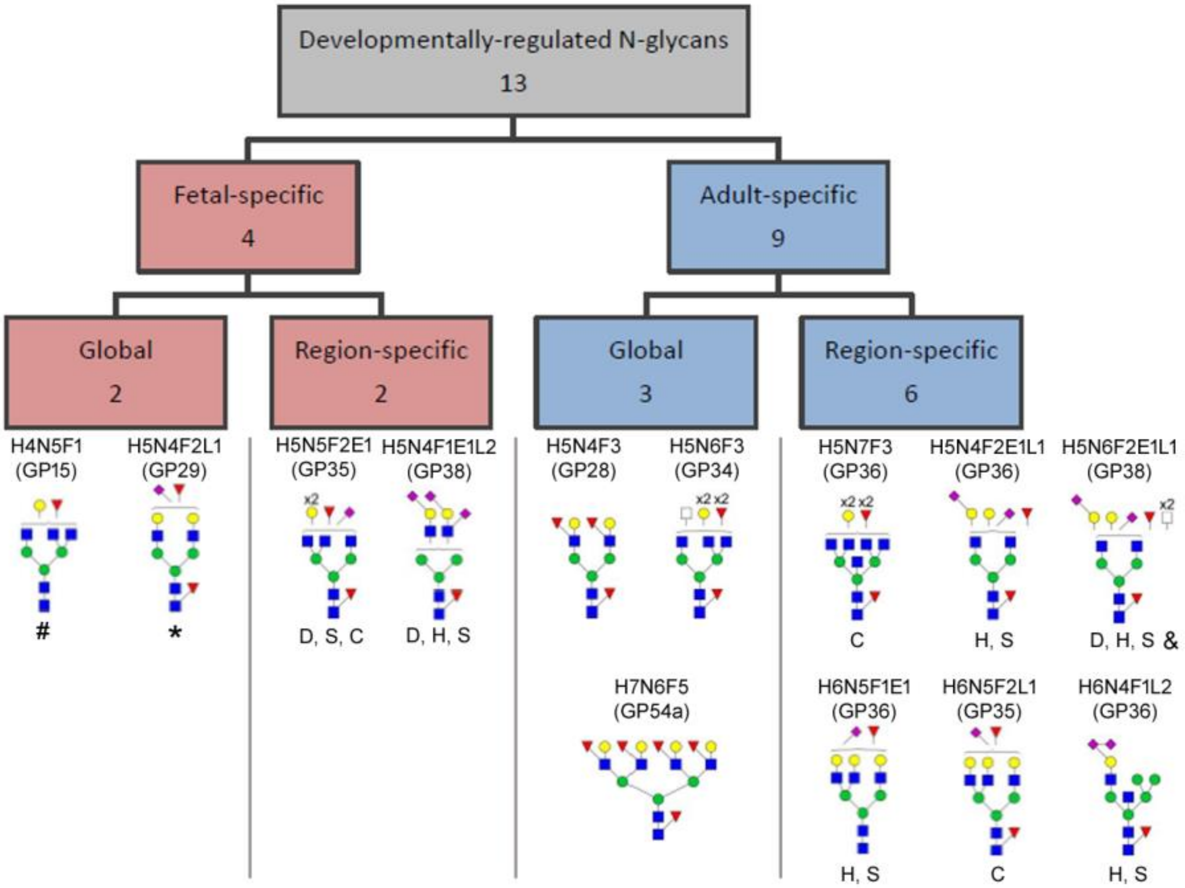


**Fig. S39. Overview of developmentally-regulated N-glycans.** In total, 13 developmentally-regulated N-glycans were identified in this study. The majority (9) were enriched in the adult brain while only four structures were enriched in the fetal brain (Table S3). These N-glycans were further subcategorized into those that were enriched globally (i.e. in all brain regions analyzed) or those that were developmentally-enriched only in certain brain regions. Proposed structures for each of the 13 developmentally-regulated N-glycans are displayed. The monosaccharide composition of each N-glycan and the chromatographic peak in which it was identified are indicated above the corresponding structure. For region-specific developmentally-regulated N-glycans, the brain region(s) in which they were enriched are indicated below the structures: D – dorsolateral prefrontal cortex; H – hippocampus; S - striatum; C – cerebellar cortex. The composition of the structure indicated by an asterisk (*) matches the composition of an N-glycan also found to be enriched in the human fetal prefrontal cortex (*13*). The composition of the structure (GU value 7.04) indicated by a hash (#) matches the compositions of two isomers (GU values 6.70 and 7.04) that were also enriched in the neonatal rat neocortex (*23*) as well as two isomers, termed Ga-BA-2 and Gb-BA-2 (GU values not reported), that were highly enriched in the embryonic mouse cerebral cortex (*26*). The composition of the structure (GU 10.84) indicated by an ampersand (&) matches the composition of a pair of N-glycan isomers (GU 10.71 and 11.07) that were also found to be enriched in the adult rat neocortex (*23*). Monosaccharide code: H – hexose; N – *N*-Acetylhexosamine; F - deoxyhexose (i.e. fucose); E – esterified Neu5Ac [i.e. α(2-6)-linked]; L – lactonized Neu5Ac [i.e. α(2-3)- or α(2-8)-linked].

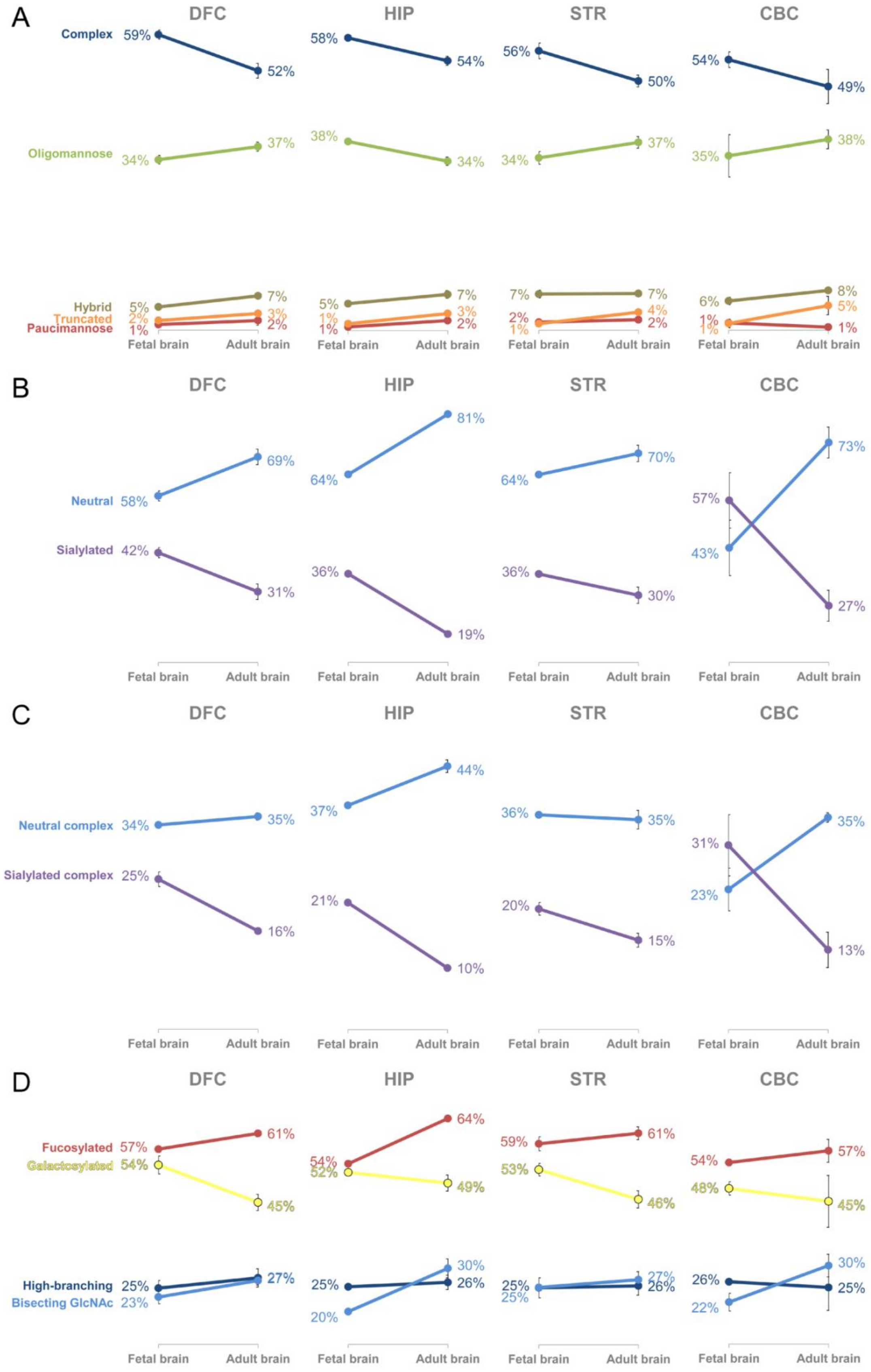
A
DFC
HIP
STR
CBC
Complex
59%
52%
58%
54%
56%
50%
54%
49%
Oligomannose
34%
37%
38%
34%
34%
37%
35%
38%
Hybrid
Truncated
Paucimannose
5%
2%
1%
7%
3%
2%
5%
1%
1%
7%
3%
2%
7%
2%
1%
7%
4%
2%
6%
1%
1%
8%
5%
1%
Fetal brain
Adult brain
B
DFC
HIP
STR
CBC
Neutral
58%
69%
64%
81%
64%
70%
43%
73%
Sialylated
42%
31%
36%
19%
36%
30%
57%
27%
Fetal brain
Adult brain
C
DFC
HIP
STR
CBC
Neutral complex
34%
35%
37%
44%
36%
35%
23%
35%
Sialylated complex
25%
16%
21%
10%
20%
15%
31%
13%
Fetal brain
Adult brain
D
DFC
HIP
STR
CBC
Fucosylated
57%
61%
54%
64%
59%
61%
54%
57%
Galactosylated
54%
45%
52%
49%
53%
46%
48%
45%
High-branching
Bisecting GlcNAc
25%
23%
27%
25%
20%
30%
26%
25%
25%
27%
26%
26%
22%
30%
25%
Fetal brain
Adult brain

**Fig. S40. Developmental trends in N-glycan derived traits by brain region.** Slope graphs show the change in relative abundance of each N-glycan derived trait from the fetal stage to adulthood by brain region. **(A)** Relative abundance of each N-glycan class expressed as a percentage of the total N-glycome. **(B)** Relative abundance of neutral and sialylated complex N-glycans expressed as a percentage of total complex N-glycans. **(C)** Relative abundance of neutral and sialylated complex N-glycans expressed as a percentage of the total N-glycome. **(D)** Relative abundance of N-glycans carrying particular modifications expressed as a percentage of the total N-glycome. DFC – dorsolateral prefrontal cortex; HIP – hippocampus; STR – striatum; CBC – cerebellar cortex. Error bars show the standard deviation.

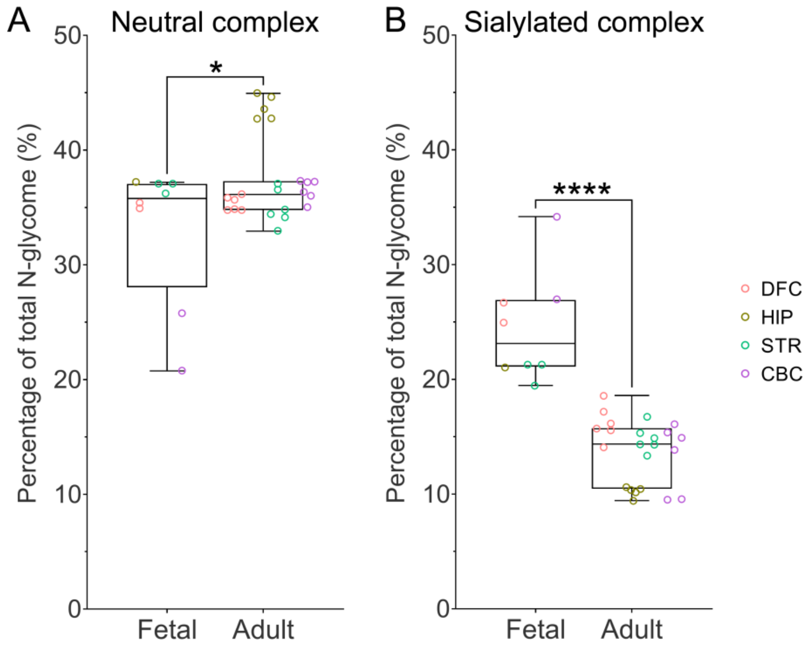


**Fig. S41. Quantification of neutral and sialylated complex N-glycans in the developing and adult human brain.** Box and whisker plots show the data spread including the median value, interquartile range, maximum, and minimum. Data points are colored by brain region. Aggregate data for **(A)** Neutral complex and **(B)** Sialylated complex N-glycans are presented as the proportion of the total N-glycome and were analyzed using unpaired, two-tailed t-tests with correction for multiple testing using the Bonferroni procedure. Adjusted P-values: * $0.01 < P < 0.05$, **** $P < 0.0001$. DFC – dorsolateral prefrontal cortex; HIP – hippocampus; STR – striatum; CBC – cerebellar cortex. Error bars show the standard deviation.

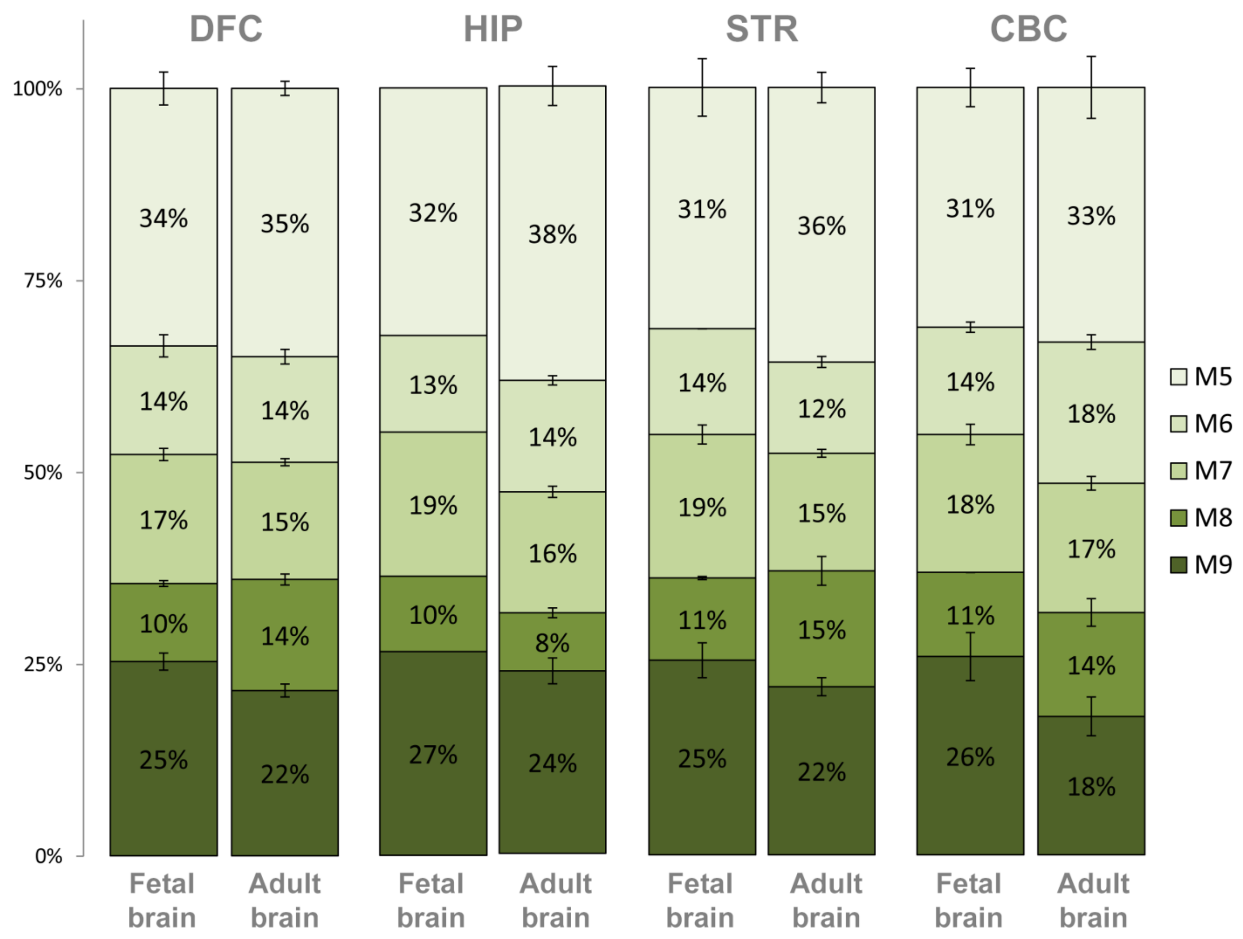


**Fig. S42 Distribution of oligomannose N-glycan subtypes in the developing and adult human brain.** Distribution of oligomannose subtypes in the developing and adult human brain by region. Data are expressed as a percentage of total standard oligomannose N-glycans (unmodified M5-M9 series only, i.e. excluding M9+Glc and the core fucosylated M5 form).

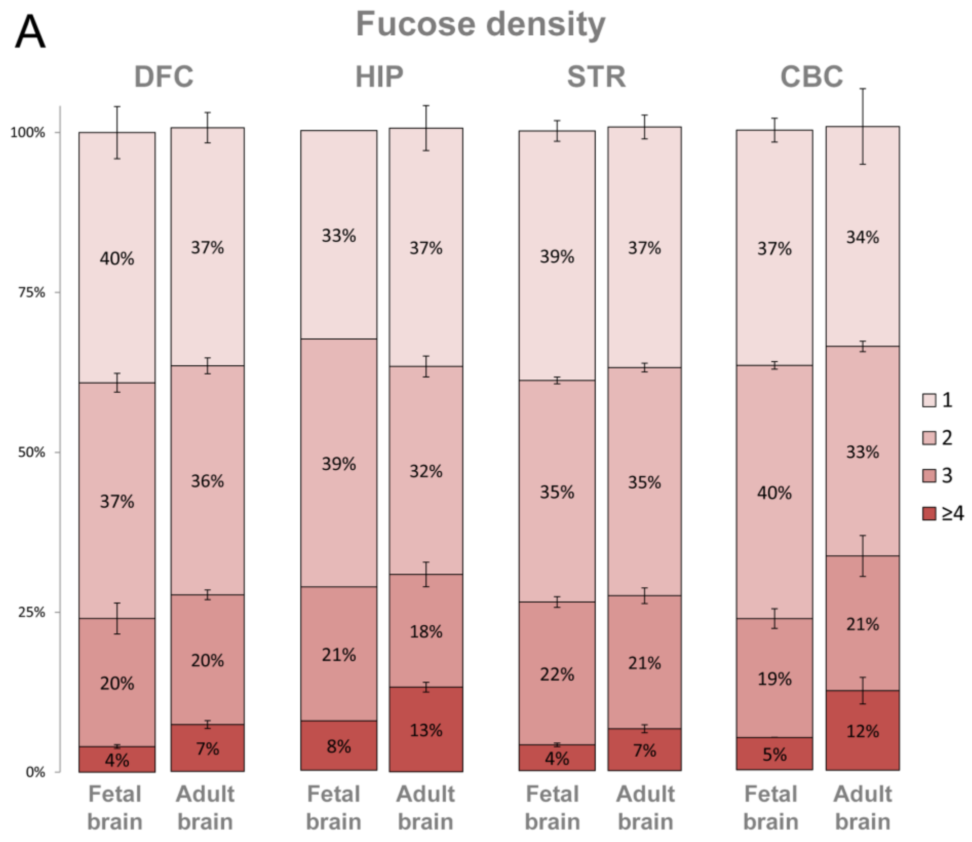
A
Fucose density
DFC
HIP
STR
CBC
100%
75%
50%
25%
0%
40%
37%
20%
4%
37%
36%
20%
7%
33%
39%
21%
8%
37%
32%
18%
13%
39%
35%
22%
4%
37%
35%
21%
7%
37%
40%
19%
5%
34%
33%
21%
12%
1
2
3
≥4
Fetal brain
Adult brain
Fetal brain
Adult brain
Fetal brain
Adult brain
Fetal brain
Adult brain

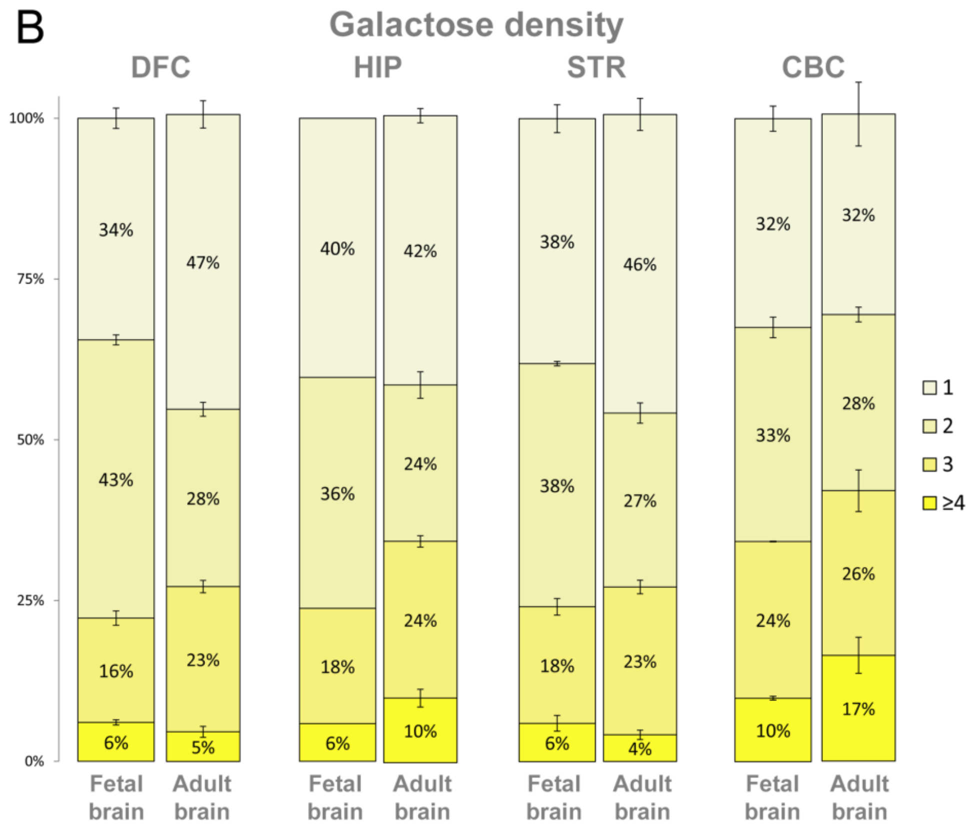
B
Galactose density
DFC
HIP
STR
CBC
100%
75%
50%
25%
0%
34%
43%
16%
6%
47%
28%
23%
5%
40%
36%
18%
6%
42%
24%
24%
10%
38%
38%
18%
6%
46%
27%
23%
4%
32%
33%
24%
10%
32%
28%
26%
17%
1
2
3
≥4
Fetal brain
Adult brain
Fetal brain
Adult brain
Fetal brain
Adult brain
Fetal brain
Adult brain

**Fig. S43. Fucosylation and galactosylation density of N-glycans in the developing and adult human brain. (A)** Fucosylation density, defined as the number of fucose residues per N-glycan molecule, in the fetal and adult human brain by region. Data are expressed as a percentage of total fucosylated N-glycans. **(B)** Galactosylation density, defined as the number of galactose residues per N-glycan molecule, in the fetal and adult human brain by region. Data are expressed as a percentage of total galactosylated N-glycans. Error bars show the standard deviation. DFC – dorsolateral prefrontal cortex; HIP – hippocampus; STR – striatum; CBC – cerebellar cortex.

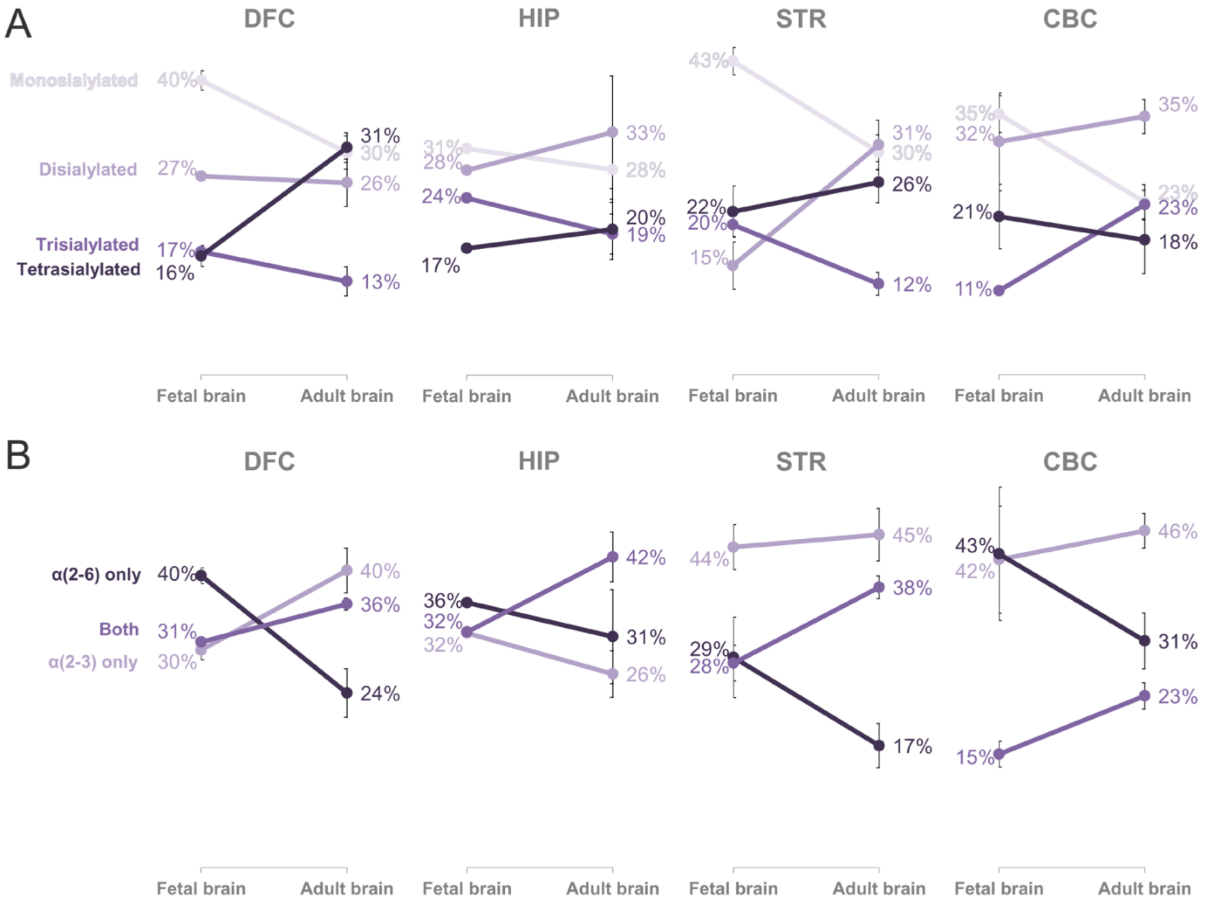


**Fig. S44. Developmental trends in sialylation by brain region.** Slope graphs show the developmental trend of each sialylation trait from the fetal stage to adulthood by brain region. **(A)** Sialylation density (defined as the number of Neu5Ac residues per N-glycan molecule) expressed as a percentage of sialylated complex N-glycans. **(B)** Distribution of Neu5Ac linkage on N-glycans of the developing and adult human brain. Sialylated complex N-glycans were grouped into those that had only α(2-3)-linked Neu5Ac, only α(2-6)-linked Neu5Ac, or both α(2-3)- and α(2-6)-linked Neu5Ac. DFC – dorsolateral prefrontal cortex; HIP – hippocampus; STR – striatum; CBC – cerebellar cortex. Error bars show the standard deviation.

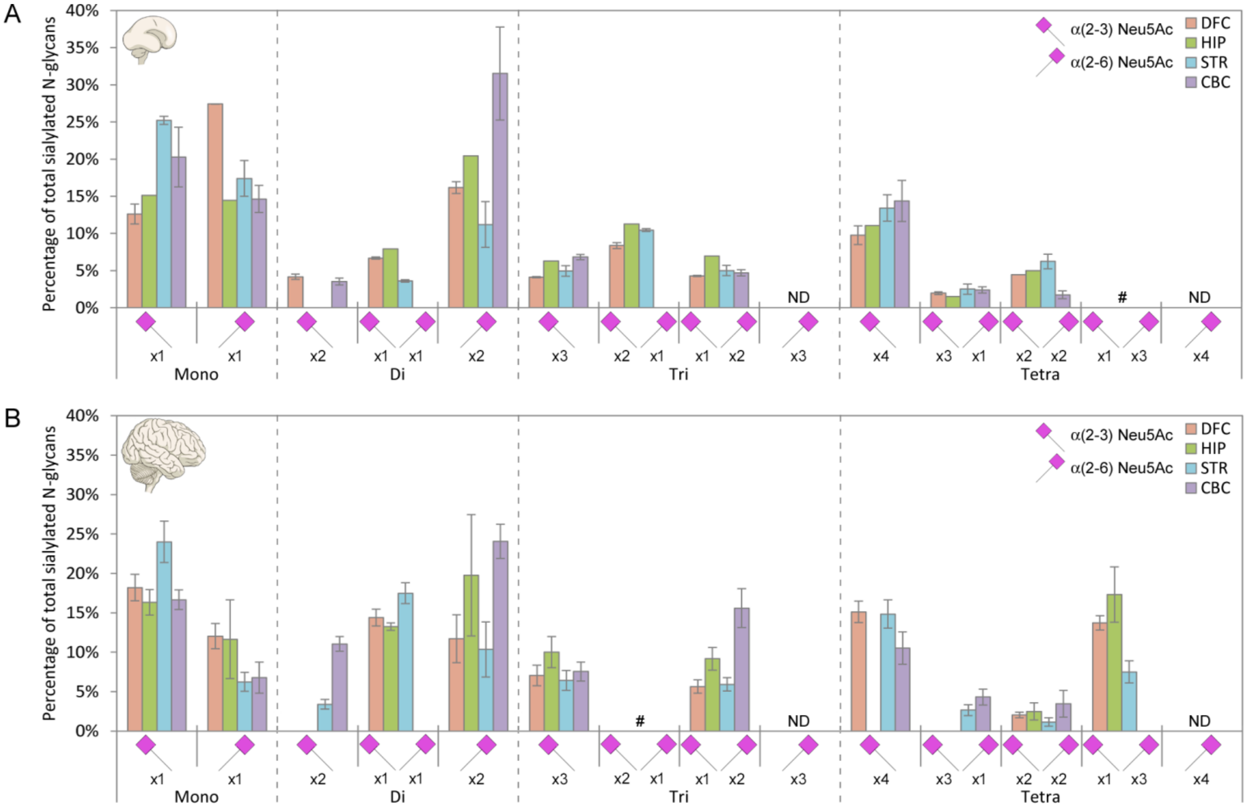


**Fig. S45 Overview of Neu5Ac linkage distribution during human neurodevelopment.** Distribution of α(2-3)- and α(2-6)-linked Neu5Ac on complex N-glycans of the **(A)** fetal brain and **(B)** adult brain by region. All possible Neu5Ac linkage combinations are depicted for mono-, di-, tri-, and tetrasialylated complex N-glycans. Error bars show the standard deviation. ND – not detected; # – detected as minor structures. All data are expressed as a percentage of total sialylated complex N-glycans. DFC – dorsolateral prefrontal cortex; HIP – hippocampus; STR - striatum; CBC - cerebellar cortex.

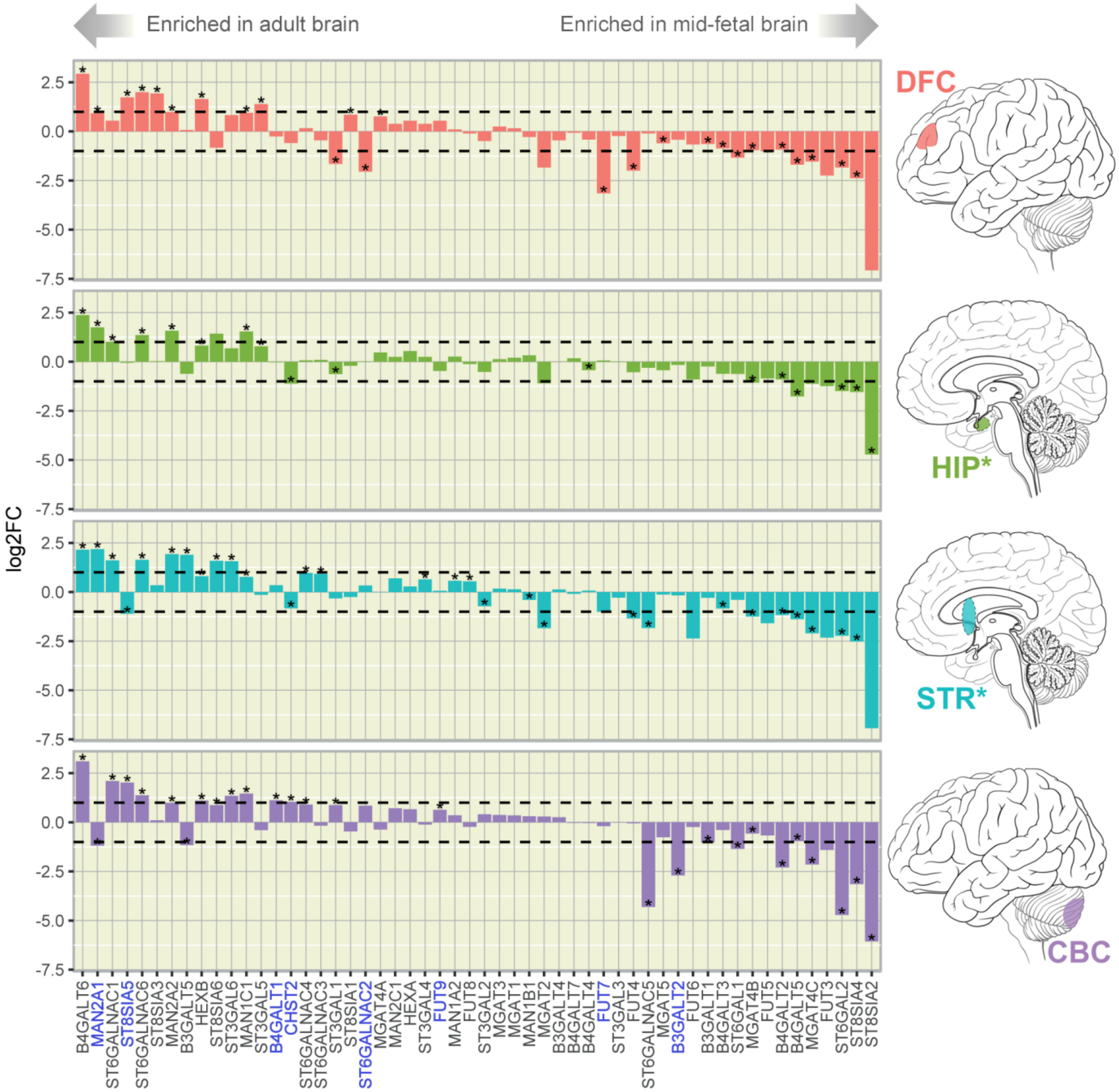


**Fig. S46. Temporal dynamics of glycogene expression during human neurodevelopment.** Each column corresponds to one glycogene with gene names displayed on the bottom. The y-axis shows the binary logarithm of the fold change in expression of glycogenes between the adult and fetal brain. Positive fold change indicates that expression of the glycogene is enriched in the adult brain, while negative fold change indicates that it is enriched in the fetal brain. Statistical significance was assessed using the Wilcoxon-Mann-Whitney test and a significant result is indicated by an asterisk. The brain region is indicated by the icon on the right.

Glycogenes whose expression is developmentally regulated in a region-specific manner are indicated in blue text. Icons depict adult brain regions; corresponding nascent structures were sampled in mid-fetal brains. DFC – dorsolateral prefrontal cortex; HIP – hippocampus; STR - striatum; CBC – cerebellar cortex. *Denotes that brain regions are internal and not visible from the view shown.

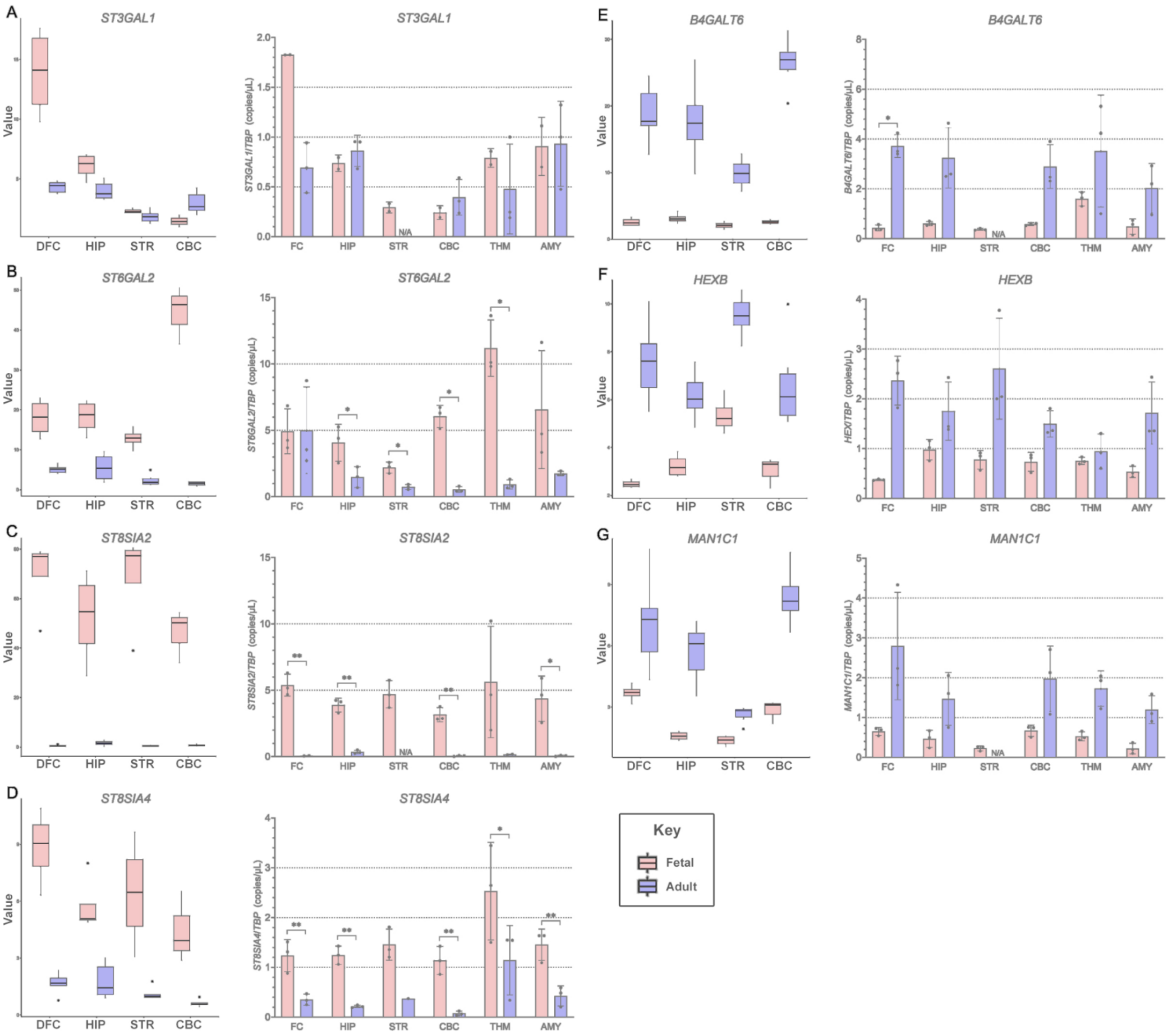


**Fig. S47. Developmentally-regulated expression of glycogenes in the human brain.** RNA-Seq data for glycogene expression in the fetal and adult human brain across six brain regions (left panel) was validated using ddPCR (right panel) for the following glycogenes: **(A)** *ST3GAL1*, **(B)** *ST6GAL2*, **(C)** *ST8SIA2*, **(D)** *ST8SIA4*, **(E)** *B4GALT6*, **(F)** *HEXB*, **(G)** *MAN1C1*. Multiple unpaired t-tests were used to assess statistically significant differences between groups. P-values were adjusted for multiple comparisons using the False Discovery Rate method. Statistically significant differences are labelled with asterisks; * 0.01<q-value<0.05, **q-value<0.01. DFC – dorsal frontal cortex; FC – frontal cortex; HIP – hippocampus;

STR - striatum; CBC – cerebellar cortex; THM – thalamus; AMY – amygdala; N/A – due to a limited amount of material, adult STR samples were lacking for some experiments.

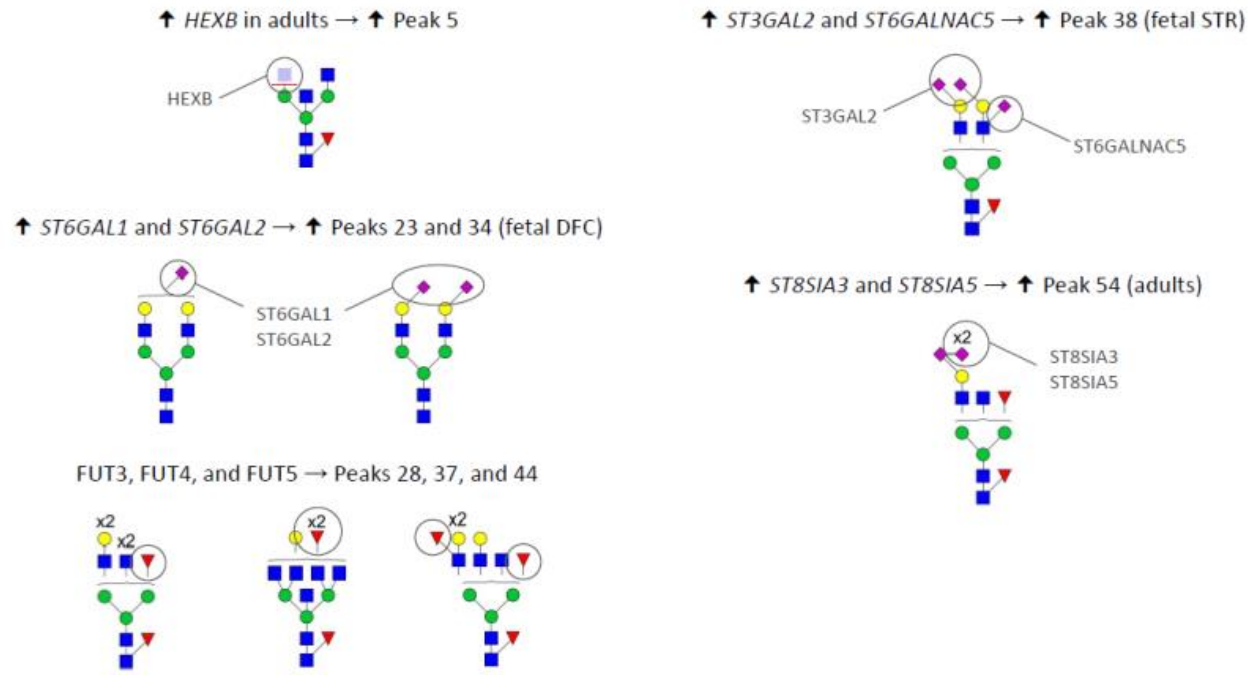


**Fig. S48. Summary of identified associations between the glycotranscriptome and the N-glycome.** Bioinformatically-deduced associations between the expression of individual glycoenzymes and the abundance of specific chromatographic peaks are displayed. Depicted are the proposed structures of the major N-glycans in each of the respective chromatographic peaks with the candidate glycoenzymes and the site of their enzymatic activity indicated.

**Supplementary Tables:**

Supplementary Tables are provided in separate Microsoft Excel files.

**Table S1. Postmortem brain tissue samples.**

Information regarding the brain tissue samples used in this study. PMI – postmortem interval.

**Table S2. Samples passing quality control.**

Information regarding the brain tissue samples that passed quality control and were used in the final analysis. dlPFC – dorsolateral prefrontal cortex; HIP – hippocampus; STR – striatum; CBC – cerebellar cortex; M – male; F – female.

**Table S3. N-glycan content of chromatographic peaks.**

The brain N-glycome was separated into 58 chromatographic peaks by HILIC-UPLC and masses of individual glycan structures were detected by MALDI-TOF/TOF-MS. Heatmaps show the cumulative relative abundance of N-glycans in a given chromatographic peak as determined by ion intensity and expressed as a percentage. Where MS data were not available, asterisks indicate N-glycans that were imputed to be the dominant N-glycan in that peak from equivalent peaks in other brain regions of the same species, or, in the case of CBC peaks, equivalent peaks in other species, for the purposes of calculating derived traits. In the human specimen table, N-glycans shaded in red are enriched in the fetal brain, while N-glycans shaded in blue are enriched in the adult brain. N-glycans comprising <10% of the cumulative relative abundance are not shown in the adult specimen table. Compositional notation: H – hexose; N – *N*-Acetylhexosamine; F - deoxyhexose (i.e. fucose); E – esterified Neu5Ac [i.e. α(2-6)-linked]; L – lactonized Neu5Ac [i.e. α(2-3)- or α(2-8)-linked]; S – sulfate group. Brain region: dlPFC – dorsolateral prefrontal cortex; HIP – hippocampus; STR – striatum; CBC – cerebellum.

**Table S4. Core brain N-glycans.**

A list of the 60 ubiquitous N-glycans that were detected in all brain regions of all species. H - hexose; N – *N*-Acetylhexosamine; F – deoxyhexose (i.e. fucose); E – esterified Neu5Ac [i.e. α(2-6)-linked]; L – lactonized Neu5Ac [i.e. α(2-3)- or α(2-8)-linked].

**Table S5. Intra-species regional differences in the abundance of individual N-glycan peaks.**

Statistical significance between brain regions was tested using ANCOVA, with age and sex as covariates, with correction for multiple testing using the Bonferroni procedure. P-values $< 0.05$ after correction for multiple testing are shaded green. GP – glycan peak; SE – standard error of estimated effect; dlPFC – dorsolateral prefrontal cortex; HIP – hippocampus; STR – striatum; CBC – cerebellum.

**Table S6. Inter-species differences in the abundance of individual N-glycan peaks across homologous brain regions.**

Statistical significance between species was tested using ANCOVA, with sex as a covariate, and correction for multiple testing was performed using the Bonferroni procedure. P-values $< 0.05$ after correction for multiple testing are shaded green. GP – glycan peak; SE – standard error of estimated effect; dlPFC – dorsolateral prefrontal cortex; HIP – hippocampus; STR – striatum; CBC – cerebellum.

**Table S7. Calculation of derived traits.**

Each derived trait was calculated as the summed area of all chromatographic peaks in which the dominant N-glycan displays that particular trait. Derived traits were calculated individually for each sample group (i.e. each specific N-glycome) due to the qualitative differences in peak content across brain regions and species. Peaks labelled with red text were imputed from equivalent peaks in other brain regions of the same species, or, in the case of CBC peaks, equivalent peaks in other species in cases where mass spectrometry data were unavailable. dlPFC – dorsolateral prefrontal cortex; HIP – hippocampus; STR – striatum; CBC – cerebellum.

**Table S8. Statistical analysis of N-glycan derived traits.**

Analyses of differences in N-glycan derived traits between sample groups were performed using the analysis of covariance (ANCOVA) model. When regional differences in N-glycosylation were analyzed within a single species, age and sex were included as additional covariates, while sex was included as a covariate when comparing N-glycomes across species. Correction for multiple testing was performed using the Bonferroni procedure. P-values < 0.05 after correction for multiple testing are shaded green. dlPFC – dorsolateral prefrontal cortex; HIP - hippocampus; STR – striatum; CBC – cerebellum.

**Table S9. Glycogene list.**

List of 47 genes whose expression was analyzed in this study. These glycogenes encode enzymes that are predominantly involved in N-glycan synthesis.

**Table S10. Regional differences in the abundance of individual N-glycan peaks during human brain development.**

Relative N-glycan peak abundance was compared between fetal and adult samples for each brain region (except the hippocampus due to an insufficient number of fetal samples). Positive fold change indicates peak abundance is greater in the adult brain, while negative fold change indicates that it is greater in the fetal brain. Statistical significance was assessed using the Wilcoxon-Mann-Whitney test. P-values < 0.05 are shaded green. No statistically significant differences were found after P-values were adjusted for multiple testing. dlPFC – dorsolateral prefrontal cortex; HIP – hippocampus; STR – striatum; CBC – cerebellum.
